# Comparative analysis of neuronal proteolytic pathways reveals neuron-specific and sub-compartmental-specific capacities with aging

**DOI:** 10.64898/2026.06.02.729733

**Authors:** Mira Sleiman, Dimitra Ranti, Sudarson Baskaran, Patricia Kreis, Saskia Muth, Christian Gallrein, Agnieszka Münster-Wandowski, Norman Rahnis, Emilio Cirri, Britta J. Eickholt, Janine Kirstein

**Affiliations:** Leibniz Institute on Aging – Fritz-Lipmann-Institute (FLI), Beutenbergstrasse 11 07745 Jena, Germany; Institute of Molecular Biology and Biochemistry, Charité Universitätsmedizin Berlin, 10117 Berlin, Germany; Friedrich-Schiller-Universität, Institute for Biochemistry & Biophysics, Hans Knöll-Strasse 2 07745 Jena, Germany; Institut für Integrative Neuroanatomie, Charité - Universitätsmedizin Berlin, 10117 Berlin, Germany

**Author notes:** co-corresponding authorship, Contact. contributed equally (Mira Sleiman, Dimitra Ranti, Sudarson Baskaran).

**Keywords:** protein turnover, synaptosome, neuron, aging, proteasome, autophagy

## Abstract

Proteostasis is essential for maintaining neuronal function, and its dysregulation is a hallmark of aging and neurodegeneration. The ubiquitin-proteasome system (UPS) and macroautophagy are the two major proteolytic pathways responsible for protein degradation. However, their capacity and regulation differ between cell types and across aging. To elucidate the activity of both proteolytic pathways with aging, we performed a comparative analysis of the activity of UPS and macroautophagy in distinct neuronal subcellular compartments, in the cytosol and at synaptic terminals, across aging in neurons of *Mus musculus* (mouse) and *Caenorhabditis elegans* (nematode). In mice, our results identified differences between brain areas. While the cortical proteasomal activity declined with aging in both the cytoplasmic as well as synaptic neuronal subcompartments, the cerebellar proteasomal activity decreased only in the cytoplasmic compartment with aging. In *C. elegans,* we detected a decrease of proteasomal activity in both cytoplasmic and synaptic compartments of neurons. Interestingly, we observed a dysregulation of macroautophagy in both neuronal subcompartments of the cortex and cerebellum in mice as well as in *C. elegans* neurons with aging. Thus, we uncovered neuron-specific and subcompartmental-specific proteolytic capacities with aging that could manifest in different neuronal vulnerabilities for proteotoxic challenges with aging.

## Introduction

Misfolded and damaged proteins that accumulate with aging must be removed to preserve cellular function throughout life ^1^. This is of particular importance in post-mitotic cells such as neurons. The ubiquitin proteasome system (UPS) is known to degrade regulatory proteins as well as dysfunctional and misfolded proteins ^2^. The UPS consists of an elaborate network of ubiquitin ligases and the 26S proteasome. Protein substrates are first marked with ubiquitin that can be elongated to form ubiquitin chains that are then recognized by the 19S component of the proteasome together with cofactors. The substrate protein is unfolded and translocated into the proteolytic 20S core component of the proteasome by the 19S component for subsequent degradation of the protein into smaller peptides ^3^. An alternative degradation route that also eliminates protein aggregates and even dysfunctional organelles is macroautophagy. Here, substrates are primed for degradation by first binding to autophagy adaptor proteins such as p62, which then associate with the lipidated form of microtubule-associated protein 1 light chain 3b (LC3B-II) of the forming autophagosome. The autophagosome then fuses with lysosomes to degrade the engulfed substrates ^4–6^. It has been demonstrated that suppression of basal autophagy can cause neurodegeneration ^6–8^. Thus, the elimination of misfolded and aggregated proteins by autophagy prevents an accumulation of damaged proteins and thereby preserves neuronal integrity.

Our understanding of the proteolytic capacity of neurons and at their distinct subcellular sites during aging, when protein damage occurs, is very limited ^9,10^. Proteomic analyses that mainly relied on rodent brain analyses, demonstrated that the half-life time of synaptic protein varies from days to weeks ^11–13^. Notably, neuronal activity correlated with the degradation rate of synaptic proteins. A suppression of neuronal activity led to an increase in protein lifetimes ^14^, suggesting that synaptic proteins need to be replaced regularly to maintain synaptic function. A clearance of proteins from the presynapse could be achieved by either transporting them to different subcellular sites e.g. the soma or by local proteolysis at the presynaptic site.

The components of the UPS including E3 ligases have been detected at the presynapse and several proteasome-specific presynaptic protein substrates have been identified, including Synphilin, CDCrel-1, Synaptophysin, DUNC13, RIM and Bassoon ^15–17^. Proteins related to autophagic and endolysosomal degradation such as AMBRA1, AP-2, LC3, GABARAPs and ATG proteins have also been detected at the presynaptic site (summarized in ^18^). The presence of components of both proteolytic systems argues for local synaptic protein degradation by the UPS and autophagy. While neuronal UPS and autophagy have been studied in various species with aging including differences between sexes and in the context of neurodegenerative diseases ^19–22^, much less is known about the synaptic proteolytic activity.

With this study, we set out to elucidate the contribution of the UPS and macroautophagy to protein degradation in neuronal sub-compartments and how their activity changes with the progression of aging. Moreover, we analyzed the two proteolytic mechanisms in two different brain areas of mice, the cerebral cortex and the cerebellum that differ substantially in their susceptibility to neurodegeneration ^23–25^. The cortex is responsible for cognitive functions and neuronal activity that decreases with aging due to e.g. synaptic loss, reduced neurotransmitter release, and increased oxidative stress that can lead to protein misfolding and aggregation ^26,27^. The cerebellum on the other hand is more resistant towards age-associated functional defects ^28,29^. We thus speculated that this difference could be connected to their proteolytic capacity and its change with aging. The murine analyses were complemented with studies in the nematode *Caenorhabditis elegans* to assess the proteolytic activity of the UPS and autophagy at the synapse and in the cytosol with the progression of aging in a short-lived animal. We used and established new methodologies to isolate and study the synaptic and cytoplasmic proteolytic pathways of two murine brain regions and *C. elegans* with the progression of aging of the animals. In mice, we observed a brain region- and subcompartmental-specific decline of proteasomal activity with aging. We could validate the age-associated reduction of proteasomal activity in *C. elegans* and observed a decline of the UPS activity in both, cytoplasmic and synaptosomal compartments. Macroautophagy is also negatively affected by aging in mice and in nematodes at both subcompartments, suggesting that in neurons both major proteolytic pathways are compromised with aging and that the lack of clearance capacity of damaged proteins could contribute to the vulnerability of specific brain regions and neuronal subcompartments to age-associated neurodegeneration.

## Results

### Establishing and adapting a protocol for the isolation of murine and nematode synaptosomes

To assess synaptic UPS activity and macroautophagy, we isolated crude synaptosome-enriched fractions (for clarity referred to as synaptosomes throughout the manuscript) from the cerebral cortex and cerebellum of murine brains, as well as from *C. elegans* neurons (Fig. 1A). Murine synaptosomes were prepared using biochemical differential centrifugation, which separates cellular components based on molecular weight and detergent solubility of the lysate (^30^; Fig. 1A 1)). The enrichment in synaptosomes was confirmed by western blot analysis, using synaptophysin and PSD95 as a pre- and post-synaptosomal marker, respectively (P2 fraction) and GAPDH as marker that is enriched in the cytoplasm (S2 fraction) (Fig. 1B). Additionally, electron microscopy (EM) analysis validated the integrity of the synaptosomal preparations (Fig. 1C).

**Figure 1.**
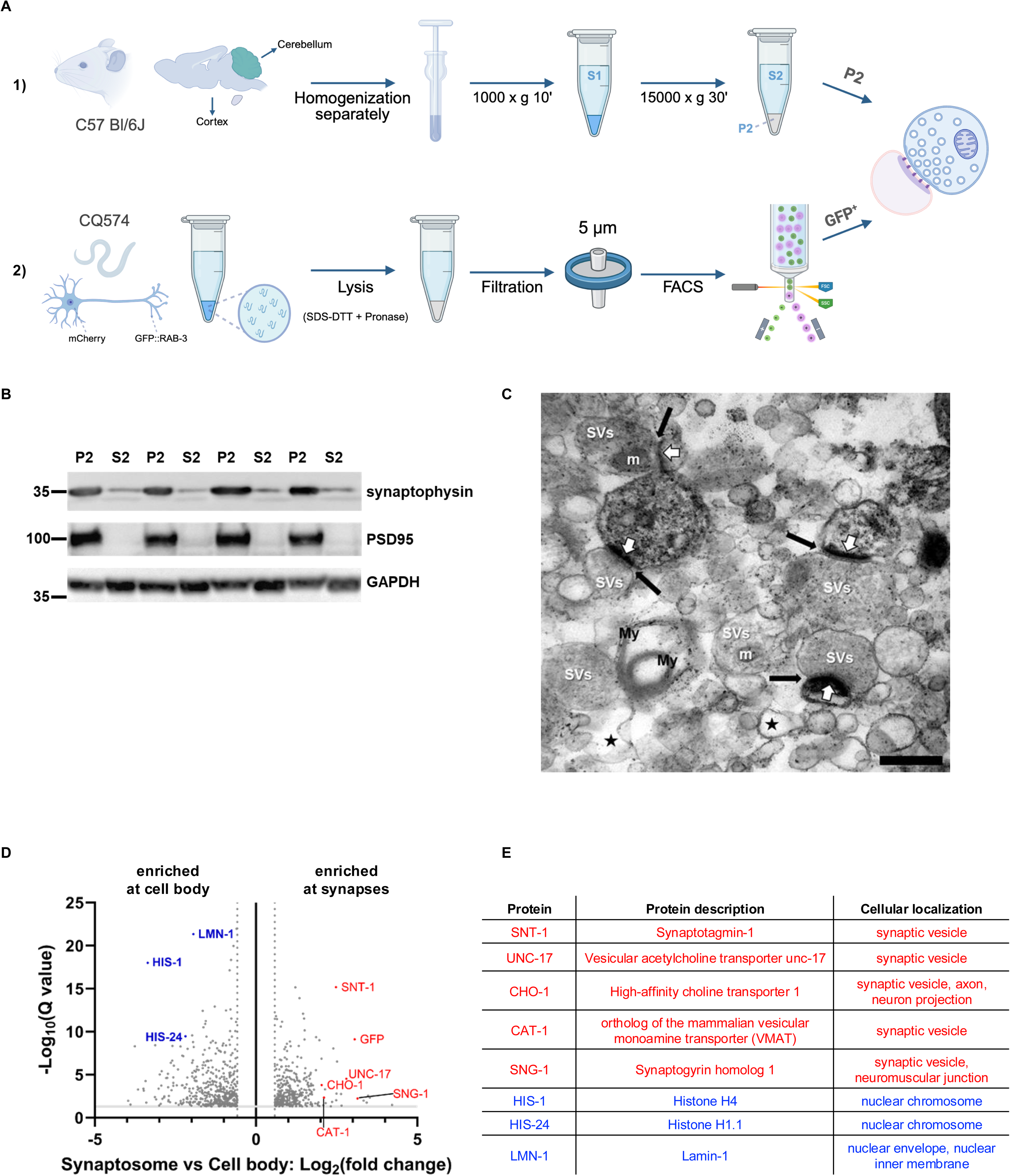
Establishing and adapting a protocol for the isolation of murine and nematode synaptosomes. **A)** Workflow of the isolation of crude synaptosomes of mice (1) and *C. elegans* (2). For the biochemical sedimentation-based fractionation of the murine samples, we used the mouse line C57BI/6J and for the FACS-mediated fractionation of the nematode samples, we used the *C. elegans* strain CQ574 that expresses mCherry and GFP::RAB-3 under the control of the *rab-3* promoter. **B)** Validation of the preparation of the S2 (cytoplasmic fraction) and P2 (crude synaptosomes) fractions by western blot. The samples were obtained from four different murine brains and analyzed for established pre- and post synaptosomal markers, synaptophysin and PSD95, respectively as well as GAPDH as a cytoplasmic marker. **C)** Electron micrograph of crude murine synaptosomes. The image shows a uniform distribution of vesicle-filled axonal profiles (SVs), intact synapses with adjacent spines showing the post-synaptic density characteristic of asymmetric / excitatory synapses (white arrows), and synaptic clefts (black arrows). Note that the crude synaptosome fraction is slightly contaminated with empty membrane fragments, mostly outer synaptosomes (asterisks) and myelin (My). No large structures such as cell bodies or nuclei were present. Mitochondria (m) observed in axonal terminals confirm the integrity of the isolated synaptosomes. Scale bar is 500 nm. **D)** Volcano plot depicting the proteomic comparison of FACS-isolated synaptic (GFP-positive) versus somatic (mCherry-positive) compartments of *C. elegans*, pooled from three independent biological experiments (N = 3; ∼6000 nematodes per experiment), validating the compartment-specific enrichment of our method. Proteins that are enriched at the synapses are on the right side whereas on the left are proteins that are enriched at the cell body. Proteins were considered significantly enriched or depleted if they exhibited a fold change greater than 1.5 (or below -1.5) and a false discovery rate (FDR) below 0.05 (Benjamini-Hochberg correction on moderated t-tests). Highlighted in red and blue are proteins known for their sub-cellular localization at synapses and cell body, respectively. **E)** The table lists the identified proteins highlighted in the volcano plot shown in (D) using the same color code. Proteins that were enriched in the synapses are shown in red and proteins that were enriched in the cell body are depicted in blue.

For *C. elegans*, we adapted a fluorescence-activated cell sorting (FACS)-based protocol to isolate fluorescently labeled synaptic and cytoplasmic compartments (^31^; Fig. 1A 2)). The CQ574 strain expresses the pre-synaptic protein RAB-3 fused to GFP that localizes at the synapse, while mCherry is targeted to the cytoplasmic compartment including the soma and neurites. Day 4 old nematodes (day 1 of adulthood) were gently lysed in a detergent-containing buffer and treated with Pronase to digest the cuticle. The resulting cell suspension was then subjected to FACS to separate GFP-positive synaptic compartments from mCherry-positive cytoplasmic fractions. In line with the published protocol by the lab of Coleen Murphy ^31^ that isolated RAB-3::GFP-positive presynaptic regions by FACS and referred to them as presynaptic/synaptic compartments rather than classical synaptosomes, we also refer to the GFP-positive material as presynaptic/synaptic compartments and to the mCherry-positive material as soma-enriched neuronal compartments. The mCherry-positive fraction contains proteins associated with somatic and dendritic structures and general neuronal signaling, including AAK-2, AKT-1, ASNA-1, MEC-7, MEC-12, MEC-17, MEC-6, CHDP-1, and PPM-2, which are linked to neuronal soma, mechanosensory microtubule architecture, and dendritic development ^32–37^. In addition, this fraction is enriched for major cytoplasmic and nuclear components such as histones (HIS-1, HIS-24) and the nuclear lamina protein LMN-1, further supporting its soma-enriched identity. By contrast, the GFP-positive fraction is enriched for canonical presynaptic and vesicle-trafficking proteins such as SNT-1/4, SNG-1, UNC-17, UNC-31/CAPS, CAT-1, CHO-1, GLT-1, CASY-1, UNC-104 and UNC-32, all with established roles at presynaptic terminals and synaptic vesicles ^38–42^. Together, this distribution of markers, paralleling the presynaptic enrichment and validation strategy of Arey et al. ^31^, supports our interpretation that the GFP-sorted material represents presynaptic/synaptic compartments, whereas the mCherry-sorted material represents soma- and dendrite-enriched neuronal cytosol (Fig. 1D+E).

### Proteomic profiling of synaptosomes with aging

Using these established isolation methods, we conducted a proteomic content analysis of synaptosomal enriched fractions (P2 from murine samples and FACS-isolated synapses from *C. elegans*) at different time points throughout aging. We chose three different ages, 2 months for young adults, 8 months for mature adults and 18 months for old mice in which locomotor activity and cognition were impaired ^43^. We worked with male mice and are aware that there could be sex-specific differences in the composition and capacity of proteases in the synaptosomal enriched fractions as recently reported for chaperone-mediated autophagy ^22^. Murine synaptosome samples from three age groups (2, 8, and 18 months) contained over 5000 protein groups (Uniprot classification) in each cohort (Fig. S1A). In contrast, the FACS-sorted *C. elegans* fractions (synaptic, cytoplasmic and whole neurons) exhibited a lower proteomic complexity, with up to 2000 quantified protein groups (Fig. S1B+C). This difference aligns with the more complex proteome of the murine system ^44,45^. For *C. elegans* we chose to study day 4 old animals that are referred to as young adults and reached maturity that marks the onset of aging and day 7 old animals that are at the end of their fertile period and are considered as older animals exhibiting many hallmarks of aging ^46–48^. Comparative proteomic analysis of synaptic samples from *C. elegans* at day 4 of life (young adults) and day 7 of life (older animals) identified 2085 proteins common to both time points (Table 1, top half and Table S1). Of these, 145 were significantly enriched (P < 0.01, fold change > 2), whereas 54 were significantly depleted in day 4 old nematodes. In contrast, the corresponding comparison of cytoplasmic samples revealed 1552 shared proteins, with 55 enriched and 24 depleted in day 4 old nematodes. A comparative proteomic analysis of synaptic samples of day 4 and even older animals (day 10 of life) identified 736 proteins common to both ages with 138 enriched and 109 depleted proteins in day 4 old nematodes compared to day 10 old nematodes. The same number of proteins (736) were identified for the cytoplasmic samples of both ages that showed an enrichment of 84 proteins and a depletion of 158 proteins in young versus day 10 old animals. These results show a dynamic remodeling of both the synaptic as well as cytoplasmic proteome with aging. Because the *C. elegans* synaptic datasets contained a lower total protein input, applying the same FDR < 0.05 threshold used for the murine samples yielded only a small number of significant proteins. We therefore used a more permissive criterion (p < 0.01, fold-change >2) to define age-regulated candidates in nematodes and concentrated our interpretation on consistent functional enrichments rather than on single targets.

**Table 1:** Number of differentially regulated proteins of the *C. elegans* samples. Shown are the comparisons of day 4 and day 7 old nematodes (upper rows) and day 4 vs day 10 old nematodes (lower rows) for the synaptic and somatic fractions, respectively. The samples are then further split into the number of total proteins that refer to proteins common to both ages and the number of enriched and depleted proteins of day 4 vs day 7 and day 4 vs day 10 old animals, respectively.

| Condition | Sample | Proteins Identified | Number of proteins |
| --- | --- | --- | --- |
| *C. elegans*<br>day 4 vs day 7 | Synaptic | Total | 2085 |
|  |  | Enriched | 145 |
|  |  | Depleted | 54 |
|  | Somatic | Total | 1552 |
|  |  | Enriched | 55 |
|  |  | Depleted | 24 |
| *C. elegans*<br>day 4 vs day 10 | Synaptic | Total | 736 |
|  |  | Enriched | 138 |
|  |  | Depleted | 106 |
|  | Somatic | Total | 736 |
|  |  | Enriched | 84 |
|  |  | Depleted | 158 |

Proteomic comparison of synaptosomes isolated from the cerebral cortex of 2 months old and 18 months old mice identified 5836 proteins shared between the two age groups. Among these, 118 were significantly (Q < 0.05, fold change > 1.5) enriched and 171 were significantly depleted in synaptosomes of 2 months old mice (Table 2). In comparison with the *C. elegans* synaptic aging dataset, these findings indicate that the mouse cortical synaptosome proteome changes less with aging, suggesting species-specific differences in synaptic proteome landscape during aging. While we cannot exclude that methodological factors (crude synaptosome preparation for mice vs FACS sorting for nematodes) may affect these differences, the overall trend is consistent with known differences in organismal complexity and has been observed in previous comparative proteomic studies ^49^.

**Table 2.** Number of differentially regulated synaptic proteins of the murine samples of 2 month old vs 18 month old animals. Proteins that were identified in the synaptosomes of both ages are listed as total and enriched and depleted proteins refer to the comparison between 2 month vs 18 month-old mice.

| Condition | Sample | Proteins Identified | Number of proteins |
| --- | --- | --- | --- |
| Mice<br>2 months old vs<br>18 months old | Synaptosomes<br>from cortex | Total | 5836 |
|  |  | Enriched | 118 |
|  |  | Depleted | 171 |

To obtain more insights on the physiological change in synapses with aging, Gene Ontology (GO) analyses were performed of the synaptic proteome from 2 months old vs 18 months old, 2 months old vs 8 months old and 8 months old vs 18 months old mice (Fig. S2 + S3) and from *C. elegans* young adults (day 4 of life) versus older animals (day 7 of life) and old animals (day 10 of life) (Figs. S4). GO analysis of mouse cortical synaptosomes from 2 months and 18 months old animals revealed an enrichment for proteins involved in protein synthesis, regulation of apoptosis and in nucleotide and lipid metabolism in synaptosomes of young 2 months old mice (Fig. S2A). The observed enriched lipid and nucleotide metabolic pathways are also enriched in synaptosomes of 2 months old mice in comparison with 8 months old mice (Fig S2C). The GO term comparisons of 2 months old mice vs 18 months old and also 8 months old mice show a depletion of components of the synthesis and assembly of the mitochondrial respiratory chain and metabolic and here mainly catabolic mitochondrial pathways as well as lipid metabolic categories, suggesting changes in the energy and mitochondrial metabolism in the aging synapse (Fig. S2B+D). The comparison of the synaptic proteome between 8 months old and 18 months old mice did not reveal any significantly depleted pathways and showed only a limited number of enriched pathways in the 8 months old mice including protein synthesis and the regulation of apoptosis that are also enriched in 2 months old animals compared to 18 months old mice (Fig. S2A), suggesting that the synaptic proteome undergoes only minor pathway changes between 8 months and 18 months and most changes occur early in life between 2 and 8 months of age.

For *C. elegans*, we used WormEnrichr to compare the data sets of enriched and depleted proteins across 10–15 annotation databases ^50,51^. We analyzed the cytoplasmic and synaptic fractions of day 4, day 7 and day 10 old nematodes. With aging, we observed a depletion of proteases in both subcompartments that we describe below in more detail. We however also observed that a high number of ribosomal proteins are significantly depleted in day 10 old animals in the cytoplasmic (50 ribosomal proteins) and synaptic (15 ribosomal proteins) compartments (repository; see data availability). This severe reduction of ribosomal proteins with aging that we also observed in the murine synaptosomal proteome data validates the documented decline in protein biosynthesis with aging ^48^. The observed reduction of splicing components (SNR-1/2/6, RSP-2/6 and TEG-4) in the neuronal cytoplasmic compartment indicates that RNA processing is already impaired in day 10 old nematodes. However, there are also enriched proteins in older nematodes. Lysosomal proteins such as ARL-8 and ASP-4 that were depleted on day 7 are enriched in day 10 old animals and could point to an accumulation of lysosomal components upon its dysfunction with aging (see below, Fig. 2B). In addition, mitochondrial catabolic enzymes (e.g. ICL-1, ACDH-7/10 and CPT-1) are specifically enriched in the synapse of day 10 old animals, suggesting an altered mitochondrial metabolism in the aging synapse (repository; see data availability).

**Figure 2.**
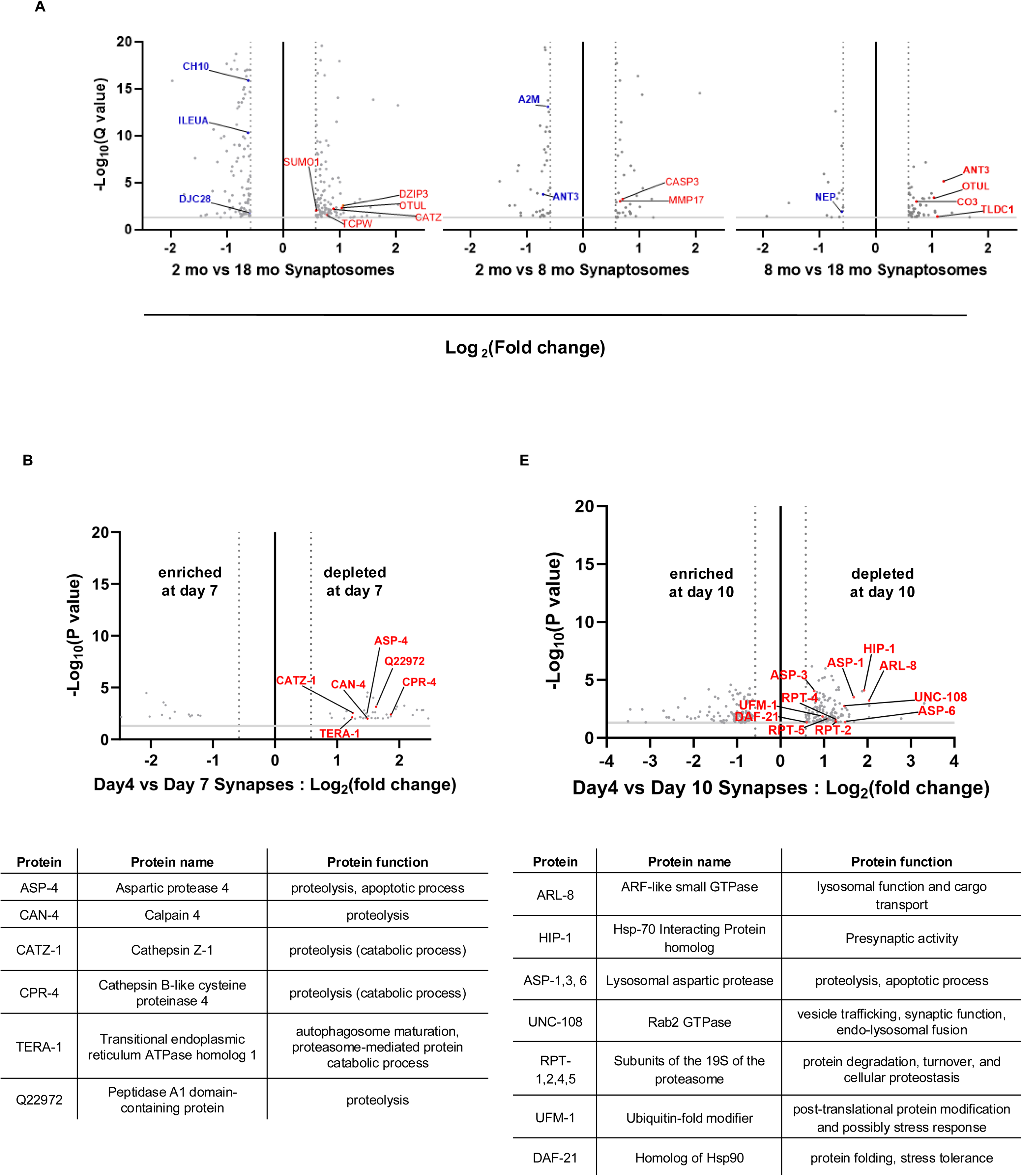
Analysis of the proteome data from murine and nematode subcompartments. **A)** Volcano plots depicting the comparison of proteins of murine cortical synaptosomes of 2 vs 18 months old (left), 2 vs 8 months old (middle) and 8 vs 18 months old (right). Proteins were considered significantly enriched or depleted if they exhibited a fold change greater than 1.5 (or below -1.5) and a false discovery rate (FDR) below 0.05 (Benjamini-Hochberg correction on moderated t-tests). Points on the right side of each graph represent proteins significantly enriched in 2 months relative to 18 months old mice (left plot), in 2 months relative to 8 months old mice (middle plot) and 8 months relative to 18 months old mice (right plot). Points on the left indicate depleted proteins for the same comparative volcano plots. Highlighted in red (enriched in the younger cohort) and blue (enriched in the older cohort) are proteostasis components such as proteases and regulators of the proteasome and chaperones. **B)** Volcano plots depicting the proteomic comparison of synaptic compartments isolated from *C. elegans* from three independent biological experiments (N = 3; ∼6000 nematodes per experiment) each at 4 days of life versus 7 days old animals (left) and between 4 days of life versus 10 days old animals (right). Proteins were considered significantly enriched or depleted if they exhibited a fold change greater than 2 (or below -2) and a p value below 0.01. Points on the right side represent proteins significantly enriched in day 4 samples relative to the older cohort, whereas points on the left indicate proteins enriched in older animals relative to day 4. Highlighted in red are proteostasis components such as proteases that are enriched in the synaptosomes of young (day 4 old) nematodes. The tables list these enriched proteases highlighted in red in the volcano plots above with the respective protein name and function.

A subsequent analysis of specifically the synaptic proteome revealed that proteins enriched in synaptosomes of day 4 old nematodes compared to day 7 old animals showed strong associations with proteolysis (Fig. S4). To obtain finer spatial resolution at the synaptic level, we performed a GO analysis using the SynGO database (Fig. S5A+B) ^52^. The analysis of the nematode data confirmed the WormEnrichr findings, showing that synapses of day 4 old animals were enriched in proteins associated with presynaptic, endocytic processes, transport and signaling. Together, these findings suggest that aging-related depletion of endo-lysosomal regulators could compromise synaptic autophagy with aging. The analogous comparative analyses between synaptosomes of day 4 and day 10 old nematodes, revealed an enrichment of proteasome components and other proteases, RNA-binding proteins and protein synthesis in synaptosomes of young day 4 old animals (Figs. S4C, S5C+D).

Next, we specifically focused on proteases and potential regulators of the proteolytic pathways in the synaptosomal fraction at different time points throughout aging for the murine and *C. elegans* synaptosomes and depicted the data in volcano plots (Fig. 2). For the murine samples, we observed an enrichment of proteases e.g. cathepsin Z (CATZ), components or regulators of the proteasome e.g. the deubiquitinase Otulin (OTUL), the E3 Ubiquitin ligase DZIP3, SUMO1 and a component of the chaperone complex TRIC (TCPW) in the synaptosomes of young (2 months old) mice whereas other chaperones such as the J-domain protein DJC28 and HSP10 (CH10) and a protease inhibitor (ILEUA) are enriched in 18 months old mice (Fig. 2A). These data indicate that the abundance of proteases and regulators of the proteolytic pathways as well as molecular chaperones change with aging in murine synaptosomes. The differences were as expected most pronounced between 2 months and 18 months (Fig. 2A, left) and fewer changes were observed for the comparisons of 2 months vs 8 months (Fig. 2A, middle) and 8 months vs 18 months (Fig. 2A, right). For *C. elegans*, we compared the synaptic proteome of young adult animals (day 4 of life) with that of older animals that are at the end of their fertile period (day 7 of life) and old animals (day 10 of life) (Fig. 2B). Analogous to the murine data, we observed an enrichment of lysosomal degradative enzymes e.g. cathepsins (CATZ-11, CPR-14), aspartic proteases (ASP-4, Q22972), Calpain (CAN-4) and the molecular chaperone p97 (TERA-1) in the synaptosomes of young animals compared to day 7 (Fig. 2B, left) and an enrichment of proteasome components (RPT-1, RPT-2, RPT-4 and RPT-5), proteases (ASP-1, ASP-3, ASP-6), ubiquitin-like proteins (UFM-1), chaperones and chaperone-binding proteins (HSP90 homolog, HIP-1) and GTPases involved in vesicular trafficking (UNC-108, ARL-8) in the synaptosomes of young animals compared to day 10 old nematodes (Fig. 2B, right). Combined, our synaptosomal proteome studies showed a decline in the abundance of proteases and changes of regulators of the proteolytic pathways with aging in mice and nematodes, suggesting a reduced synaptic proteolytic capacity with aging.

### Synaptic proteasomal activity declines with aging

The reduced abundance of proteases in the synaptosomes of both mice and nematodes (Fig. 2) suggest a reduced proteolytic activity with aging. We set out to assess the activity of the UPS and macroautophagy of the synaptic and cytoplasmic fraction in both animal models at different ages (Figs. 3-6). First, we performed proteasomal activity analyses using an ex vivo approach with the murine fractions of the cortex and the cerebellum (Fig. 3). We analyzed the cytoplasmic S2 fraction that is composed of soma and neurites and the synaptosome-enriched P2 fraction of mice of different ages: 2 months (young adult), 8 months (mature adult) and 18 months (aged mice). The protein lysates of both fractions were incubated with the fluorogenic 20S substrate Suc-LLVY-AMC. The chymotrypsin-like activity of the proteasome can cleave and then release the unquenched AMC fluorophore. The increase in AMC fluorescence over time thus reports on the activity of the proteasome. We observed a decreased activity of the proteasome of 18 months old mice compared to earlier time points (2 and 8 months) for the synaptosomal compartment of the cortex (Fig. 3A 3). The reduction of the proteasomal activity could be due to a reduced abundance of the proteasome or due to a decrease in the activity of the available pool of proteasomes. We thus quantified the proteasome levels and observed a decrease in the abundance of the 20S proteasome subunits (Figs. 3A 4) + S5). These data show that the age-dependent reduction of the proteasomal activity in the synaptic compartment is based on a reduced proteasome abundance with aging. No significant activity changes of the proteasome were observed for the cytoplasmic compartment, despite a reduced abundance of the 20S subunits with aging (Figs. 3A 1)+2) + S6). For the cerebellum, we detected a reduced proteasomal activity and abundance for the cytoplasmic fraction (Fig. 3B 1)+2)) and no significant change of the proteasomal activity for the synaptosomal compartment with aging (Fig. 3B 3)+4)).

**Figure 3.**
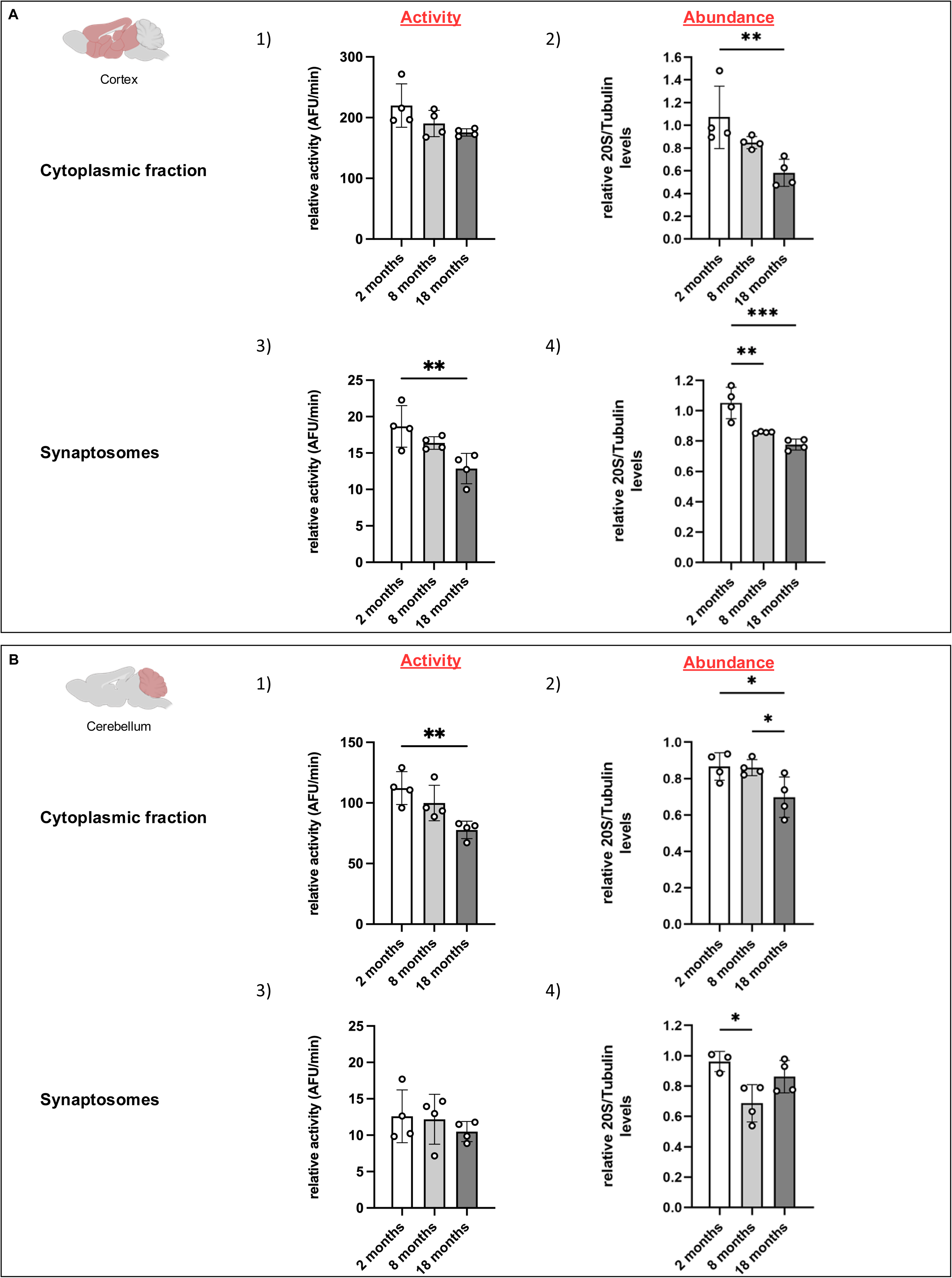
Analysis of the proteasomal activity of murine cortical and cerebellar neuronal sub-compartments with aging. **A)** Chymotrypsin-like activity (1) and abundance (2) of 20S ɑ-subunit of cortical brain samples from 2, 8 and 18 months old mice. The scatter dot plots depict the average chymotrypsin-like activity (1 and 3) in cytoplasmic (upper row) and synaptosomal (bottom row) fractions of the cortex, respectively. Every dot represents the average of technical duplicates of one animal of the curve’s slope normalized to that of its respective control. The abundance of the 20S proteasome subunits was assessed by western blot analysis of the same samples as used in (1 and 3). The plot depicts the relative 20S levels normalized to tubulin. **B)** As in (A) but for cerebellar fractions. In total, fractions from 4 different mice of each age were used. Significance was assessed by a one-way ordinary ANOVA and Tukey‘s test post hoc (* = p < 0.05; ** = p < 0.01; *** = p < 0.001).

In sum, we observed an age-associated decline in proteasomal activity in different neuronal subcompartments of the murine cortex and cerebellum.

### Reduced proteasomal activity at *C. elegans* soma and synapses with aging

Next, we wanted to assess the proteasomal activity of the nematode neuronal subcompartments. Although we could isolate neurons and synaptosomes from *C. elegans* for proteome analysis, the protein concentration was not sufficiently high for biochemical assays such as the AMC proteasome assay used for murine samples (Fig. 3). We thus used an in vivo genetic approach and generated a new *C. elegans* model where we targeted a UPS reporter ^53^ to distinct subcellular neuronal sites. For that, we used the promoter of the gene synaptogyrin (*sng-1*, Fig. 4A). The UPS reporter is based on a non-cleavable ubiquitin variant (Ub^G76V^) that is fused to the photoconvertible fluorophore Dendra2. The G76V mutation of ubiquitin prevents a deubiquitination of Dendra2 that is hence destined for degradation by the proteasome ^54,55^. Exposure to 405 nm induces a break in the backbone of Dendra2 that irreversibly converts the excitation and emission spectra of the fluorescence protein from 490/507 nm (ex/em) to 553/573 nm (ex/em). The photoconversion allows to monitor the then red-shifted abundance of Ub^G76V^-Dendra2 that can only be regulated by the activity of the proteasome. Any newly synthesized Ub^G76V^-Dendra2 will have the fluorescence properties of the non-converted Dendra2 (490/507 nm) reflected by “green” fluorescence. The new transgenic line expressing the UPS sensor allowed us to assess the activity of the proteasome in the living nematode at different ages. We analyzed day 3 old (L4 larvae), day 4 old (young adults) and day 7 old (older animals) nematodes. The UPS sensor is expressed in all neurons and localizes at different subcellular structures of the neurons including synaptic boutons (Fig. 4A; white arrows). We validated the synaptic localization of the new UPS sensor using the established synaptic marker RAB-3 ^56^ that is fused to mCherry and observed co-localization at synaptic boutons (Fig. 4B). In addition to the synaptic sites, Ub^G76V^-Dendra2 also localizes to the soma and neurites (cytoplasmic subcompartment) that hence allowed us to analyze the activity of the proteasome with subcompartmental resolution. Photoconversion was performed in the soma as well as at synaptic sites by a short exposure of the selected region of interest (ROI) to 405 nm. Ub^G76V^-Dendra2 fluorescence at 563 nm was then quantified immediately after photoconversion and 1 hour later (Fig. 4C). The difference in fluorescence intensity reports on the activity of the proteasome. The abundance of ATP does not change significantly in the analyzed time window and should thus not be a limiting factor for the activity of the proteasome^57^. For the soma, we observed that about 35% of the reporter was degraded within 1 hour in day 3 and day 4 old animals, whereas almost no degradation occurred anymore in day 7 old animals (Fig. 4D). Thus, we did not analyze later time points. Interestingly, we detected a higher proteasomal activity on days 3 and 4 at the presynaptic site compared to the soma as reflected by the lower level of remaining photo-converted Ub^G76V^-Dendra2, suggesting a higher proteasomal activity in synapses in young animals. With further aging, proteasomal activity ceased as well and day 7 old animals did not show any proteasomal activity anymore regardless of the neuronal subcompartment (Fig. 4D).

**Figure 4.**
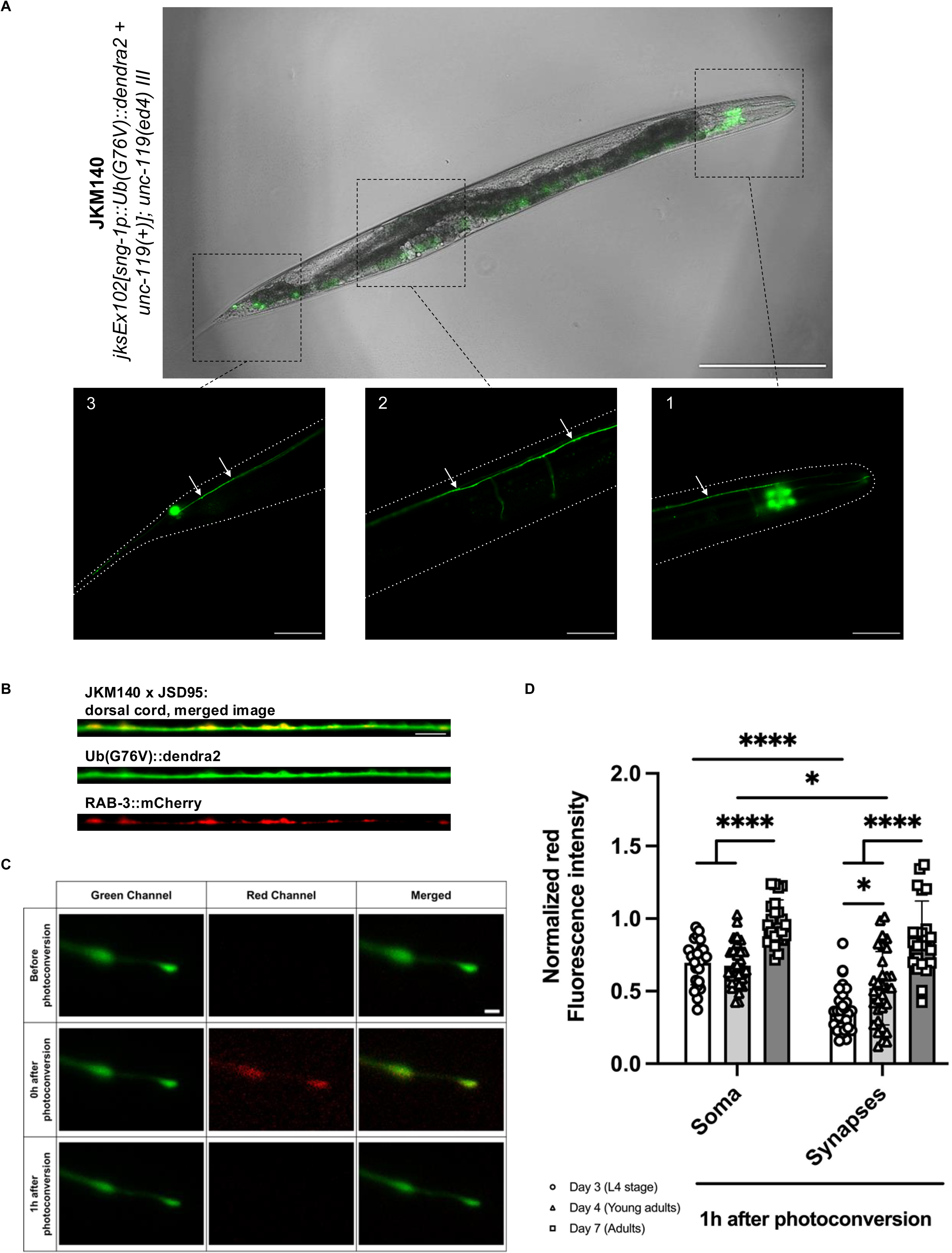
Analysis of somatic and synaptosomal UPS activity in *C. elegans* with aging. **A)** Representative microscopy images of the new UPS reporter strain, JKM140 (*jksEx102[sng-1p::Ub(G76V)::dendra2 + unc-119(+)]; unc-119(ed4) III*). A merge of the brightfield and GFP channel is shown on top and close-ups of three regions depicting the neuronal expression in the tail (1; on the right), the mid body (2; middle) and the head (3; on the left) are shown below. Arrows highlight synaptic boutons. Dashed white lines highlight the outline of the nematodes. Scale bars are 200 µm for the brightfield image and 50 µm for the insets. **B)** Validation of the synaptic localization of the UPS reporter of (A) with the RAB-3::mCherry expressing strain JSD95. Depicted is a representative image of the dorsal nerve cord of an animal expressing both the UPS reporter, *sng-1*::Ub(G76V)::Dendra2 and the synaptic marker RAB-3::mCherry. The merge shows, in yellow, the co-localization of Ub(G76V)::Dendra2 with RAB-3::mCherry at the synapses (GFP- and mCherry-positive). Scale bar is 5 µm. **C)** Exemplary photoconversion experiment of the Ub^G76V^::Dendra2 UPS reporter. Depicted are the individual and merged channels of synaptic boutons of the dorsal nerve cord at the tail region before (top row), immediately after (middle row), and 1 hour after photoconversion (bottom row). Scale bar is 1 µm. **D)** Relative red (561 nm) fluorescence of the photoconverted Ub^G76V^::Dendra2 as a readout for the UPS activity of day 3, 4 and 7 old nematodes. The scatter plot illustrates the average remaining red fluorescence 1 h after photoconversion at the soma (left) and presynapse (right) of the tail region. Every dot represents the relative remaining fluorescence intensity from one animal inversely correlated with the UPS activity. Photoconversion experiments were performed in triplicates (n = 23-31). Significance was assessed by an ordinary one-way ANOVA with Tukey’s multiple comparison test (p < 0.05 (*), p < 0.01 (**), p < 0.001 (***) and p < 0.0001 (****)).

We conclude that neuronal proteasomal activity is higher at synapses compared to the soma and rapidly declines at both sites with the progression of aging. Thus, the age-associated decrease of neuronal proteasome activity is conserved between nematodes and mice.

### Dysregulation of neuronal autophagy in aging mice

Next, we set out to analyze autophagy and started with the murine samples containing the cytoplasmic and synaptosomal fraction of the cortex and cerebellum. We used the same ages as for the proteasomal analysis (Fig. 3) and quantified the protein abundance of autophagy markers in the cytoplasmic and synaptosomal fraction of the murine cortex (Fig. 5A-D) and cerebellum (Fig. 5E-H) ^58^. For that, we performed western blot analyses of p62, LC3B, ATG9A, LAMP1 and ATP6V1A (Fig. 5). The selected markers cover distinct steps of the autophagy–lysosomal pathway, allowing for a comprehensive yet steady state assessment of autophagosomal and lysosomal markers ^4,58^. Specifically, ATG9A is one of the earliest proteins recruited to the phagophore assembly site. It functions as a lipid scramblase and is the only transmembrane protein among the core autophagy components. LC3B, a ubiquitin-like protein, is used to monitor autophagosome maturation. p62 is an autophagy adaptor protein that binds ubiquitinated cargo and delivers it to the expanding autophagosome. Importantly, it is also a substrate of autophagy. Together, LC3B and p62 provide complementary insights into autophagic flux and cargo load. Since macroautophagy relies on lysosomal degradation, we also assessed two markers of lysosomal function: LAMP1 (Lysosomal-Associated Membrane Protein 1), a major component of the lysosomal membrane, and ATP6V1A, a catalytic subunit of the vacuolar-type H⁺-ATPase (V-ATPase) complex, which is essential for lysosomal acidification.

**Figure 5.**
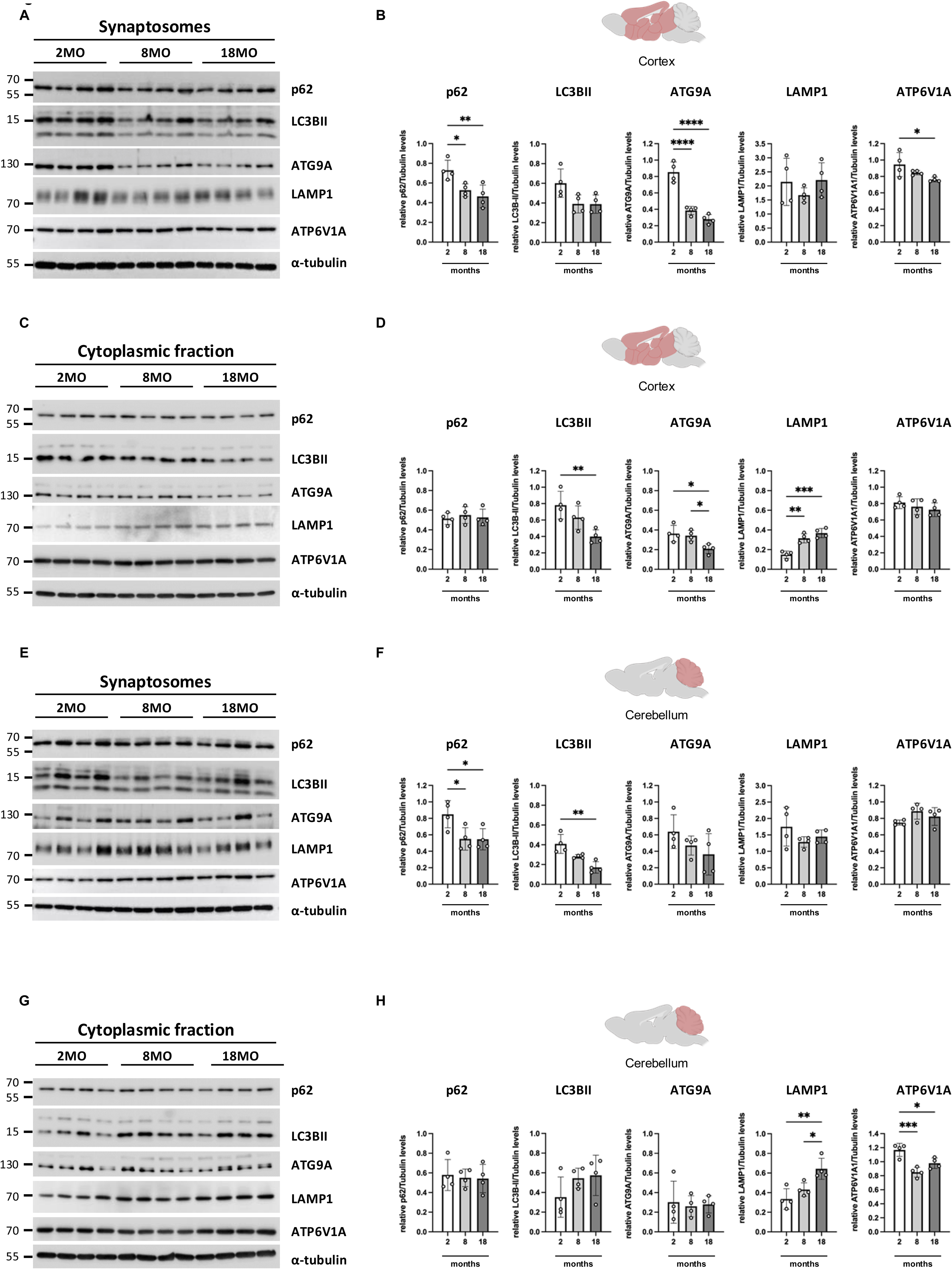
Analysis of murine somatic and synaptosomal autophagy with aging. **A)** Western Blot analysis of autophagy markers (p62, LC3BII, ATG9A, LAMP1, and ATP6V1A1) of synaptosomes (P2 pellet) extracted from the cortex of four different mice at different ages (2 months, 8 months and 18 months). α-tubulin was used as loading control. The two LC3 bands correspond to LC3BI (upper band) and the lipidated LC3BII (lower band). Molecular weight protein ladder is in kilodaltons. **B)** Quantitative analysis of (A). Depicted are the signal intensities of the autophagy-lysosomal markers p62, LC3BII, ATG9A, LAMP1, ATP6V1A relative to α-tubulin. For each animal, two technical replicates were averaged Mean ±SEM. One-way ANOVA Bonferroni test.*p<0.05, **p < 0.01, ****p<0.0001. **C)** Western Blot analysis of autophagy markers of the cytoplasmic fraction (S2) from the cortex as described in (A). **D)** Quantitative analysis of cytoplasmic fraction derived from cortex as described in (B). One-way ANOVA Bonferroni test. *p<0.05, **p < 0.01, ***p<0.001. **E)** Western Blot analysis as in (A), but of synaptosomes (P2) extracted from the cerebellum. **F)** Quantitative analysis of (E) as described in (B). One-way ANOVA Bonferroni test. *p<0.05, **p < 0.01. **G)** Western Blot analysis as in (C) but of cytoplasmic fraction extracted from the cerebellum as described in. **H)** Quantitative analysis of (G) as described in (D). One-way ANOVA Bonferroni test. *p<0.05, **p < 0.01, ***p<0.001.

We observed significant changes for p62, ATG9A and APT6V1A with the progression of aging from 2 to 8 and 18 months for the synaptosomes of the cortex (Fig. 5A+B). The downregulation of these components suggests a suppression of the autophagy machinery. The reduction in ATG9A indicates impaired autophagosome formation, while ATP6V1A points to poor lysosomal acidification with age. The decrease of p62 together with the other decreasing autophagy markers suggests an impaired turnover of the autophagosomes. In the cytoplasmic fraction of the cortex, we observed significantly diminished levels of ATG9A and LC3B between 2 and 8 or 18 months of age (Fig. 5C+D), implying compromised autophagosome formation and maturation with aging. Interestingly, LAMP1 levels were increased without a corresponding increase in ATP6V1A. In a recent study it was found that LAMP1 positive structures expand in size with aging that could explain the observed increase in LAMP1 levels and that the acidification varied between with lysosomes ^59^.

For the cerebellum, we detected a decrease of synaptosomal p62 and LC3B between 2 and 8 or 18 months of age (Fig. 5E+F), suggesting an impaired formation of autophagosomes with aging. The cytoplasmic compartment showed an increase of LAMP1 levels with aging (Fig. 5G+H), similarly as observed for the cortex. Here, we detected a decrease of ATP6V1A with aging, suggesting a compromised acidification of the lysosomes. We cannot exclude the possibility of a reduced number of synaptic vesicles with aging that could partially contribute to the observed decreased signals.

Taken together, our data suggest a dysregulation of autophagy with aging at both the synaptosomes and the somatic region in the murine cortex and cerebellum. The response of the specific autophagic markers however differed between the brain areas and neuronal subcompartments, suggesting that the age-associated regulation of autophagy differs between synaptosomes and cytoplasm and between brain regions as observed for the UPS (Fig. 3).

### Reduced neuronal autophagy in aging *C. elegans*

To analyze autophagy in the sub-neuronal compartments of *C. elegans*, we again used an imaging-based approach. For that, we used an established double-fluorescent reporter mCherry::GFP::LGG-1 ^60^ and generated a new transgenic *C. elegans* model by targeting the expression of mCherry::GFP::LGG-1 to different neuronal subcompartments using the *sng-1* promoter as shown above for the proteasomal reporter (Fig. 4A). LGG-1 (LC3 homolog) is fused to the pH sensitive GFP (Fig. 6A depicted in green) and pH-insensitive mCherry (Fig. 6A depicted in magenta). This reporter allows an imaging-based analysis of the formation of foci that either exhibit both fluorescence signals (autophagosomes; depicted in white) or only mCherry fluorescence (magenta) upon fusion of the autophagosome with the acidic lysosome to give rise to autolysosomes (Fig. 6A). An age-associated decline of the autophagic flux has been reported for the head neurons that however did not discriminate between neuronal subcompartments ^60^. Here, we focused our analysis on the synapses by selectively imaging the mCherry and GFP signals at the synapse. For that, we used the tandem reporter to quantify autophagosomes and autolysosomes with the progression of aging from day 3 old nematodes, to day 4 old animals, and 7 days old animals exclusively at the synapses. We observed an increase in the number of autophagosomes and no change of autolysosomes from day 3 to day 4 old animals suggesting an induction of the autophagic flux from the larval stage to the adult stage at the synapse (Fig. 6B). However, with aging (transition to day 7 of life), the number of autophagosomes and autolysosomes decreased substantially, demonstrating impaired autophagy at the synapse of nematodes (Fig. 6B). We cannot rule out the possibility of an increased retrograde transport at day 7 and that autophagosomes may fuse with lysosomes at another neuronal subcompartment e.g. the soma. We have however not observed trafficking or an increase in autophagosome or autolysosome formation at different neuronal subcompartments. To assess the autophagic flux, we treated the nematodes with chloroquine (CQ), that inhibits autophagy by impairing the fusion of the autophagosomes with lysosomes ^61^. Thus, CQ treatment should lead to an accumulation of Autophagosomes in autophagy-active nematodes. Indeed, treatment with CQ led to an increase of autophagosomes in young day 4 old animals whereas no change was observed in older day 7 old nematodes (Fig. S7). These data further support the conclusion that autophagic flux decreases with aging.

**Figure 6.**
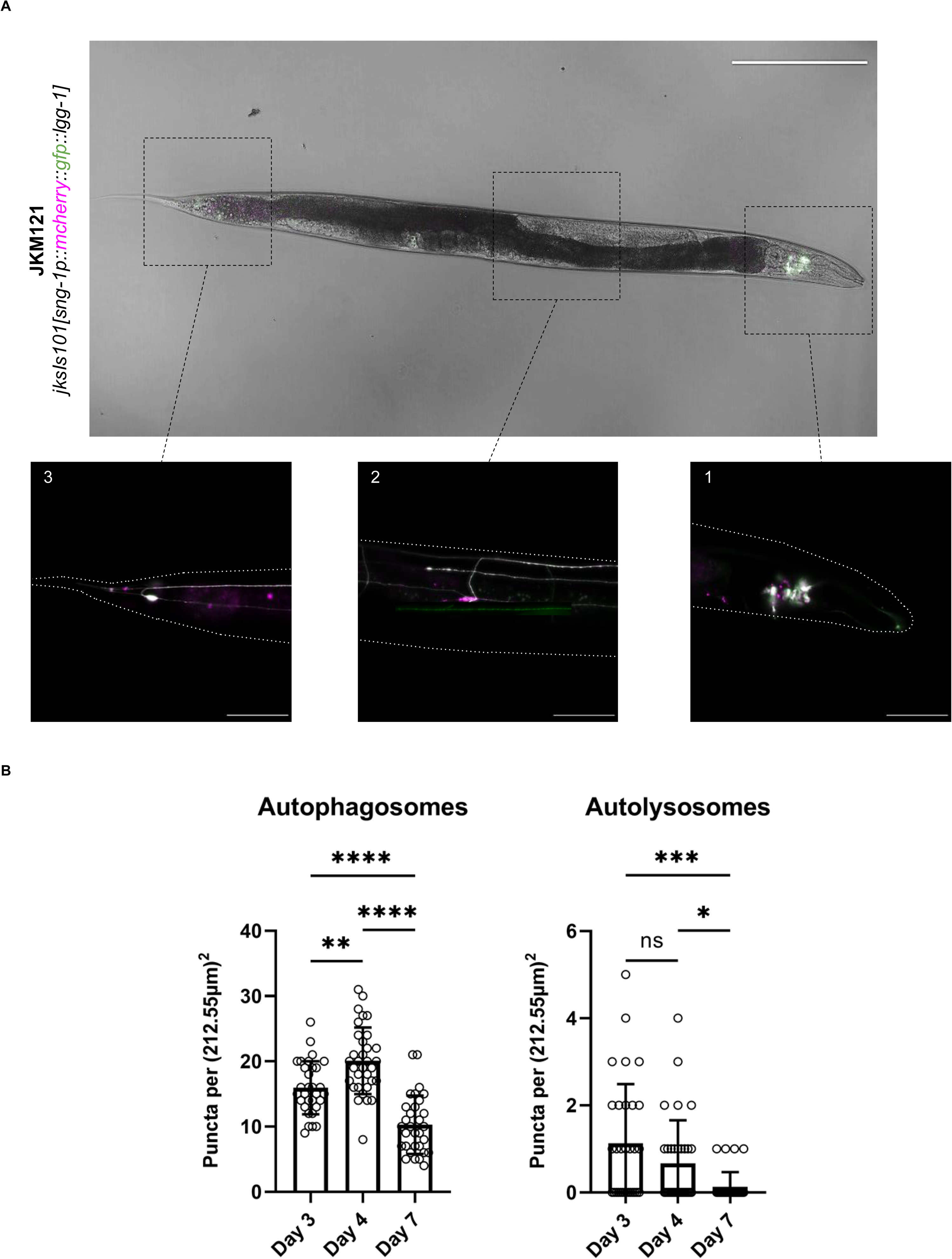
Analysis of synaptic autophagy in *C. elegans* with aging. **A)** Confocal fluorescent image of the JKM121 strain (*jksIs101[sng-1p::mcherry::gfp::lgg-1]*) expressing LGG-1 fused with both mCherry (pseudo colored in magenta) and GFP (pseudo colored in green) with close-ups of the tails (1; right), mid body (2; middle) and the head (3; left) regions. The co-localization of mCherry and GFP appears in white. The brightfield image was taken with a 10X objective, scale bar equals 200 µm. The insets were taken with a 40X objective, scale bar equals 50 µm. **B)** Scatter plots depicting the average number of autophagosomes (on the left) and autolysosomes (on the right) counted at the dorsal nerve cord of day 3 (circle), day 4 (rectangle) and day 7 (triangle) old nematodes. Experiments were carried out in triplicates, and each data point represents one animal. Significance was assessed with an ordinary one way ANOVA with Tukey’s multiple comparison test for autophagosomes and with Kruskal-Wallis test with Dunn’s post hoc test for autolysosomes (p > 0.05 (ns), p < 0.05 (*), p < 0.01 (**), p < 0.001 (***) and p < 0.0001 (****)).

In summary, our data suggest that the somatic and synaptic autophagy decreases in both animal models with aging.

## Discussion

The age-associated loss of proteostasis in the brain is a major contributor to a decline of neuronal function with aging and pathologies such as neurodegenerative diseases^62,63^. The balance of a functional neuronal proteome is regulated by protein synthesis and the degradation of proteins that are no longer required, or that are misfolded, or aggregated. It is established that protein synthesis in neurons decreases with aging ^64^, yet much less is known about the activity of protein degradation pathways in neurons and in particular at the synapses with the progression of aging ^10^.

Previous proteomic studies of protein stabilities in the brain showed that protein turnover occurs at a slower rate in aged brains ^11,65,66^. While these studies focused on the changes of the half-life times of proteins with aging, we studied here the activity of the two major proteolytic pathways, the UPS and macroautophagy, and how their capacity changes with aging. We studied the proteolytic pathways in two brain areas of mice and pan-neuronally in *C. elegans* and at different neuronal subcompartments, the cytosol (including soma, axons and dendrites) and the synapses. For that, we established and adapted new tools and workflows to allow the isolation of murine and *C. elegans* synaptosomes and generated new reporter strains for UPS and autophagy at synapses for ex vivo and in vivo analyses. Using these tools, we show that neuronal proteasomal and autophagic activity decreases with age in both model systems, suggesting that the age-associated decrease of the proteolytic activity is conserved in vertebrates and invertebrates (Fig. 7). Importantly, the age-associated decline of the proteasomal activity differs between murine brain areas. We observed that the cortex is marked by a decrease of proteasome abundance and activity, while no change could be observed in the synaptosomes of the cerebellum. In contrast, autophagy was dysregulated in both brain regions with aging. These differences in the activity of the proteolytic pathways could be a contributing factor to the selected vulnerability of different brain regions and neuron types for neurodegenerative diseases ^67^. We further detected differences in the activity of the proteasome between soma and synapses in *C. elegans*. Young nematodes exhibited a higher proteasomal activity at the synapses compared to the soma, indicating a higher degradation rate for proteasome substrate proteins at the synapse. With the progression of aging, the UPS activity also rapidly decreased at the synapses and thus confirms the observed increase in half-life times of synaptic proteins with aging that could be attributed to a reduction in proteasomal capacity ^65,66^. A recent study identified synaptic proteins as particularly vulnerable to aggregation that could contribute to their decreased degradation kinetics with aging ^66^. Loss of function of these aggregated synaptic proteins could in turn promote the age-related loss of synaptic function ^68^.

**Figure 7.**
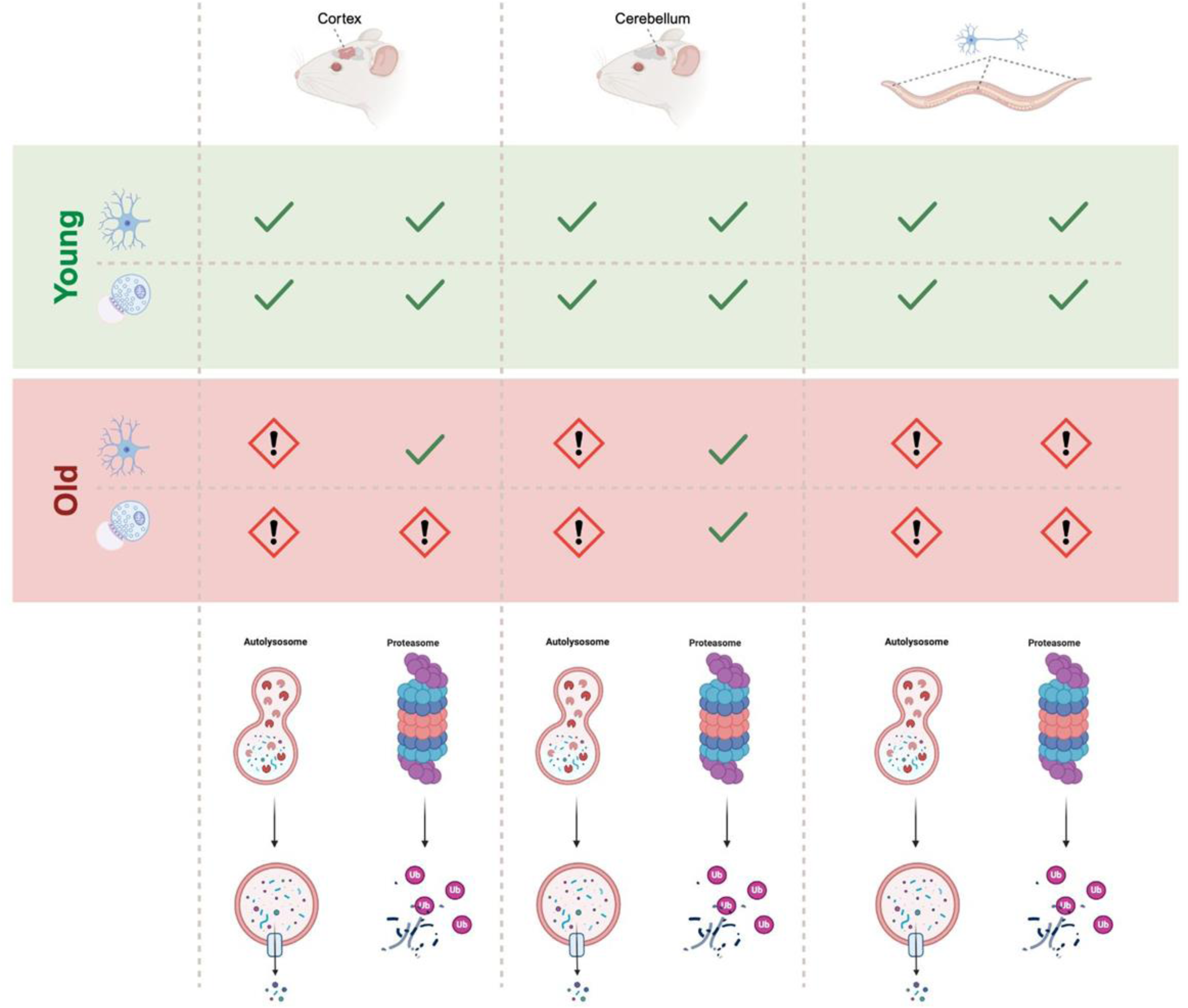
Schematic cartoon summarizing our main findings. We could uncover an age-associated decline of UPS activity in murine cortical synaptosomes, and in all neuronal subcompartments of *C. elegans* while the murine cortical somas and both, cytoplasmic and synaptosomal fractions from the cerebellum are not affected by age. The autophagic flux is perturbed upon aging in all neuronal sub-compartments and brain areas in mice and in the neurons of *C. elegans*.

Maintaining differential degradation kinetics within neurons is likely essential for sustaining specialized and compartmentalized neuronal functions. High proteolytic activity at synapses may serve a critical role in preserving synaptic integrity, as they experience intense metabolic demand, rapid protein turnover, and continuous remodeling in response to neuronal activity. Synaptic proteins are frequently exposed to oxidative stress, calcium fluctuations, and excitotoxic signals, making them particularly vulnerable to misfolding and damage. Efficient degradation at the synapse ensures the timely removal of dysfunctional proteins and allows for rapid replacement with newly synthesized proteins, thereby maintaining synaptic plasticity ^10,11,65^. With aging, a decline in local proteasomal and autophagic activity could impair this renewal process, leading to the accumulation of damaged or aggregated proteins that interfere with neurotransmission, synaptic vesicle cycling, and receptor trafficking. Consequently, diminished degradation at synapses may contribute directly to synaptic weakening and loss, hallmarks of age-related cognitive decline and neurodegeneration^65^.

In conclusion, our findings reveal that aging is accompanied by a conserved decline in proteasomal and autophagic activity across species, with differences in brain regions, and neuronal compartments, suggesting that reduced proteolytic capacity, particularly at synapses, may underlie the accumulation and aggregation of synaptic proteins and contribute to the age-related deterioration of neuronal and cognitive function. The observation that the proteolytic capacity of different brain regions is differentially affected by aging, poses the question whether an induction of e.g. the UPS in the cortex could preserve neuronal function with aging and may represent a therapeutic target for neurodegenerative diseases.

## Experimental procedures

### Molecular cloning and generation of transgenic *C. elegans*

The genomic DNA of the *C. elegans* N2 strain (wildtype) was extracted using Phire Tissue Direct PCR Master Mix (F-170, Thermo Fisher Scientific), and the synaptogyrin-1 promoter (*sng-1p*) sequence was amplified using 5’-TTAATTGTTAATTATCTAAGCTTGT-3’ and 5’-GCTAAAATAAAAGAAATATAGAGG-3’ as forward and reverse primers respectively. The *sng-1p::mCherry::GFP::lgg-1* plasmid pJK101 was created using pMH1130 (a kind gift from the lab of Malene Hansen, Buck Institute, USA), through a substitution of the *rgef-1* promoter by the *sng-1* promoter via Gibson assembly (Gibson assembly master mix E2611, BioLabs). In the same manner, we created the *sng-1p::UbG76V::Dendra2* plasmid pJK102 which expresses a mutated ubiquitin moiety fused with the photoconvertible fluorophore Dendra2. The Dendra2 sequence was amplified from a previously designed plasmid containing dendra2 (pPD95_77-hDBN(S647D)-Dendra2) and ubiquitin (G76V) was amplified from Ub-G76V-GFP (plasmid #11941 from Addgene). We used pJK101 as a backbone to replace *mCherry::GFP::lgg-1* with *Ub(G76V)::Dendra2* through a one reaction Gibson assembly. The correct sequence of all constructs was verified by DNA sequencing.

To create transgenic animals, 20 ng/µl pJK101 was co-injected with 120 ng/µl of DNA filler (DNA ladder) into N2 by InVivo Biosystems (USA). The extrachromosomal array was then integrated using UV illumination and backcrossed four times (JKM121). 30 ng/µl of pJK102 plasmid with 100 ng/µl of 100 bp DNA ladder as filler were microinjected into BR584 *[unc-119(ed4)]* to obtain JKM140.

Below is the complete list of primers used for cloning:

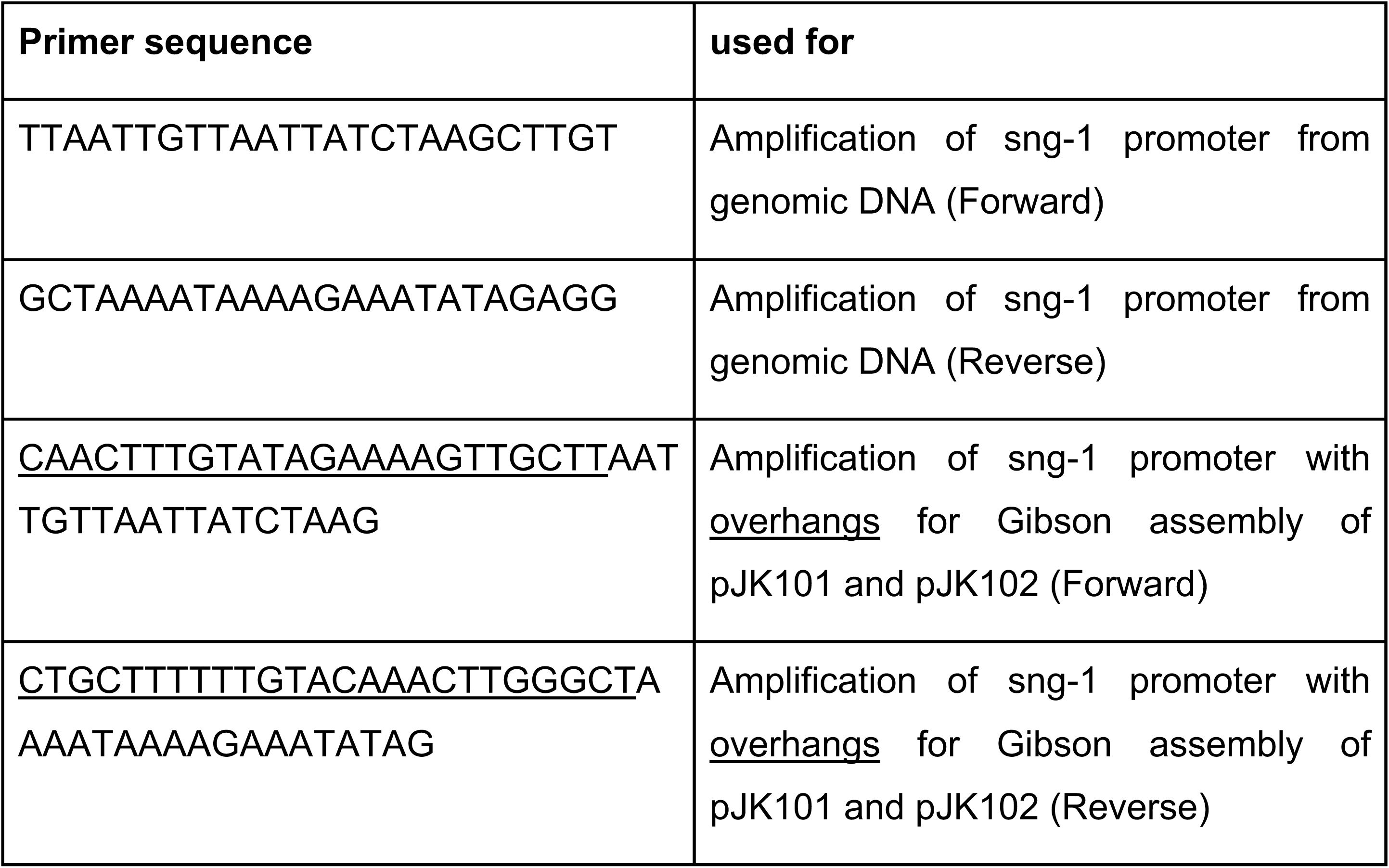
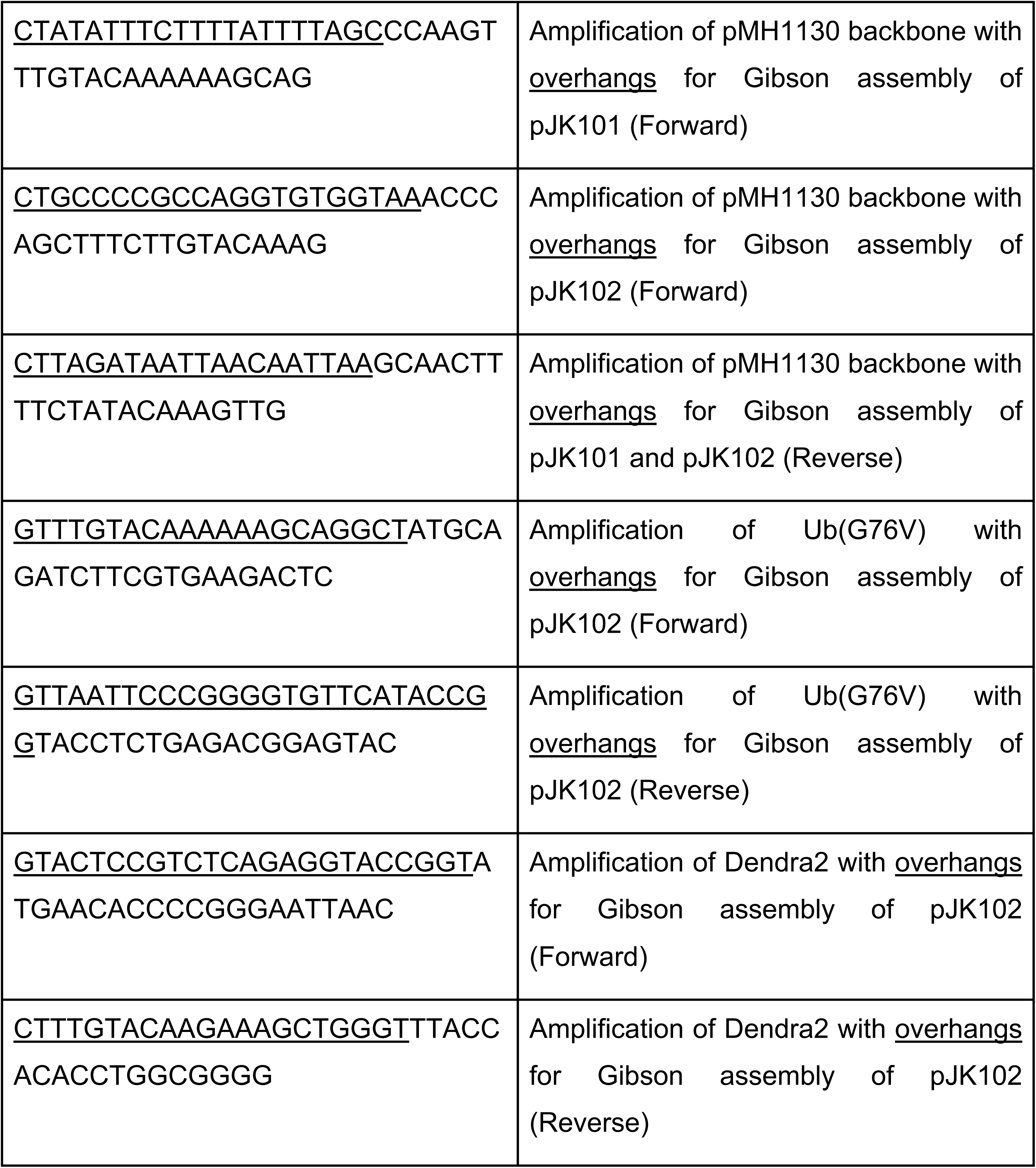

### *C. elegans* maintenance

All nematodes were maintained at 20°C on 6 cm nematode growth medium (NGM) or 10 cm high growth medium (HGM; for CQ574) plates seeded with *E. coli* OP50 as a food source. The strains were synchronized by egg-laying and picked at day 3, 4 or 7 of life for experiments or by bleaching. The *C. elegans* strains used in this study are listed in the table below.

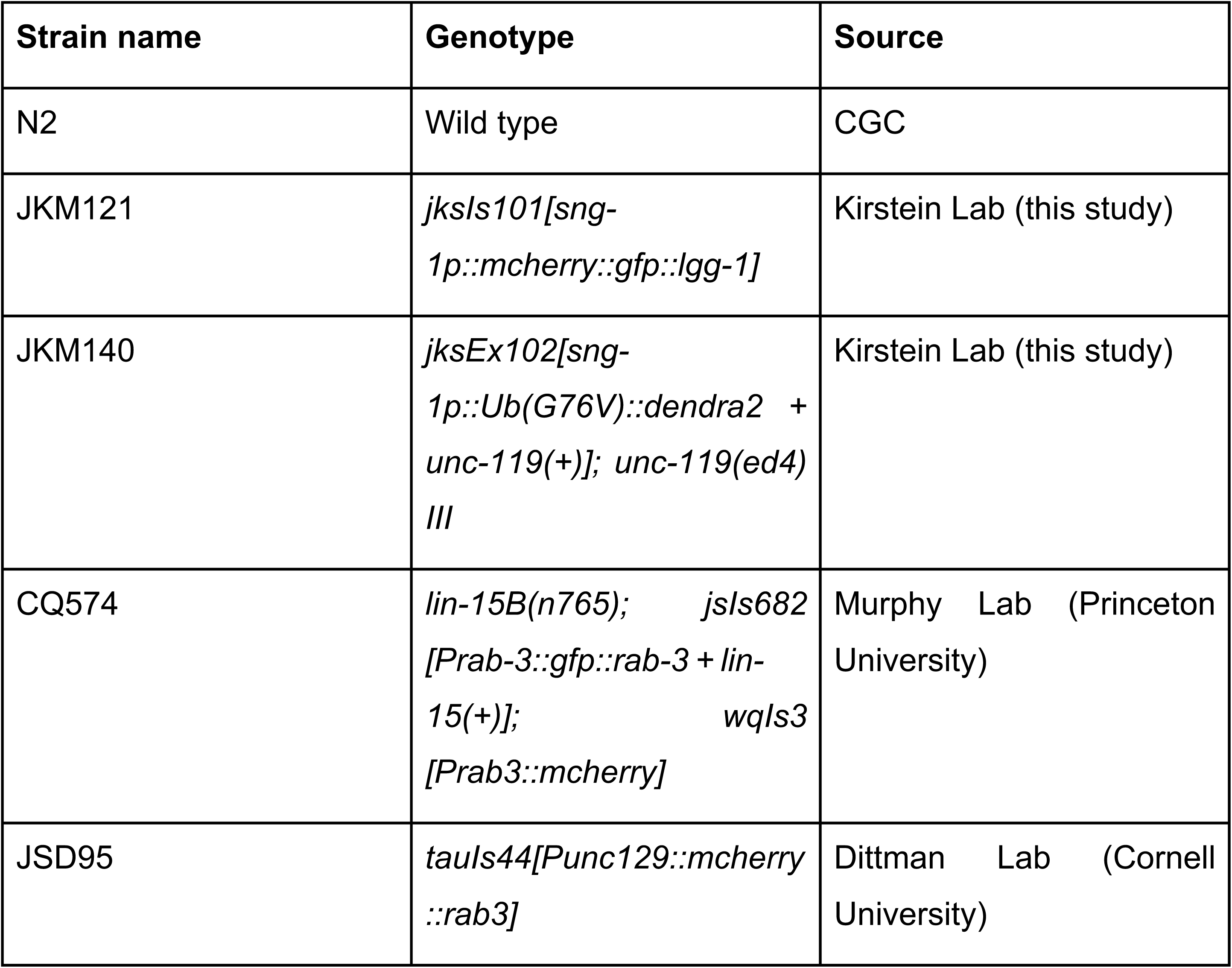

### C. elegans crossing

To verify a synaptic expression of the sensors in the novel strains, we employed a strain expressing RAB3 fused with mCherry at synapses of the D-type motor neurons (JSD95, a kind gift from Jeremy Dittman’s lab at Cornell University). We induced male generation of JSD95 through heat shock and crossed them with hermaphrodites of JKM140. The cross was then used to detect the potential co-localization between the red signal (mCherry from JSD95) and green signal (Dendra2 from JKM140) and thus verifying a synaptic localization.

### C. elegans imaging

The nematodes were mounted on a 3% agarose pad, anesthetized with 250 mM sodium azide and imaged using a confocal laser scanning microscope (LSM-880, Zeiss). Whole animal images were acquired with an EC Plan-Neofluar 10x/0.3 M27 objective. For autophagy readouts, Z-stacks at 0.6 µm interval of the dorsal nerve cord at the mid body and tail region were taken using a Plan-Apochromat 40x/1.4 Oil DIC M27 objective. GFP and mCherry were excited using an Argon laser at 493 nm and HeNe laser at 543 nm respectively, the gain and laser intensity were maintained for all acquired images. Using Fiji ImageJ, Z-stacks from each animal were combined into one tiff with maximum intensity projections, and yellow or red-only puncta were counted manually and in a blinded manner.

### *C. elegans* chloroquine treatment

To study the autophagic flux, age-synchronized animals were subjected to chloroquine (CQ, Sigma Aldrich) which impairs autophagy by blocking the fusion of lysosomes with autophagosomes ^61^. The nematodes were incubated in liquid culture supplemented with 20 mM CQ or ddH2O in low binding tubes wrapped with aluminum foil for 20 hours at 20°C on a roller shaker. After the incubation time, nematodes were recovered on freshly seeded NGM plates and imaged as described above.

### Rodent procedures

All mice used were handled in accordance with the relevant guidelines and regulations. Protocols were approved by the ‘Landesamt für Gesundheit und Soziales’ (LaGeSo; Regional Office for Health and Social Affairs) in Berlin and animals reported under the permit number T0347/11. All animals used in this work were males of genetic background C57 Bl/6J.

### Synaptosomal preparation

Synaptosomes were isolated according to a previous protocol ^69^. In brief, mouse cortex with subcortical regions and cerebellum were weighed and homogenized in 10 ml per 1 g of tissue in homogenization buffer (5 mM Tris, 1 mM EDTA, 0.32 M sucrose, pH 7.4) supplemented with protease inhibitors (Calbiochem set III) and phosphatase inhibitors (1 mM Na2MO4,1 mM NaF, 20 mM β-glycerophosphate, 1 mM Na3VO4, 500 nM cantharidin) and homogenized using a glass pestle. The lysate was cleared of nuclei and cell debris by centrifugation at 1000 xg for 10 minutes. Supernatant (S1) was collected, aliquoted and further centrifuged at 15.000 xg for 30 minutes, after which the soluble supernatant cytoplasmic fraction (S2) was collected and snap frozen in liquid N2. The remaining pellet (P2) containing the synaptosomes with myelin, membranes and mitochondria was snap frozen in liquid N2.

### *C. elegans* synaptic fraction isolation

Dissociation of total cell population from *C. elegans* was done as described previously^70^. In brief, synchronized (∼6000 nematodes per day per replicate) day 4 (young adult), day 7 or day 10 old CQ574 nematodes (a kind gift from Coleen Murphy’s lab at Princeton University) were washed five times with M9 buffer followed by treatment with 300 μl of lysis buffer (200 mM DTT, 0.25% SDS, 20 mM Hepes pH 8.0, 3% sucrose) for 6 minutes. The pellet was washed five times in M9 to remove the lysis buffer followed by treatment with 20 mg/ml Pronase from *Streptomyces griseus* (Sigma) for 20 minutes. The pellet was mechanically disrupted every 2 minutes during this incubation. When most nematodes had dissociated, the reaction was stopped by the addition of 900 μl ice-cold Lebovitz’s L-15 medium (Sigma). The cells were washed, centrifuged and filtered using a 5 μm filter for the subsequent FACS procedure. Cell sorting was done with a BD FACS Fusion 5-Laser system (2UV,6V,2B,5YG,3R) with BD FACS Diva Software version 9.0.1 with 70 µm nozzle. As a control, we used cells isolated from N2 (wild type) prepared in parallel to define the different gates for detection. DAPI was added to the sample with a final concentration 10ng/ml. GFP+ (FITC-Channel, 530/30BP, 502LP), mCherry+ (PE-Texas-Red-Channel 610/20 BP, 600LP), and DAPI (450/50 BP) filter sets and respective channels were used to detect single and double positive populations after gating on single events.

### Sample preparation for proteomics analysis

For the proteomics sample preparation 6000 synchronized nematodes of the appropriate age were used for total cell isolation and FACS analysis as described in the methods section. The sorted fractions varied in counts from 20,000 – 100,000 events per sample. The sorted fractions were homogenized and lysis buffer (fc 4% SDS, 100 mM HEPES, pH 8.5, 50 mM DTT) was added to each sample. Samples were then boiled at 95°C for 10 min and sonicated using a tweeter. Reduction was followed by alkylation with 200 mM iodoacetamide (IAA, final concentration 15 mM) for 30 min at room temperature in the dark. Samples were acidified with phosphoric acid (final concentration 2.5%), and seven times the sample volume of S-trap binding buffer was added (100 mM TEAB, 90% methanol). Samples were bound on 96-well S-trap micro plate (Protifi) and washed three times with binding buffer. Trypsin in 50 mM TEAB pH 8.5 was added to the samples (1 µg per sample) and incubated for 1 h at 47°C. The samples were eluted in three steps with 50 mM TEAB pH 8.5, elution buffer 1 (0.2% formic acid in water) and elution buffer 2 (50% acetonitrile and 0.2% formic acid). The eluates were dried using a speed vacuum centrifuge (Eppendorf Concentrator Plus, Eppendorf AG, Germany) and stored at -20° C. Before analysis, samples were reconstituted in in MS Buffer (5% acetonitrile, 95% Milli-Q water, with 0.1% formic acid), spiked with iRT peptides (Biognosys, Switzerland) and loaded on Evotips (Evosep) according to the manufacturer’s instructions. In short, Evotips were first washed with Evosep buffer B (acetonitrile, 0.1% formic acid), conditioned with 100% isopropanol and equilibrated with Evosep buffer A. Afterwards, the samples were loaded on the Evotips and washed with Evosep buffer A. The loaded Evotips were topped up with buffer A and stored until the measurement.

### LC-MS Data independent analysis (DIA)

Peptides were separated using the Evosep One system (Evosep, Odense, Denmark) equipped with a 15 cm x 150 μm i.d. packed with a 1.5 μm Reprosil-Pur C18 bead column (Evosep performance, EV-1137, Denmark). The samples were run with a pre-programmed proprietary Evosep gradient of 44 min (30 samples per day) using water and 0.1% formic acid and solvent B acetonitrile and 0.1% formic acid as solvents. The LC was coupled to an Orbitrap Exploris 480 (Thermo Fisher Scientific, Bremen, Germany) using PepSep Sprayers and a Proxeon nanospray source. The peptides were introduced into the mass spectrometer via a PepSep Emitter 360-μm outer diameter × 20-μm inner diameter, heated at 300°C, and a spray voltage of 2 kV was applied. The injection capillary temperature was set at 300°C. The radio frequency ion funnel was set to 30%. For DIA data acquisition, full scan mass spectrometry (MS) spectra with a mass range of 350–1650 m/z were acquired in profile mode in the Orbitrap with a resolution of 120,000 FWHM. The default charge state was set to 2+, and the filling time was set at a maximum of 20 ms with a limitation of 3 × 10^6^ ions. DIA scans were acquired with 40 mass window segments of differing widths across the MS1 mass range. Higher collisional dissociation fragmentation (normalized collision energy 30%) was applied, and MS/MS spectra were acquired with a resolution of 30,000 FWHM with a fixed first mass of 200 m/z after accumulation of 1 × 10^6^ ions or after filling time of 45 ms (whichever occurred first). Data were acquired in profile mode. For data acquisition and processing of the raw data, Xcalibur 4.4 (Thermo) and Tune version 4.0 were used.

### Proteomic data processing

DIA raw data were analyzed using the directDIA pipeline in Spectronaut v.18 (Biognosys, Switzerland) with BGS settings besides the following parameters: Protein LFQ method= QUANT 2.0, Proteotypicity Filter = Only protein group specific, Major Group Quantity = Median peptide quantity, Minor Group Quantity = Median precursor quantity, Data Filtering = Qvalue, Normalizing strategy = Local Normalization. The data were searched against a species specific (*M. musculus*, 16,747 entries, v. 160106 and *C. elegans,* 26,667 entries, v. 160106) and a contaminants (247 entries) Swissprot database. The identifications were filtered to satisfy FDR of 1 % on peptide and protein level. The data were then exported and further analyzed in Rstudio using MSStats removing sparse precursors and with a cutoff for missing values of 0.5. A pairwise comparison of the protein quantification and differential expression table used for volcano plots generation and Principal Component Analysis (PCA), respectively, using R version 4.1.3 and RStudio server version 1.1.463.

Murine protein groups were considered as significantly enriched if they displayed a Q value < 0.05 and fold change > 1.5 for all conditions. For the *C. elegans* synaptic proteomics, lower protein input resulted in very few proteins passing an FDR < 0.05 threshold. To identify age-regulated candidates while maintaining a degree of stringency, we applied a relaxed threshold of unadjusted p < 0.01 together with an absolute fold-change ≥ 2, as commonly used in label-free proteomics datasets with limited replication ^71,72^. These proteins should be considered putative candidates that require independent validation, and we focus our biological interpretation primarily on pathway- and GO-level changes rather than on individual proteins.

GO term and KEGG pathway enrichment for age-regulated proteins was performed using WormEnrichr (Biological Process, Molecular Function, Cellular Component, KEGG). Enrichment p-values were calculated by the WormEnrichr pipeline (Fisher’s exact/hypergeometric test) and adjusted for multiple testing using the Benjamini–Hochberg procedure. Only terms with adjusted p-value (q-value) < 0.05 were considered significantly enriched and reported.

### Proteasome activity assays

The chymotrypsin-like activity of the proteasome was analyzed in a 96-well plate using 20 µg (S2 fraction) or 10 µg (P2 fraction) of protein lysate of murine brain areas. The lysate was mixed with 100 µM of Suc-LLVY-AMC (Enzo, BML-P802) in duplicates. As a control, the protein samples were treated with the proteasome inhibitor MG132 (Sigma, C2211) for 30 minutes before the addition of the Suc-LLVY-AMC. The chymotrypsin-like activity of the proteasome can cleave and release AMC that is then an active fluorophore. The fluorescence (380 nm ex/460 nm em) was measured every 5 minutes for 2 hours in a microplate reader (Tecan). For each sample, a curve showing the variation of fluorescence over time was plotted and its slope was normalized to that of the respective control.

In *C. elegans*, the proteasome activity was measured by photoconversion experiments *in vivo* using the newly generated strain JKM140. The degradation rate of the photoconverted mutated ubiquitin fused to Dendra2 (Ub^G76V^-Dendra2) was measured as the remaining fluorescence at 561 nm after 1 hour post photo-conversion as previously described and established ^53,55,73^. A region of interest was selected and imaged before photoconversion, then photobleached with a 405 nm laser and imaged immediately (considered T0) and again after 1 hour (T1) on a laser scanning microscope LSM-880 (Zeiss). The fluorescence intensities at T0 and T1 were quantified using Fiji ImageJ. The T1 fluorescence intensity normalized to the relative T0 intensity of each animal was depicted as a negative correlation to the UPS activity. No diffusion of the photoconverted signal was observed within the time period of the experiment.

### SDS-PAGE and western blotting

An aliquot of the P2 was resuspended in the same volume as the S2 fraction and the protein amount was quantified using BCA Thermo Scientific Pierce™ Protein Assay. To study autophagy, 10 µg of protein was subjected to SDS-polyacrylamide gel electrophoresis (SDS-PAGE) and subsequent Western blot analysis was performed as previously described ^74^ with the exception of the use of PVDF membrane for the anti-LC3 and anti-p62 antibodies. Gels to visualize the abundance of the 20S ɑ-subunits were transferred to nitrocellulose membranes using the standard program (30 min at 25 V) of the Trans-Blot Turbo Transfer system (BioRad). The antibodies used were anti-p62 (Abcam ab56416 1:1000), anti-LC3B (Novus Biologicas NB100-2220 1:1000), anti α-Tubulin (Sigma-Aldrich T6199 1:20000 and T5168 1:2000), anti-LAMP1 (Cell Signaling Technology C54H11 1:1000), anti-ATP6V1A(ab199326 abcam, 1:1000), anti-ATG9A (Novus Biologica NB110-56893, 1:1000), anti-synaptophysin (Sigma-Aldrich S5768, 1:1000), anti-PSD95 (Antibodies Incorporated 75-028 1:1000), anti-GAPDH (Sigma-Aldrich CB1001, 1:10.000) and anti-20S ɑ-subunits (Enzo BML-PW8195 1:1000). Quantification of signal intensities was performed using Fiji. For relative quantifications, measurements were normalized to loading control (tubulin).

### Ultrastructure of synaptosomes

Freshly isolated synaptosomal fractions were washed in saline buffer (SB; 140 mM NaCl + 5 mM KCl + 5 mM NaHCO3 + 1.2 mM NaH2PO4 + 1 mM MgCl2) and the pellet was fixed overnight at 4°C with a solution containing 2.5% glutaraldehyde (GA), 1.25% paraformaldehyde (PFA) and 0.03% picric acid in SB buffer (pH 7.4). After fixation, pellets were washed in SB buffer at 4°C and postfixed with 1% tannic acid in SB for 1h at 4°C in the dark, and after washing in SB samples were additionally postfixed with 1% osmium tetroxide (OsO4) in SB buffer also for 1h at 4°C in the dark. Samples were then washed in cold water at room temperature (RT) followed by maleate buffer (MB) at RT. After washing with MB, samples were block-stained with 1% uranyl acetate in 0.05 M MB pH 5.5 for 1h at RT in the dark. The samples were again washed in cold water in the dark and then dehydrated in a graded series of ethanol (30%, 50%, 70%, 90%, 2x 100%) for at least 5 minutes each. The samples were then infiltrated in Hydroxypropylmethacrylate (HPMA) for 1h at RT and then in a 1:1 mixture of HPMA and Epon overnight at RT. The next day, the samples were embedded in pure Epon and finally polymerized at 60°C for at least 48h. Ultrathin sections (60 nm) were cut on a Reichert Ultracut-S microtome (Leica), mounted on formvar-coated 200 mesh copper grids (Plano), poststained with 2% aqueous uranyl acetate solution and lead citrate (according to Reynolds solution), and examined in a Zeiss transmission electron microscope 900 (Zeiss-TEM-900) at 80 kV. Images were taken using a digital camera (Proscan 2K Slow-Scan CCD camera; Tröndle).

### Statistical analyses

For statistical analyses, GraphPad Prism was employed. One-way ordinary ANOVA (with Tukey‘s test post hoc), one-way Brown-Forsythe and Welch ANOVA (with Dunnett‘s T3 post hoc test) and normality tests were performed. For the murine experiments, unless stated otherwise, technical duplicates were used in N=4 animals. The *C. elegans* experiments were executed in triplicates. P-values were considered significant when p < 0.05 (*), p < 0.01 (**), p < 0.001 (***) and p < 0.0001 (****).

## Acknowledgments

We thank Malene Hansen (The Buck Institute, USA) for pMH1130 plasmid, Coleen Murphy (Princeton University, USA) and Rachel Arey (Baylor College of Medicine, USA) for strain CQ574 and FACS instructions and Jeremy Dittman (Cornell University, USA) for strain JSD95. We further acknowledge the support of the core facilities of Imaging, FACS and Proteomics of the FLI.

## Data availability

The mass spectrometry data are stored and available in the MassIVE Repository.

Dataset for the mice synaptosomes:

http://massive.ucsd.edu/ProteoSAFe/status.jsp?task=48afc64ef4be463e8e282da039 8891c1

Data set for *C. elegans* experiments:

http://massive.ucsd.edu/ProteoSAFe/status.jsp?task=df7471d94a4a47b69b85dd1d6 c1136be

Password for both data sets is: FLIReviewer_2025

## Contributions

Conceptualization: Mira Sleiman, Dimitra Ranti, Britta J. Eickholt, Janine Kirstein; experiments performed and data analysis: Mira Sleiman, Dimitra Ranti, Sudarson Baskaran, Patricia Kreis, Saskia Muth, Christian Gallrein, Agnieszka Münster-Wandowski, Norman Rahnis, Emilio Cirri, Britta J. Eickholt, Janine Kirstein; writing of the manuscript: Britta J. Eickholt, Janine Kirstein; edited and revised manuscript: Mira Sleiman, Dimitra Ranti, Sudarson Baskaran, Patricia Kreis, Saskia Muth, Christian Gallrein, Britta J. Eickholt, Janine Kirstein; funding acquisition: Britta J. Eickholt, Janine Kirstein; supervision: Britta J. Eickholt, Janine Kirstein.

## Supplementary Information

Figure S1 Depth and consistency of proteome coverage across ages and compartments in mouse and *C. elegans* datasets. **A)** Total number of quantified protein groups identified in mouse cortical synaptosome preparations from 2-, 8-, and 18-month-old animals across all biological replicates, showing comparable depth of coverage between age groups. **B)** Total number of quantified protein groups in *C. elegans* Day 4 and Day 7 samples (N = 3; ∼6000 nematodes per experiment) for whole neurons (GFP^+^/mCherry^+^), synaptic compartments (GFP^+^), and neuronal cell bodies (mCherry^+^), demonstrating similar proteome depth between ages and compartments. **C)** Total number of quantified protein groups in *C. elegans* Day 4 and Day 10 samples (N = 3; ∼6000 nematodes per experiment) for the same three neuronal fractions, again indicating stable identification numbers across ages and compartments.

Figure S2 Gene Ontology Biological Process enrichment across aging mouse cortical synaptosomes. Differentially abundant proteins (>1.5-fold change, Q < 0.05) were analyzed with Enrichr using the GO Biological Process ontology. Bar plots display significantly enriched terms (adjusted Q < 0.05) for pairwise comparisons between 2-, 8-, and 18-month-old cortical synaptosome proteomes: **A)** terms enriched in 2- versus 18-month synaptosomes, **B)** terms depleted in 2- versus 18-month synaptosomes, **C)** terms enriched in 2- versus 8-month synaptosomes, **D)** terms depleted in 2- versus 8-month synaptosomes. Bars represent the Enrichr combined enrichment score (−log P value multiplied by the odds ratio); the higher values indicate stronger over-representation of the corresponding GO term in the indicated comparison.

Figure S3 Gene Ontology Biological Process enrichment across aging mouse cortical synaptosomes. Differentially abundant proteins (>1.5-fold change, Q < 0.05) were analyzed with Enrichr using the GO Biological Process ontology. Bar plots display significantly enriched terms (adjusted Q < 0.05) for pairwise comparisons between 8-and 18-month-old cortical synaptosome proteomes. GO analysis for the depleted candidates did not have significant (Q<0.05) hits across GO data bases (due to low number of candidate hits) and hence not shown. Bars represent the Enrichr combined enrichment score (−log P value multiplied by the odds ratio); the higher values indicate stronger over-representation of the corresponding GO term.

Figure S4 Gene Ontology Biological Process enrichment across aging *C. elegans* synapses. Differentially abundant proteins (>2-fold change, P < 0.01) were analyzed with Enrichr using the GO Biological Process ontology. Bar plots display significantly enriched terms (adjusted Q < 0.05) for pairwise comparisons between 4-, 7-, and 10-day-old synapse proteomes: **A)** terms enriched in 4- versus 7-day old synapses, **B)** terms depleted in 4- versus 7-day old synapses, **C)** terms enriched in 4- versus 10-day old synapses, **D)** terms depleted in 4- versus 10-day old synapses. Bars represent the Enrichr combined enrichment score (−log P value multiplied by the odds ratio); the higher values indicate stronger over-representation of the corresponding GO term in the indicated comparison.

Figure S5 Sunburst representations of GO enrichment for synaptic proteins during aging in *C. elegans*. Differentially abundant proteins (>2-fold change, P < 0.01) from day-4 versus day-7 and day-4 versus day-10 synaptic proteomes were analyzed using the SynGO analysis tool by Gene Ontology Biological Process and Cellular Component categories. **A)** Sunburst map of Biological Process terms enriched in day-4 versus day-7 synapses. **B)** Corresponding Cellular Component map for enriched proteins in day-4 versus day-7 synapses. **C)** Sunburst map of Biological Process terms enriched in day-4 versus day-10 synapses. **D)** Cellular Component map for enriched proteins in day-4 versus day-10 synapses. Segment color encodes the number of proteins annotated to each GO term (gene count), with warmer colors indicating higher counts.

Figure S6 Raw representative images of western blot membranes probing the 20S alpha subunits of the proteasome. Anti-Tubulin is used as a control. The membranes are from one trial (out of 2) of samples from cortical cytoplasmic fraction (upper left), cortical synaptosomes (upper right), cerebellar cytoplasmic fraction (lower left) and cerebellar synaptosomes (lower right). For each membrane, samples from 4 animals per age group (2 months old, 8 months old and 18 months old) are loaded consecutively. A protein ladder is loaded on the far left and far right of each membrane, with the molecular weights indicated in kDa.

Figure S7 Scatter plots depicting the average number of autophagosomes (in yellow) and autolysosomes (in red) counted at the dorsal nerve cord of day 4 and day 7 old nematodes treated with chloroquine (20 mM) or ddH_2_O as control. Experiments were carried out in triplicates, and each data point represents one animal. Significance was assessed with an ordinary two way ANOVA with Tukey’s multiple comparison post hoc test (p > 0.05 (ns), p < 0.05 (*), p < 0.01 (**), p < 0.001 (***) and p < 0.0001 (****)).

