## supplemental table and figures for "Comparative analysis of neuronal proteolytic pathways reveals neuron-specific and sub-compartmental-specific capacities with aging"

S 1

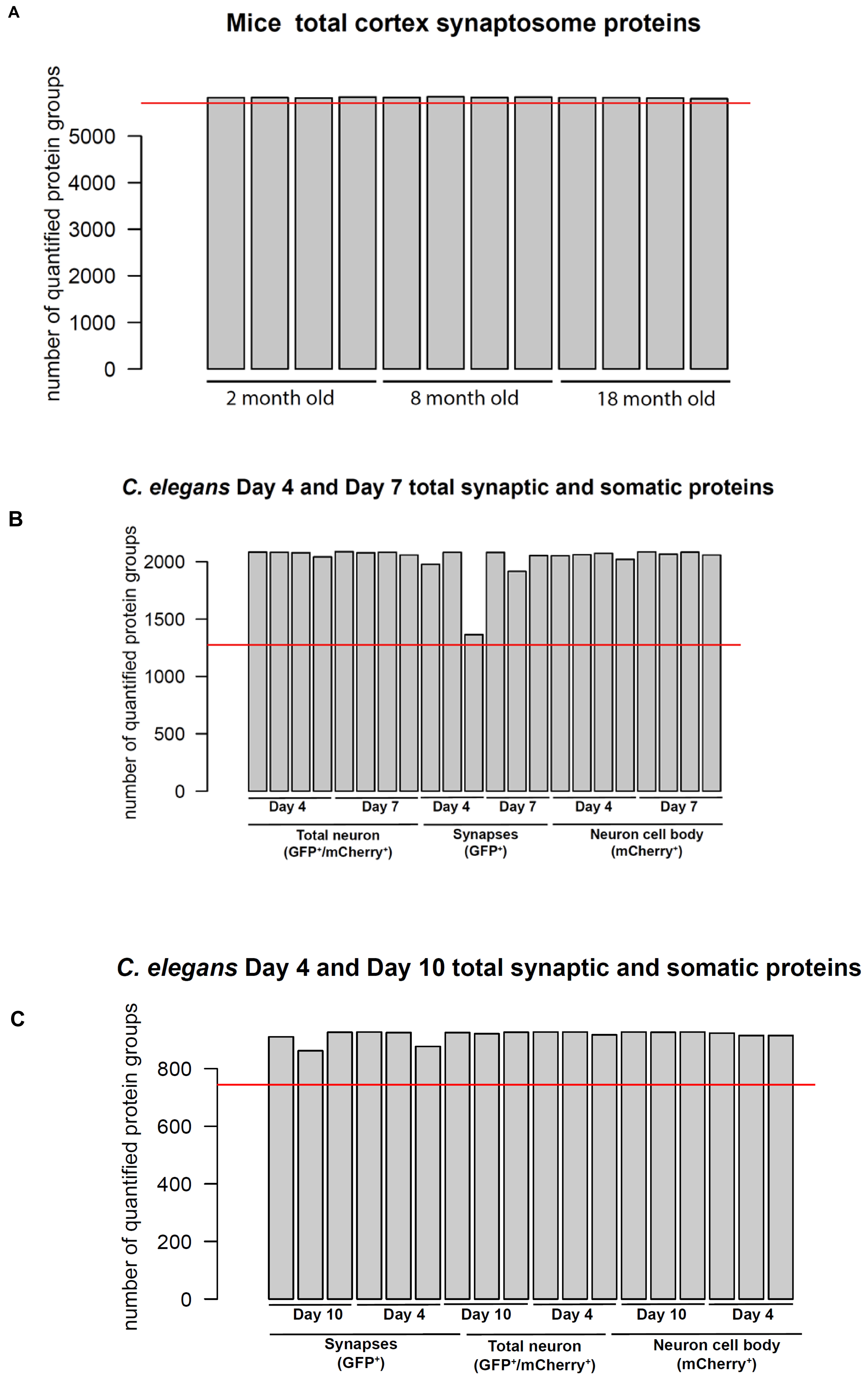
A
Mice total cortex synaptosome proteins
number of quantified protein groups
0
1000
2000
3000
4000
5000
2 month old
8 month old
18 month old
B
C. elegans Day 4 and Day 7 total synaptic and somatic proteins
number of quantified protein groups
0
500
1000
1500
2000
Day 4
Day 7
Day 4
Day 7
Day 4
Day 7
Total neuron (GFP⁺/mCherry⁺)
Synapses (GFP⁺)
Neuron cell body (mCherry⁺)
C
C. elegans Day 4 and Day 10 total synaptic and somatic proteins
number of quantified protein groups
0
200
400
600
800
Day 10
Day 4
Day 10
Day 4
Day 10
Day 4
Synapses (GFP⁺)
Total neuron (GFP⁺/mCherry⁺)
Neuron cell body (mCherry⁺)

S2

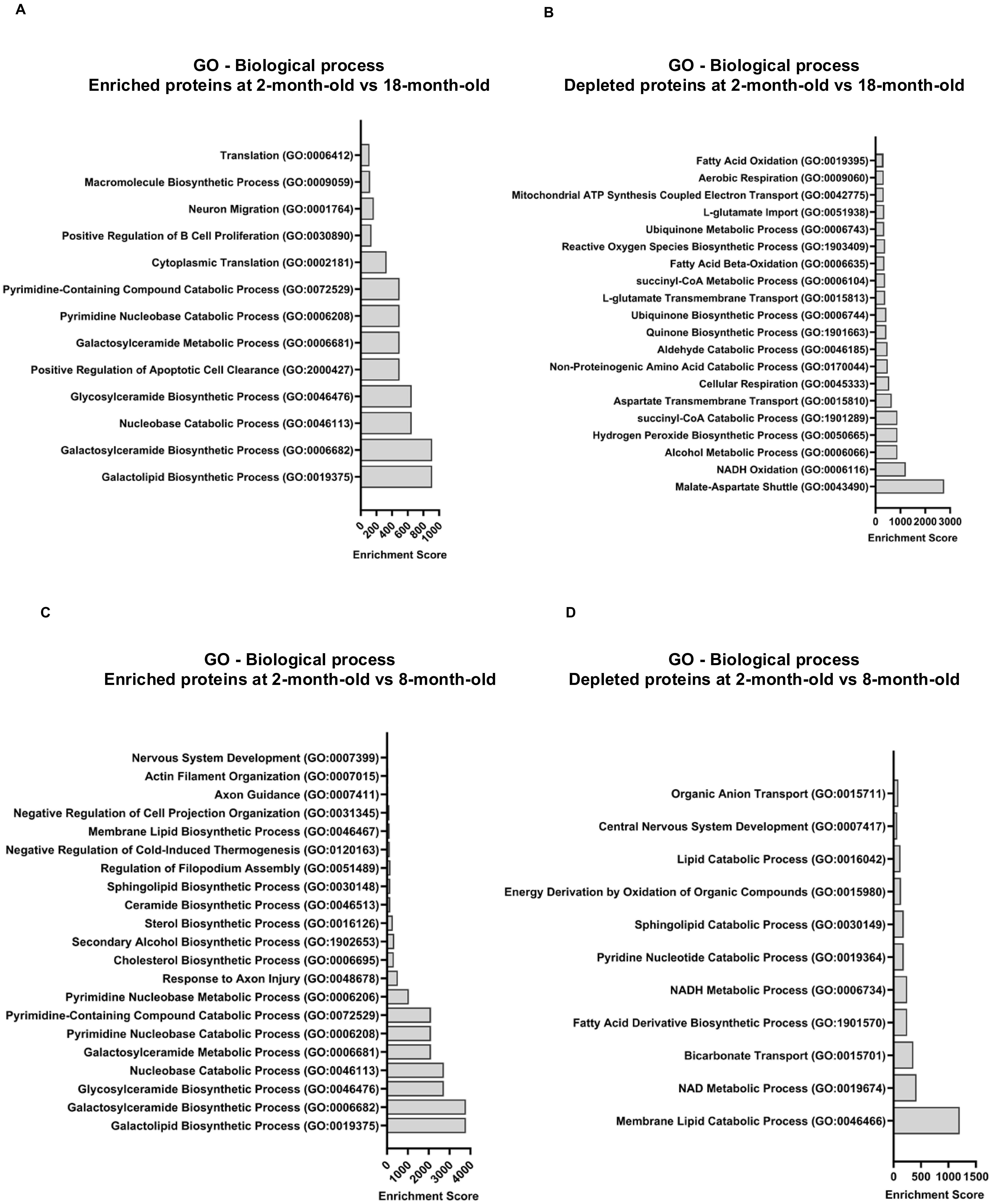
A
GO - Biological process
Enriched proteins at 2-month-old vs 18-month-old
Translation (GO:0006412)
Macromolecule Biosynthetic Process (GO:0009059)
Neuron Migration (GO:0001764)
Positive Regulation of B Cell Proliferation (GO:0030890)
Cytoplasmic Translation (GO:0002181)
Pyrimidine-Containing Compound Catabolic Process (GO:0072529)
Pyrimidine Nucleobase Catabolic Process (GO:0006208)
Galactosylceramide Metabolic Process (GO:0006681)
Positive Regulation of Apoptotic Cell Clearance (GO:2000427)
Glycosylceramide Biosynthetic Process (GO:0046476)
Nucleobase Catabolic Process (GO:0046113)
Galactosylceramide Biosynthetic Process (GO:0006682)
Galactolipid Biosynthetic Process (GO:0019375)
0 200 400 600 800 1000
Enrichment Score
B
GO - Biological process
Depleted proteins at 2-month-old vs 18-month-old
Fatty Acid Oxidation (GO:0019395)
Aerobic Respiration (GO:0009060)
Mitochondrial ATP Synthesis Coupled Electron Transport (GO:0042775)
L-glutamate Import (GO:0051938)
Ubiquinone Metabolic Process (GO:0006743)
Reactive Oxygen Species Biosynthetic Process (GO:1903409)
Fatty Acid Beta-Oxidation (GO:0006635)
succinyl-CoA Metabolic Process (GO:0006104)
L-glutamate Transmembrane Transport (GO:0015813)
Ubiquinone Biosynthetic Process (GO:0006744)
Quinone Biosynthetic Process (GO:1901663)
Aldehyde Catabolic Process (GO:0046185)
Non-Proteinogenic Amino Acid Catabolic Process (GO:0170044)
Cellular Respiration (GO:0045333)
Aspartate Transmembrane Transport (GO:0015810)
succinyl-CoA Catabolic Process (GO:1901289)
Hydrogen Peroxide Biosynthetic Process (GO:0050665)
Alcohol Metabolic Process (GO:0006066)
NADH Oxidation (GO:0006116)
Malate-Aspartate Shuttle (GO:0043490)
0 1000 2000 3000
Enrichment Score
C
GO - Biological process
Enriched proteins at 2-month-old vs 8-month-old
Nervous System Development (GO:0007399)
Actin Filament Organization (GO:0007015)
Axon Guidance (GO:0007411)
Negative Regulation of Cell Projection Organization (GO:0031345)
Membrane Lipid Biosynthetic Process (GO:0046467)
Negative Regulation of Cold-Induced Thermogenesis (GO:0120163)
Regulation of Filopodium Assembly (GO:0051489)
Sphingolipid Biosynthetic Process (GO:0030148)
Ceramide Biosynthetic Process (GO:0046513)
Sterol Biosynthetic Process (GO:0016126)
Secondary Alcohol Biosynthetic Process (GO:1902653)
Cholesterol Biosynthetic Process (GO:0006695)
Response to Axon Injury (GO:0048678)
Pyrimidine Nucleobase Metabolic Process (GO:0006206)
Pyrimidine-Containing Compound Catabolic Process (GO:0072529)
Pyrimidine Nucleobase Catabolic Process (GO:0006208)
Galactosylceramide Metabolic Process (GO:0006681)
Nucleobase Catabolic Process (GO:0046113)
Glycosylceramide Biosynthetic Process (GO:0046476)
Galactosylceramide Biosynthetic Process (GO:0006682)
Galactolipid Biosynthetic Process (GO:0019375)
0 1000 2000 3000 4000
Enrichment Score
D
GO - Biological process
Depleted proteins at 2-month-old vs 8-month-old
Organic Anion Transport (GO:0015711)
Central Nervous System Development (GO:0007417)
Lipid Catabolic Process (GO:0016042)
Energy Derivation by Oxidation of Organic Compounds (GO:0015980)
Sphingolipid Catabolic Process (GO:0030149)
Pyridine Nucleotide Catabolic Process (GO:0019364)
NADH Metabolic Process (GO:0006734)
Fatty Acid Derivative Biosynthetic Process (GO:1901570)
Bicarbonate Transport (GO:0015701)
NAD Metabolic Process (GO:0019674)
Membrane Lipid Catabolic Process (GO:0046466)
0 500 1000 1500
Enrichment Score

**S3**

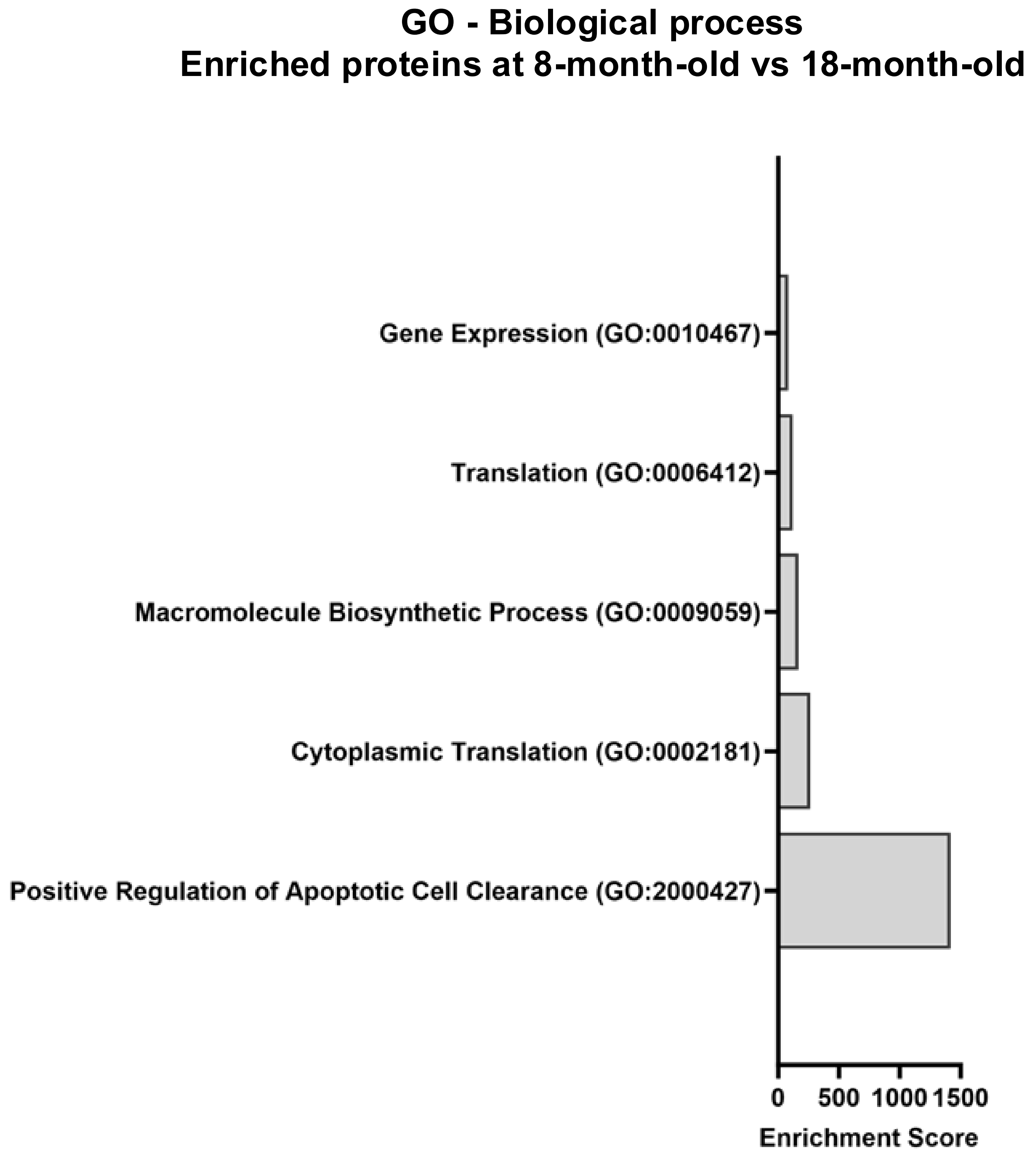
GO - Biological process
Enriched proteins at 8-month-old vs 18-month-old
Gene Expression (GO:0010467)
Translation (GO:0006412)
Macromolecule Biosynthetic Process (GO:0009059)
Cytoplasmic Translation (GO:0002181)
Positive Regulation of Apoptotic Cell Clearance (GO:2000427)
0
500
1000
1500
Enrichment Score

S 4

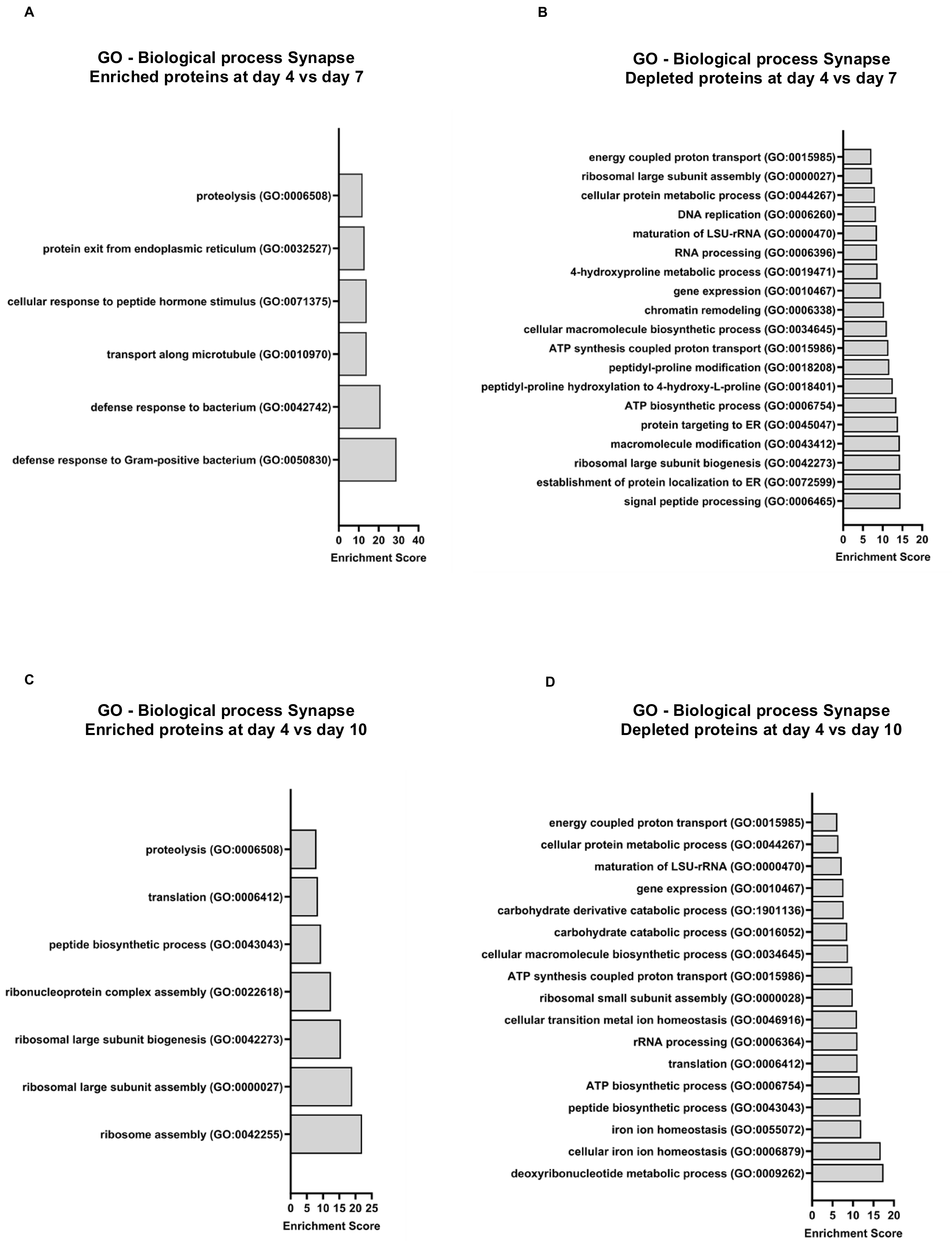
A
GO - Biological process Synapse
Enriched proteins at day 4 vs day 7
proteolysis (GO:0006508)
protein exit from endoplasmic reticulum (GO:0032527)
cellular response to peptide hormone stimulus (GO:0071375)
transport along microtubule (GO:0010970)
defense response to bacterium (GO:0042742)
defense response to Gram-positive bacterium (GO:0050830)
0 10 20 30 40
Enrichment Score
B
GO - Biological process Synapse
Depleted proteins at day 4 vs day 7
energy coupled proton transport (GO:0015985)
ribosomal large subunit assembly (GO:0000027)
cellular protein metabolic process (GO:0044267)
DNA replication (GO:0006260)
maturation of LSU-rRNA (GO:0000470)
RNA processing (GO:0006396)
4-hydroxyproline metabolic process (GO:0019471)
gene expression (GO:0010467)
chromatin remodeling (GO:0006338)
cellular macromolecule biosynthetic process (GO:0034645)
ATP synthesis coupled proton transport (GO:0015986)
peptidyl-proline modification (GO:0018208)
peptidyl-proline hydroxylation to 4-hydroxy-L-proline (GO:0018401)
ATP biosynthetic process (GO:0006754)
protein targeting to ER (GO:0045047)
macromolecule modification (GO:0043412)
ribosomal large subunit biogenesis (GO:0042273)
establishment of protein localization to ER (GO:0072599)
signal peptide processing (GO:0006465)
0 5 10 15 20
Enrichment Score
C
GO - Biological process Synapse
Enriched proteins at day 4 vs day 10
proteolysis (GO:0006508)
translation (GO:0006412)
peptide biosynthetic process (GO:0043043)
ribonucleoprotein complex assembly (GO:0022618)
ribosomal large subunit biogenesis (GO:0042273)
ribosomal large subunit assembly (GO:0000027)
ribosome assembly (GO:0042255)
0 5 10 15 20 25
Enrichment Score
D
GO - Biological process Synapse
Depleted proteins at day 4 vs day 10
energy coupled proton transport (GO:0015985)
cellular protein metabolic process (GO:0044267)
maturation of LSU-rRNA (GO:0000470)
gene expression (GO:0010467)
carbohydrate derivative catabolic process (GO:1901136)
carbohydrate catabolic process (GO:0016052)
cellular macromolecule biosynthetic process (GO:0034645)
ATP synthesis coupled proton transport (GO:0015986)
ribosomal small subunit assembly (GO:0000028)
cellular transition metal ion homeostasis (GO:0046916)
rRNA processing (GO:0006364)
translation (GO:0006412)
ATP biosynthetic process (GO:0006754)
peptide biosynthetic process (GO:0043043)
iron ion homeostasis (GO:0055072)
cellular iron ion homeostasis (GO:0006879)
deoxyribonucleotide metabolic process (GO:0009262)
0 5 10 15 20
Enrichment Score

S 5

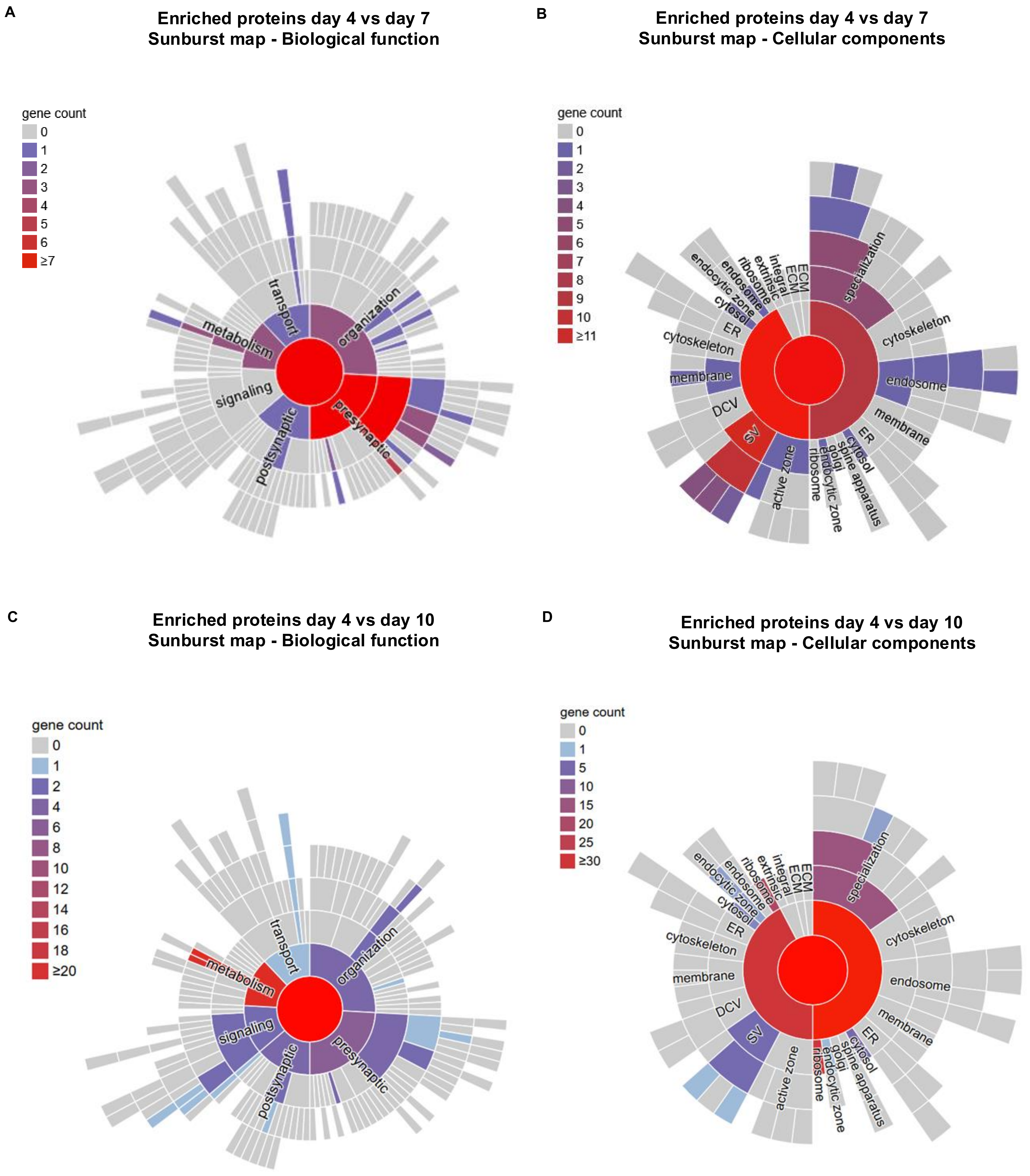
A
Enriched proteins day 4 vs day 7
Sunburst map - Biological function
gene count
0
1
2
3
4
5
6
≥7
transport
metabolism
organization
signaling
presynaptic
postsynaptic
B
Enriched proteins day 4 vs day 7
Sunburst map - Cellular components
gene count
0
1
2
3
4
5
6
7
8
9
10
≥11
ECM
ECM
integral
extrinsic
ribosome
endosome
endocytic zone
cytosol
ER
cytoskeleton
membrane
DCV
SV
active zone
ribosome
golgi
endocytic zone
spine apparatus
cytosol
ER
membrane
endosome
cytoskeleton
specialization
C
Enriched proteins day 4 vs day 10
Sunburst map - Biological function
gene count
0
1
2
4
6
8
10
12
14
16
18
≥20
transport
metabolism
organization
signaling
presynaptic
postsynaptic
D
Enriched proteins day 4 vs day 10
Sunburst map - Cellular components
gene count
0
1
5
10
15
20
25
≥30
ECM
ECM
integral
extrinsic
ribosome
endosome
endocytic zone
cytosol
ER
cytoskeleton
membrane
DCV
SV
active zone
ribosome
golgi
endocytic zone
spine apparatus
cytosol
ER
membrane
endosome
cytoskeleton
specialization

**S 6**

**Cortex - Cytoplasmic fraction**

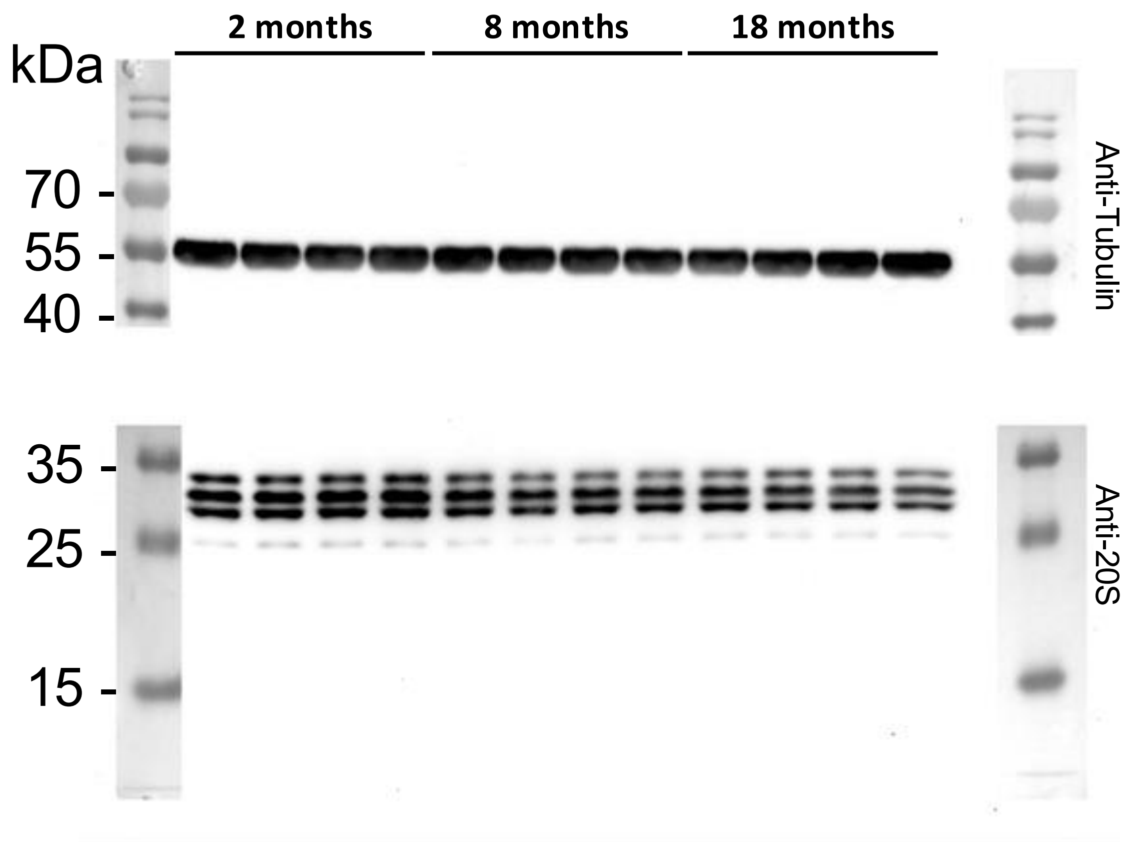


**Cortex - Synaptosomes**

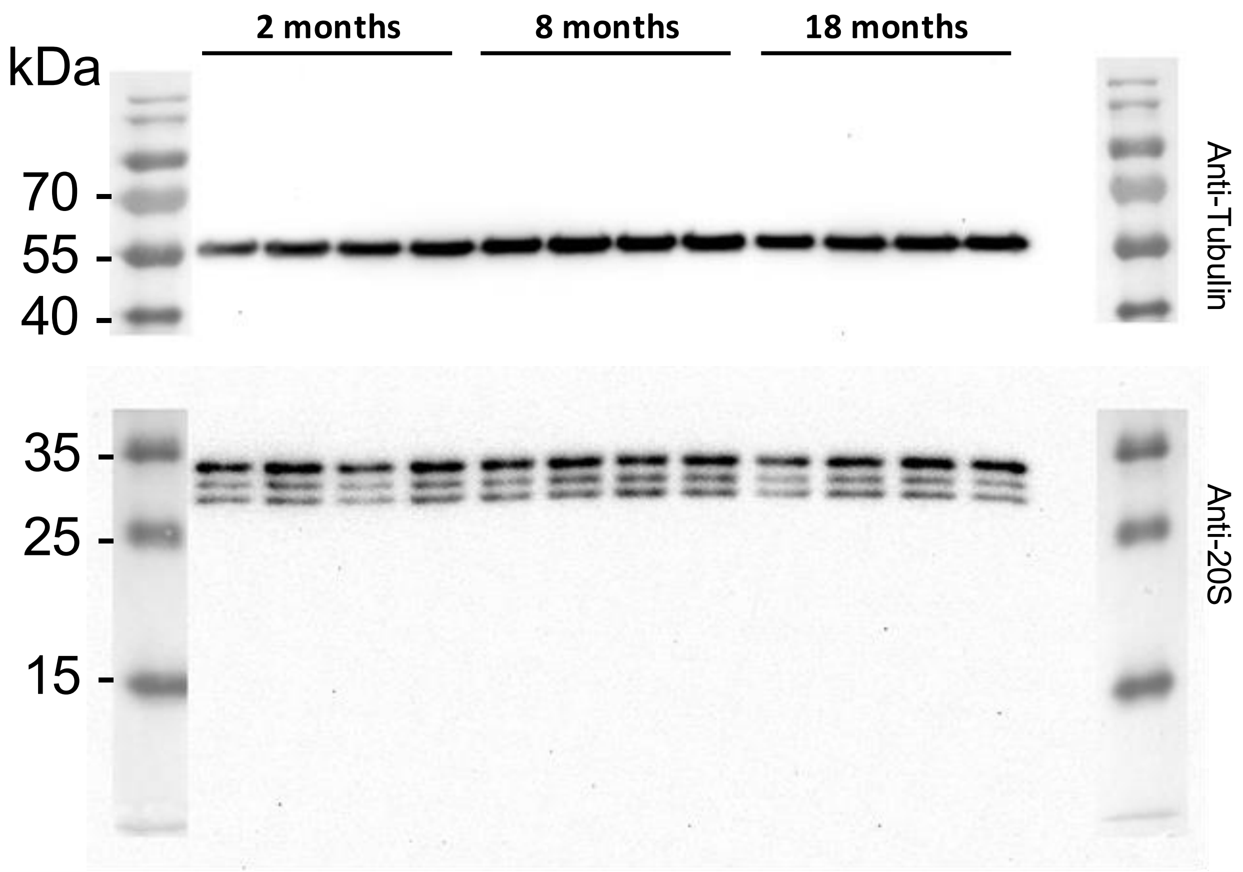


**Cerebellum - Cytoplasmic fraction**

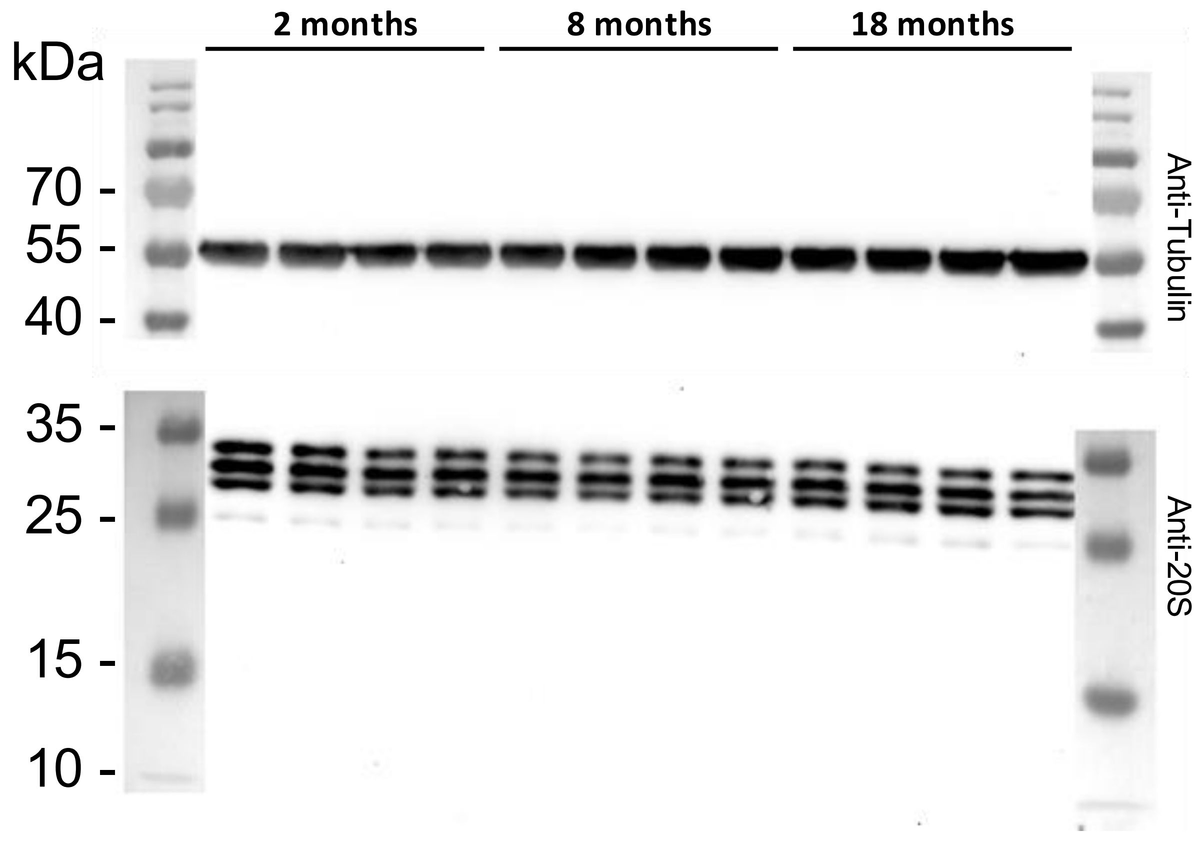


**Cerebellum - Synaptosomes**

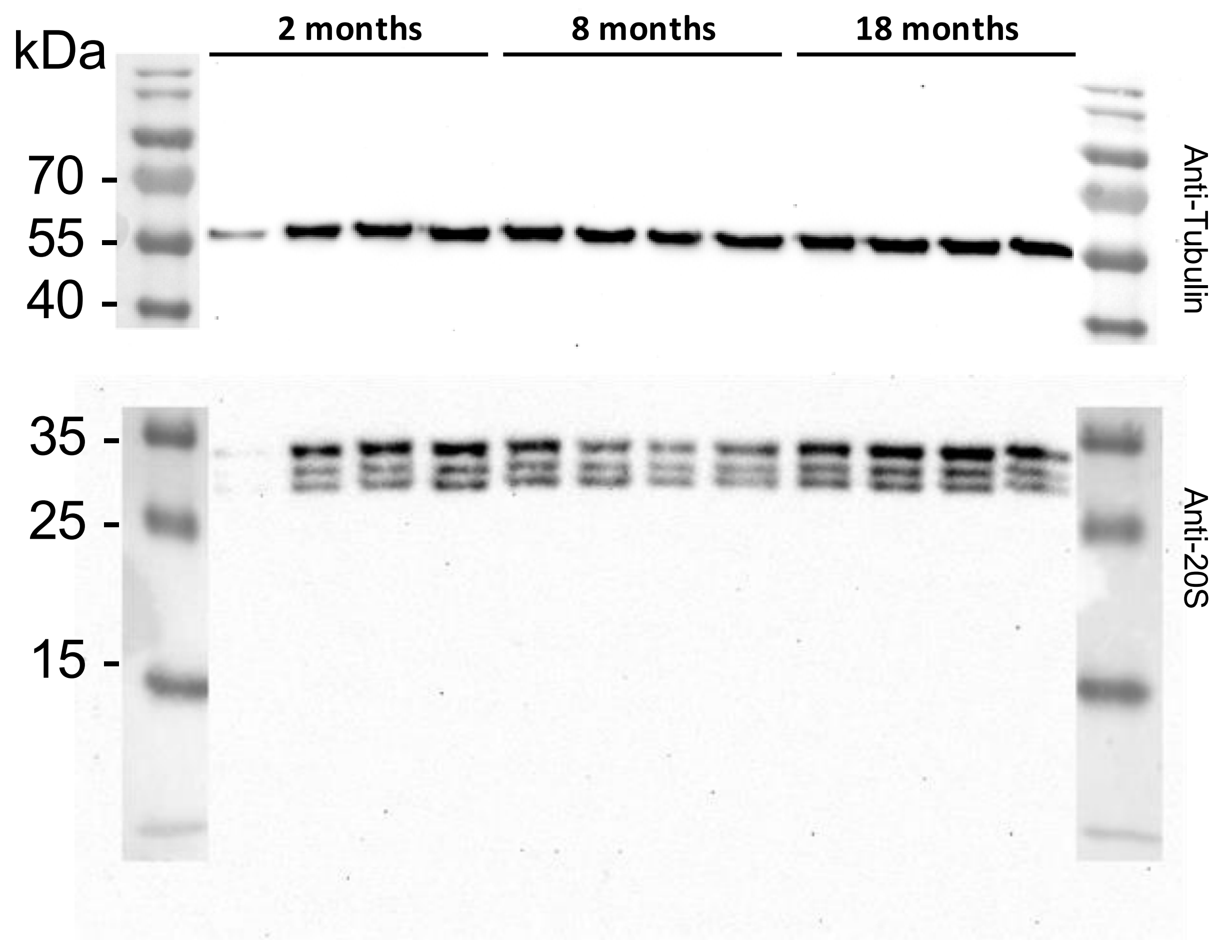

S 7

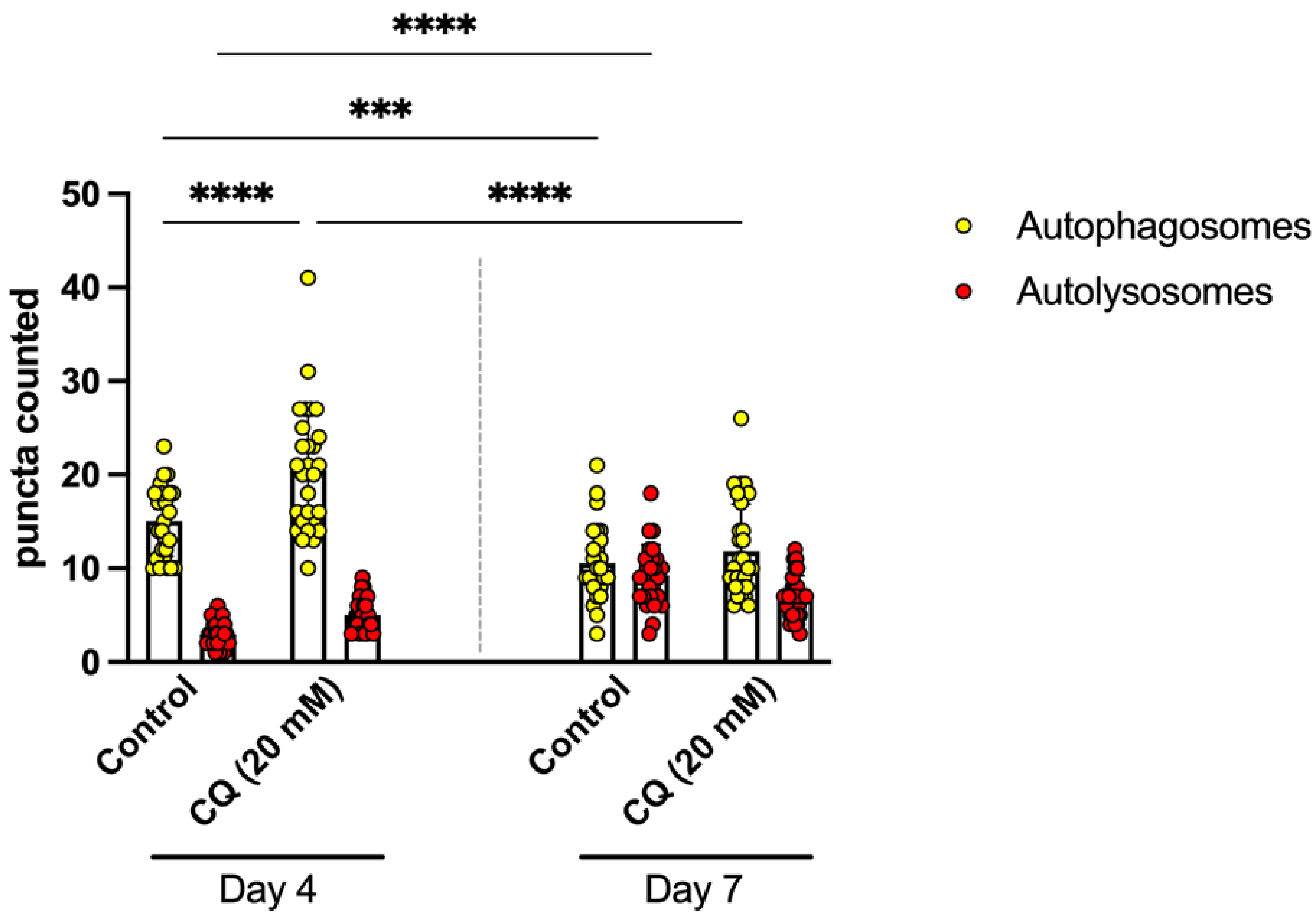

****
***
****
****
puncta counted
50
40
30
20
10
0
Control
CQ (20 mM)
Control
CQ (20 mM)
Day 4
Day 7
Autophagosomes
Autolysosomes

**Table S.1 Synaptic versus somatic candidate proteins with significant differential expression between subcompartments.**

Synaptic versus somatic candidate proteins with significant age-dependent differential expression in *C. elegans*. Listed are proteins that are significantly enriched (A) or depleted (B) in the synaptic fraction relative to the somatic fraction at day 4, compared with day 7 animals (Q < 0.05, fold change > 1.5). Protein ID, description, average $log_2$ ratio (synapse/soma), and adjusted Q value are shown for each candidate.

A. **Enriched in Day 4 and depleted in Day 7**

| Protein ID | Protein Description | AVG.Log2.Ratio | Q value |
|---|---|---|---|
| O44145 | PERMeable eggshell | 4.22 | 3.3E-02 |
| SUN1 | Sun domain-containing protein 1 | 3.53 | 4.6E-03 |
| RM35 | Probable 39S ribosomal protein L35, mitochondrial | 3.48 | 2.7E-03 |
| SNG1 | Synaptogyrin homolog 1 | 3.15 | 5.7E-03 |
| UNC17 | Vesicular acetylcholine transporter unc-17 | 2.80 | 2.9E-05 |
| RL5 | 60S ribosomal protein L5 | 2.51 | 3.0E-07 |
| G5EGI7 | 2 (Zwei) IG domain protein | 2.49 | 9.4E-03 |
| SYT1 | Synaptotagmin-1;SyNapTotagmin | 2.48 | 6.5E-16 |
| Q21925 | ProstaGlandin E Synthase homolog | 2.40 | 9.0E-09 |

| | | | |
|---|---|---|---|
| CAN5 | Calpain-5 | 2.32 | 4.6E-02 |
| RL31 | 60S ribosomal protein L31 | 2.14 | 5.0E-04 |
| NDUB7 | NADH dehydrogenase [ubiquinone] 1 beta subcomplex subunit 7 | 2.09 | 3.5E-02 |
| SC5A7 | High-affinity choline transporter 1 | 2.04 | 1.5E-04 |
| RL6 | 60S ribosomal protein L6 | 2.03 | 1.4E-07 |
| DYL1 | Dynein light chain 1, cytoplasmic | 1.96 | 4.4E-02 |
| Q86S81 | ELRR (Extracellular Leucine-Rich Repeat) ONly | 1.92 | 3.5E-02 |
| Q86NE0 | ASpartyl Protease | 1.91 | 2.4E-02 |
| O16619 | DeHydrogenases, Short chain | 1.86 | 4.3E-10 |
| EAA1 | Excitatory amino acid transporter | 1.74 | 4.4E-02 |
| UNC9 | Innexin unc-9 | 1.74 | 2.0E-03 |
| CATA1 | Peroxisomal catalase 1 | 1.73 | 1.0E-03 |
| Q9XWE1 | DNaJ domain (Prokaryotic heat shock protein) | 1.72 | 1.3E-03 |
| APOP1 | APOPT family protein Y39B6A.34, mitochondrial | 1.70 | 1.7E-02 |

| | | | |
|---|---|---|---|
| Q9GYJ9 | Sorting NeXin | 1.69 | 2.8E-02 |
| Q95YD8 | Isocitrate dehydrogenase [NAD] subunit, mitochondrial | 1.69 | 9.2E-06 |
| RL15 | 60S ribosomal protein L15 | 1.67 | 6.2E-04 |
| Q20992 | Acyl-coenzyme A oxidase | 1.67 | 1.8E-03 |
| Q22800 | NADH Ubiquinone oxidoreductase Fe-S protein | 1.67 | 3.0E-05 |
| H2KZ88 | UDP-GlucuronosylTransferase | 1.64 | 1.9E-04 |
| GLYC | Serine hydroxymethyltransferase | 1.61 | 8.6E-05 |
| PISD | Phosphatidylserine decarboxylase proenzyme, mitochondrial | 1.57 | 4.6E-02 |
| P91493 | SyNapTotagmin | 1.56 | 1.2E-03 |
| SUMO | Small ubiquitin-related modifier | 1.55 | 5.0E-05 |
| Q9U3Q0 | Mitochondrial Ribosomal Protein, Small | 1.55 | 5.2E-03 |
| CHSSB | Chondroitin sulfate synthase mig-22 | 1.54 | 2.1E-02 |
| SUV3 | ATP-dependent RNA helicase SUV3 homolog, mitochondrial | 1.54 | 3.4E-03 |

| | | | |
|---|---|---|---|
| O16521 | NADH-cytochrome b5 reductase | 1.54 | 3.3E-02 |
| O01869 | Ribosomal Protein, Small subunit | 1.53 | 1.7E-06 |
| Q9N4P2 | GLutaRedoXin | 1.51 | 4.7E-03 |
| Q95YB2 | Acyl CoA DeHydrogenase | 1.51 | 7.1E-11 |
| Q21083 | Mitochondrial Ribosomal Protein, Large | 1.51 | 8.2E-10 |
| OXDD2 | D-aspartate oxidase 2 | 1.50 | 2.5E-05 |
| Q93896 | MonoCarboxylate Transporter family | 1.50 | 5.6E-04 |
| O16922 | UDP-GlucuronosylTransferase | 1.47 | 8.9E-03 |
| A5PEY3 | DNA-directed RNA polymerase | 1.45 | 1.2E-03 |
| O17694 | RNAi-Induced Longevity | 1.43 | 4.9E-02 |
| RT24 | 28S ribosomal protein S24, mitochondrial | 1.41 | 4.2E-04 |
| Q21154 | MICOS complex subunit | 1.41 | 2.9E-03 |
| A0A0K3ARZ7 | F-box A protein | 1.40 | 3.7E-02 |
| STX1A | Syntaxin-1A homolog | 1.40 | 2.9E-03 |

| | | | |
|---|---|---|---|
| KMO | Kynurenine 3-monooxygenase | 1.40 | 1.1E-07 |
| G5EEG6 | Carrier protein (C2) | 1.40 | 2.1E-03 |
| Q95R12 | ACyLtransferase-like | 1.40 | 1.5E-03 |
| Q95X39 | Mitochondrial Ribosomal Protein, Large | 1.39 | 1.2E-03 |
| NNRD | ATP-dependent (S)-NAD(P)H-hydrate dehydratase | 1.38 | 3.8E-02 |
| Q95PZ8 | Asparaginyl(N) Amino-acyl tRNA Synthetase | 1.38 | 1.3E-03 |
| O02147 | Carboxylic ester hydrolase | 1.36 | 4.3E-02 |
| ALH13 | Probable delta-1-pyrroline-5-carboxylate synthase | 1.35 | 1.2E-14 |
| DHTK1 | Probable 2-oxoglutarate dehydrogenase E1 component DHKTD1 homolog, mitochondrial | 1.35 | 6.5E-04 |
| Q9TZ90 | Mitochondrial Ribosomal Protein, Large | 1.35 | 2.9E-02 |
| O17643 | Isocitrate dehydrogenase [NADP] | 1.35 | 2.0E-03 |
| B2MZD0 | Vacuolar H ATPase;V-type proton ATPase subunit a | 1.34 | 1.9E-02 |

| | | | |
|---|---|---|---|
| Q18660 | Fatty Acid CoA Synthetase family | 1.34 | 5.0E-05 |
| Q22968 | Aminomethyltransferase | 1.33 | 3.2E-03 |
| I2HA98 | PhenylAlanine Hydroxylase;Probable phenylalanine-4-hydroxylase 1 | 1.29 | 8.2E-08 |
| G5EFC8 | Mammalian cell Death Associated Protein related | 1.28 | 4.3E-05 |
| Q23066 | Carnitine Palmitoyl Transferase | 1.27 | 4.6E-03 |
| G8JY83 | UBiQuitin;Polyubiquitin-A;Ubiquitin-60S ribosomal protein L40 | 1.27 | 1.2E-06 |
| G5EDB6 | ParaPleGiN AAA protease family | 1.27 | 1.9E-03 |
| AP2M | AP-2 complex subunit mu | 1.27 | 2.6E-05 |
| G5EG62 | UNC-45 | 1.26 | 2.9E-02 |
| Q23295 | Mitochondrial Processing Peptidase Beta | 1.26 | 1.8E-05 |
| ANI2 | Anillin-like protein 2 | 1.25 | 6.4E-03 |
| Q7KPW7 | Mitochondrial Ribosomal Protein, Large | 1.25 | 9.3E-06 |
| Q21021 | Nuclear Pore complex Protein | 1.24 | 4.6E-02 |

| | | | |
|---|---|---|---|
| Q17403 | UDP-GlucuronosylTransferase | 1.24 | 1.2E-02 |
| O01795 | DUal OXidase;Dual oxidase 1 | 1.24 | 2.4E-02 |
| CLPP1 | ATP-dependent Clp protease proteolytic subunit 1, mitochondrial | 1.24 | 2.2E-02 |
| ATPK | Putative ATP synthase subunit f, mitochondrial | 1.23 | 9.4E-04 |
| GSTK2 | Glutathione s-transferase kappa 2 | 1.22 | 2.7E-02 |
| D6R8W9 | RaBConnectin related | 1.21 | 1.9E-03 |
| Q20600 | TBC (Tre-2/Bub2/Cdc16) domain family | 1.21 | 2.1E-02 |
| RT05 | Putative 28S ribosomal protein S5, mitochondrial | 1.21 | 6.0E-05 |
| LACB2 | Beta-lactamase-like protein 2 homolog | 1.19 | 1.3E-03 |
| RS16 | 40S ribosomal protein S16 | 1.19 | 6.2E-06 |
| D9N122 | Eps15 (Endocytosis protein) Homologous Sequence;EHS-1 | 1.18 | 6.7E-03 |
| NDUA5 | Probable NADH dehydrogenase [ubiquinone] 1 alpha subcomplex subunit 5 | 1.18 | 1.1E-02 |

| | | | |
|---|---|---|---|
| SIP1 | Stress-induced protein 1 | 1.18 | 2.7E-03 |
| AT1B1 | Sodium/potassium-transporting ATPase subunit beta-1 | 1.18 | 4.0E-05 |
| ATP5E | Putative ATP synthase subunit epsilon, mitochondrial | 1.18 | 1.5E-02 |
| GSK3 | Glycogen synthase kinase-3 | 1.17 | 1.3E-02 |
| Q8MXI1 | Ubiquinol-Cytochrome c oxidoReductase complex | 1.16 | 1.3E-02 |
| NDX7 | Putative nudix hydrolase 7 | 1.16 | 9.4E-04 |
| Q19813 | COLlagen | 1.14 | 4.8E-03 |
| CIA30 | Probable complex I intermediate-associated protein 30, mitochondrial | 1.14 | 3.0E-03 |
| RL13 | 60S ribosomal protein L13 | 1.13 | 3.9E-07 |
| O44727 | CalPoNin | 1.13 | 1.3E-03 |
| CRI3 | Conserved regulator of innate immunity protein 3 | 1.13 | 6.4E-03 |
| Q95XT5 | TRanslocon-Associated Protein | 1.13 | 3.6E-03 |
| COQ8 | Atypical kinase coq-8, mitochondrial | 1.12 | 7.4E-04 |

| | | | |
|---|---|---|---|
| UNC89 | Muscle M-line assembly protein unc-89 | 1.11 | 1.6E-02 |
| A9QY30 | Mitochondrial genome maintenance exonuclease 1 | 1.11 | 2.0E-03 |
| Q965T4 | Translational Activator of Cytochrome c Oxidase | 1.11 | 3.6E-05 |
| TIM50 | Mitochondrial import inner membrane translocase subunit TIM50 | 1.10 | 3.5E-06 |
| Q22470 | Mitochondrial Ribosomal Protein, Small | 1.09 | 8.3E-04 |
| MTX1 | Metaxin-1 homolog | 1.09 | 3.0E-02 |
| Q94056 | O-ACyltransferase homolog | 1.09 | 3.3E-02 |
| Q9XUB7 | Fatty Acid/Retinol binding protein | 1.08 | 2.1E-03 |
| ADF2 | Actin-depolymerizing factor 2, isoform c | 1.08 | 1.9E-03 |
| O18180 | Mitochondrial Ribosomal Protein, Large | 1.07 | 4.3E-04 |
| RM18 | 39S ribosomal protein L18, mitochondrial | 1.07 | 3.4E-02 |
| GPDM | Probable glycerol-3-phosphate dehydrogenase, mitochondrial | 1.07 | 2.9E-05 |

| | | | |
|---|---|---|---|
| GSTK1 | Glutathione S-transferase kappa 1 | 1.06 | 1.7E-02 |
| RS9 | 40S ribosomal protein S9 | 1.06 | 3.6E-06 |
| G5EFU2 | Adenine Nucleotide Translocator | 1.05 | 4.6E-03 |
| HGD | Homogentisate 1,2-dioxygenase | 1.05 | 4.8E-02 |
| EFGM | Elongation factor G, mitochondrial | 1.05 | 1.4E-09 |
| Q95XN2 | Mitochondrial Processing Peptidase Alpha | 1.05 | 5.5E-03 |
| Q20122 | Acyl carrier protein | 1.04 | 9.0E-04 |
| FABP3 | Fatty acid-binding protein homolog 3;Lipid Binding Protein | 1.04 | 3.2E-02 |
| Q95Y25 | Glutamyl-tRNA(Gln) amidotransferase subunit A, mitochondrial;Glutamyl-tRNA(Gln) amidotransferase subunit A, mitochondrial | 1.04 | 3.0E-02 |
| B1Q250 | UDP-glucuronosyltransferase | 1.04 | 1.7E-03 |
| Q7K742 | Daf-16-Dependent Longevity (WT but not daf-16 lifespan increased) | 1.04 | 1.9E-05 |
| Q6A575 | Mitochondrial Ribosomal Protein, Large | 1.04 | 1.7E-03 |

| | | | |
|---|---|---|---|
| COX5A | Cytochrome c oxidase subunit 5A, mitochondrial | 1.03 | 6.8E-04 |
| UNC18 | Putative acetylcholine regulator unc-18 | 1.02 | 2.4E-02 |
| SAM50 | SAM50-like protein gop-3 | 1.01 | 2.5E-03 |
| Q20203 | Calsequestrin | 1.01 | 2.0E-04 |
| DEOC | Putative deoxyribose-phosphate aldolase | 1.01 | 3.5E-02 |
| COX1 | Cytochrome c oxidase subunit 1 | 1.00 | 3.3E-06 |
| Q17447 | Fatty Acid Amide Hydrolase homolog | 1.00 | 2.7E-02 |
| Q95Y73 | Mitochondrial elongation factor G2 | 0.98 | 2.9E-02 |
| A0A0K3ATC0 | Tyramine Beta Hydroxylase;Tyramine beta-hydroxylase | 0.98 | 2.9E-02 |
| O17983 | ARRestin Domain protein | 0.98 | 1.1E-03 |
| Q9UA63 | Mitochondrial Ribosomal Protein, Large | 0.98 | 8.0E-05 |
| Q9NF11 | Hypoxanthine PhosphoRibosylTransferase homolog | 0.98 | 6.1E-04 |
| G5EBH7 | CALUmenin (Calcium-binding protein) homolog | 0.97 | 8.3E-03 |

| | | | |
|---|---|---|---|
| HCDH2 | Probable 3-hydroxyacyl-CoA dehydrogenase B0272.3 | 0.97 | 4.9E-03 |
| RT14 | Probable 40S ribosomal protein S14, mitochondrial | 0.96 | 3.6E-03 |
| Q9TZ33 | Ubiquinol-Cytochrome c oxidoReductase complex | 0.96 | 1.8E-05 |
| THILH | Acetyl-CoA acetyltransferase homolog, mitochondrial | 0.96 | 7.7E-05 |
| VPP1 | Probable V-type proton ATPase 116 kDa subunit a | 0.96 | 1.7E-08 |
| H2KYN3 | Acyl CoA DeHydrogenase | 0.96 | 2.5E-09 |
| H2KZF5 | Phosphoethanolamine MethylTransferase | 0.96 | 5.1E-05 |
| Q18697 | Phosphate transporter | 0.96 | 3.3E-02 |
| Q21569 | Beta-LACTamase domain containing | 0.95 | 4.3E-03 |
| ETFD | Electron transfer flavoprotein-ubiquinone oxidoreductase, mitochondrial | 0.95 | 1.6E-07 |
| FOLT2 | Folate-like transporter 2 | 0.95 | 3.8E-03 |
| TAM41 | Phosphatidate cytidylyltransferase, mitochondrial | 0.95 | 1.0E-04 |

| | | | |
|---|---|---|---|
| NLTP1 | Non-specific lipid-transfer protein-like 1 | 0.95 | 3.2E-02 |
| G4SRS5 | ABC Transporter, Mitochondrial | 0.95 | 2.4E-06 |
| Q9U1X0 | NADPH:adrenodoxin oxidoreductase, mitochondrial | 0.94 | 1.6E-02 |
| ACOX5 | Probable peroxisomal acyl-coenzyme A oxidase 5 | 0.94 | 6.1E-04 |
| SFXN5 | Sideroflexin-5 | 0.93 | 5.7E-04 |
| RT17 | 28S ribosomal protein S17, mitochondrial | 0.93 | 2.5E-02 |
| LONM | Lon protease homolog, mitochondrial | 0.93 | 1.0E-06 |
| MECR1 | Probable trans-2-enoyl-CoA reductase 1, mitochondrial | 0.93 | 1.2E-03 |
| O02207 | OXA mitochondrial inner membrane insertase homolog | 0.93 | 2.4E-05 |
| Q22251 | Cytochrome OXidase assembly protein | 0.92 | 1.1E-02 |
| UNC33 | Protein unc-33;Uncharacterized protein | 0.92 | 3.5E-03 |
| Q9BL86 | Mitochondrial Ribosomal Protein, Large | 0.91 | 6.0E-09 |

| | | | |
|---|---|---|---|
| SDHB | Succinate dehydrogenase [ubiquinone] iron-sulfur subunit, mitochondrial | 0.91 | 1.1E-04 |
| O44954 | Succinate dehydrogenase [ubiquinone] flavoprotein subunit, mitochondrial | 0.89 | 1.0E-07 |
| COQ5 | 2-methoxy-6-polyprenyl-1,4-benzoquinol methylase, mitochondrial | 0.88 | 3.5E-03 |
| G5ED08 | Stomatin-like protein UNC-1;Protein unc-1 | 0.88 | 2.8E-02 |
| Q9U2Q8 | Peptidyl-prolyl cis-trans isomerase | 0.88 | 1.3E-03 |
| DPYD | Dihydropyrimidine dehydrogenase [NADP(+)] | 0.87 | 1.3E-03 |
| MTX2 | Metaxin-2 homolog | 0.87 | 1.6E-03 |
| G5ECG7 | Defecation Suppressor of Clk-1 | 0.87 | 5.2E-05 |
| Q9NEI6 | Mitochondrial Ribosomal Protein, Small | 0.87 | 2.7E-02 |
| H2KYJ2 | TransThyretin-Related family domain | 0.86 | 5.9E-03 |
| RT23 | Probable 28S ribosomal protein S23, mitochondrial | 0.86 | 2.9E-03 |
| Q9GYI2 | Mitochondrial Ribosomal Protein, Small | 0.86 | 5.7E-03 |

| | | | |
|---|---|---|---|
| RM49 | Probable 39S ribosomal protein L49, mitochondrial | 0.85 | 1.3E-04 |
| G5EDJ3 | Equilibrative Nucleoside Transporter | 0.85 | 4.6E-03 |
| Q9U2S4 | ADP-Ribosylation Factor related | 0.84 | 8.5E-03 |
| Q8MPW0 | DEgenerin Like | 0.84 | 1.9E-05 |
| Q9XX18 | Mitochondrial Ribosomal Protein, Large | 0.84 | 2.3E-03 |
| Q9U295 | Carboxylic ester hydrolase | 0.84 | 1.7E-04 |
| Q18853 | CYtochrome C | 0.83 | 1.5E-05 |
| RT25 | Probable 28S ribosomal protein S25, mitochondrial | 0.83 | 3.6E-03 |
| G5ECI8 | Mitochondrial phenylalanyl-tRNA synthetase;Phenylalanyl Amino-acyl tRNA Synthetase | 0.83 | 1.9E-04 |
| O45148 | DihydroLipoamide S-SuccinylTransferase | 0.82 | 4.5E-02 |
| O44887 | Innexin | 0.82 | 5.8E-03 |
| LIAS | Lipoyl synthase, mitochondrial | 0.82 | 1.9E-02 |
| Q9XX28 | DeHydrogenases, Short chain | 0.82 | 8.0E-03 |

| | | | |
|---|---|---|---|
| Q4W5T0 | Mitochondrial Ribosomal Protein, Large | 0.81 | 3.6E-03 |
| Q9U348 | COLlagen | 0.81 | 7.8E-03 |
| Q9XXN2 | MICOS complex subunit MIC60 | 0.81 | 6.9E-12 |
| RL9 | 60S ribosomal protein L9 | 0.81 | 1.1E-02 |
| A0A061AKV1 | UDP-glucuronosyltransferase | 0.80 | 6.2E-04 |
| Q9N3T5 | SPG (Spastic paraplegia) | 0.80 | 6.1E-03 |
| Q94055 | Serine--pyruvate aminotransferase | 0.80 | 1.1E-06 |
| PAT2 | Integrin alpha pat-2 | 0.79 | 1.4E-02 |
| SODC | Superoxide dismutase [Cu-Zn] | 0.79 | 6.7E-03 |
| O44619 | Carnitine Palmitoyl Transferase | 0.78 | 1.1E-06 |
| O61818 | Mitochondrial Ribosomal Protein, Small | 0.78 | 1.3E-06 |
| DCXR | L-xylulose reductase | 0.78 | 1.5E-02 |
| DIF1 | Protein dif-1 | 0.78 | 3.5E-02 |
| LRP | Low-density lipoprotein receptor-related protein | 0.77 | 9.9E-05 |

| | | | |
|---|---|---|---|
| G5ED31 | Ubiquinol-Cytochrome c oxidoReductase complex | 0.77 | 1.0E-03 |
| ODB2 | Lipoamide acyltransferase component of branched-chain alpha-keto acid dehydrogenase complex, mitochondrial | 0.77 | 3.6E-02 |
| RM41 | 39S ribosomal protein L41, mitochondrial | 0.77 | 2.7E-03 |
| TBCB | Tubulin-specific chaperone B | 0.76 | 2.7E-02 |
| Q9XWD1 | Fatty Acid CoA Synthetase family | 0.75 | 2.5E-03 |
| G5ECD0 | CYtochrome P450 family | 0.75 | 2.8E-03 |
| DNJ10 | DnaJ homolog dnj-10 | 0.75 | 1.7E-07 |
| O62107 | UDP-GALactose 4-Epimerase | 0.73 | 1.4E-02 |
| ACD11 | Acyl-CoA dehydrogenase family member 11 | 0.73 | 2.5E-06 |
| CYSK2 | Bifunctional L-3-cyanoalanine synthase/cysteine synthase | 0.72 | 3.8E-03 |
| Q9XTU9 | Glutaredoxin | 0.72 | 3.3E-03 |
| RT09 | Probable 40S ribosomal protein S9, mitochondrial | 0.72 | 4.0E-05 |

| | | | |
|---|---|---|---|
| Q9NAG2 | Mitochondrial Ribosomal Protein, Large | 0.72 | 8.8E-04 |
| G5EDQ0 | Chloride channel protein | 0.72 | 4.8E-02 |
| MFRN | Mitoferrin | 0.72 | 2.5E-02 |
| Q19130 | Glutamine-Fructose 6-phosphate AminoTransferase homolog | 0.71 | 7.3E-03 |
| MPC1 | Probable mitochondrial pyruvate carrier 1 | 0.70 | 2.0E-02 |
| NDUS7 | Probable NADH dehydrogenase [ubiquinone] iron-sulfur protein 7, mitochondrial | 0.70 | 7.7E-04 |
| CPR5 | Cathepsin B-like cysteine proteinase 5 | 0.69 | 2.3E-02 |
| UGT47 | Putative UDP-glucuronosyltransferase ugt-47 | 0.69 | 1.5E-04 |
| AGT2L | Alanine--glyoxylate aminotransferase 2-like | 0.69 | 2.6E-02 |
| Q9TYJ8 | Mitochondrial Ribosomal Protein, Large | 0.69 | 2.8E-02 |
| RM45 | Probable 39S ribosomal protein L45, mitochondrial | 0.68 | 2.5E-04 |
| O17612 | Enoyl-CoA Hydratase | 0.68 | 2.5E-03 |

| | | | |
|---|---|---|---|
| INX12 | Innexin-12 | 0.68 | 1.3E-02 |
| ODP2 | Dihydrolipoyllysine-residue acetyltransferase component of pyruvate dehydrogenase complex, mitochondrial | 0.67 | 9.1E-03 |
| RT15 | 28S ribosomal protein S15, mitochondrial | 0.67 | 5.2E-05 |
| Q95PY4 | Mitochondrial Ribosomal Protein, Small | 0.66 | 1.3E-05 |
| RM51 | 39S ribosomal protein L51, mitochondrial | 0.66 | 1.0E-02 |
| TPP2 | Tripeptidyl-peptidase 2 | 0.66 | 6.8E-07 |
| GCP | Bifunctional glyoxylate cycle protein | 0.65 | 3.4E-12 |
| P4HA2 | Prolyl 4-hydroxylase subunit alpha-2 | 0.65 | 4.3E-03 |
| O17805 | COLlagen | 0.64 | 4.0E-02 |
| HACD | Very-long-chain (3R)-3-hydroxyacyl-CoA dehydratase hpo-8 | 0.64 | 2.5E-02 |
| GRPE | GrpE protein homolog, mitochondrial | 0.64 | 2.3E-02 |
| Q9BPN6 | Mitochondrial Ribosomal Protein, Large | 0.62 | 2.2E-06 |

| | | | |
|---|---|---|---|
| RS4 | 40S ribosomal protein S4 | 0.62 | 1.6E-12 |
| O18000 | Patterned Expression Site | 0.62 | 2.3E-02 |
| UCR1 | Cytochrome b-c1 complex subunit 1, mitochondrial | 0.62 | 7.1E-03 |
| P91306 | C. Elegans Y-box | 0.61 | 4.3E-02 |
| ADAS | Alkyldihydroxyacetonephosphate synthase | 0.61 | 3.3E-02 |
| Q22341 | TransThyretin-Related family domain | 0.61 | 3.5E-02 |
| COX2 | Cytochrome c oxidase subunit 2 | 0.60 | 6.4E-03 |
| G5EG54 | Carnitine Palmitoyl Transferase | 0.60 | 2.6E-02 |
| RT31 | 28S ribosomal protein S31, mitochondrial | 0.59 | 1.0E-04 |
| Q9GZD1 | UDP-GlucuronosylTransferase | 0.59 | 8.5E-03 |
| U562 | UPF0562 protein C29E4.12 | 0.59 | 3.2E-02 |
| Q20062 | Mitochondrial Associated RiBonuclease homolog | 0.59 | 2.7E-03 |
| BST14 | Bestrophin homolog 14 | 0.58 | 1.5E-02 |

### B. Enriched in Day 7 and depleted in Day 4

| Protein ID | Protein Description | AVG.Log2.Ratio | Q value |
|---|---|---|---|
| P91350 | Nuclear Pore complex Protein | -0.58 | 1.0E-02 |
| RCC1 | Regulator of chromosome condensation | -0.58 | 7.4E-03 |
| CAN | Calpain clp-1 | -0.58 | 7.5E-03 |
| RL17 | 60S ribosomal protein L17 | -0.58 | 6.4E-05 |
| PROF1 | Profilin-1 | -0.59 | 4.0E-06 |
| Q965S5 | SyNapTotagmin | -0.59 | 3.8E-02 |
| RL38 | 60S ribosomal protein L38 | -0.59 | 1.4E-02 |
| G5ED36 | Endocytosis protein RME-8;Receptor Mediated Endocytosis | -0.59 | 9.8E-03 |
| O17766 | Lipase | -0.59 | 5.7E-03 |
| ILKH | Integrin-linked protein kinase homolog pat-4 | -0.60 | 5.0E-03 |
| Q9GPA0 | UDP-Glucose Glycoprotein glucosylTransferase | -0.60 | 1.1E-22 |
| CYP5 | Peptidyl-prolyl cis-trans isomerase 5 | -0.60 | 2.6E-03 |

| | | | |
|---|---|---|---|
| GBB1 | Guanine nucleotide-binding protein subunit beta-1 | -0.61 | 3.6E-03 |
| Q9XVS9 | DeHydrogenases, Short chain | -0.62 | 2.2E-02 |
| RLA0 | 60S acidic ribosomal protein P0 | -0.62 | 7.5E-12 |
| SYRC | Probable arginine--tRNA ligase, cytoplasmic;Arginyl(R) Amino-acyl tRNA Synthetase | -0.62 | 1.6E-03 |
| NDX6 | Putative nudix hydrolase 6 | -0.62 | 3.7E-02 |
| RS26 | 40S ribosomal protein S26 | -0.62 | 2.2E-04 |
| SURF4 | Surfeit locus protein 4 homolog | -0.63 | 4.8E-02 |
| Q94230 | Pur alpha Like Protein | -0.63 | 1.2E-07 |
| RL13A | 60S ribosomal protein L13a | -0.64 | 9.8E-07 |
| Q9U2Z1 | Yeast SEC homolog | -0.64 | 2.9E-02 |
| Q5TKA3 | Fatty Acid CoA Synthetase family | -0.64 | 1.7E-03 |
| U520 | Putative U5 small nuclear ribonucleoprotein 200 kDa helicase | -0.64 | 8.3E-11 |
| PSMD3 | 26S proteasome non-ATPase regulatory subunit 3 | -0.65 | 6.4E-04 |

| | | | |
|---|---|---|---|
| RPN2 | Dolichyl-diphosphooligosaccharide--protein glycosyltransferase subunit 2 | -0.65 | 1.4E-04 |
| IMP2 | Intramembrane protease 2 | -0.65 | 1.1E-08 |
| Q19371 | Yeast SEC homolog | -0.66 | 1.0E-02 |
| KINH | Kinesin heavy chain | -0.66 | 1.3E-06 |
| YVRI | GILT-like protein F37H8.5 | -0.67 | 1.2E-04 |
| Q9GZH5 | Proteasome Regulatory Particle, Non-ATPase-like | -0.67 | 3.2E-05 |
| O61792 | Proteasome Regulatory Particle, Non-ATPase-like | -0.67 | 8.0E-03 |
| RS13 | 40S ribosomal protein S13 | -0.67 | 4.1E-06 |
| GYS | Glycogen [starch] synthase | -0.67 | 6.7E-03 |
| Q18231 | Ribosomal Protein, Small subunit | -0.67 | 1.0E-03 |
| PHB2 | Mitochondrial prohibitin complex protein 2 | -0.68 | 2.6E-02 |
| VIP1 | Inositol hexakisphosphate and diphosphoinositol-pentakisphosphate kinase | -0.68 | 2.9E-03 |

| | | | |
|---|---|---|---|
| H1UBL1 | MItochondrial Calcium Uptake protein;Calcium uptake protein 1 homolog | -0.69 | 4.0E-02 |
| RL23 | 60S ribosomal protein L23 | -0.69 | 7.4E-06 |
| P91495 | Nuclear Pore complex Protein | -0.69 | 1.1E-20 |
| G3MU79 | Ubiquitin carboxyl-terminal hydrolase 7 | -0.69 | 1.2E-03 |
| RS15 | 40S ribosomal protein S15 | -0.70 | 9.7E-06 |
| MOGS1 | Mannosyl-oligosaccharide glucosidase | -0.70 | 1.1E-03 |
| NOC2L | Nucleolar complex protein 2 homolog | -0.71 | 2.9E-03 |
| Q23378 | TransThyretin-Related family domain | -0.71 | 8.8E-03 |
| Q9XWH0 | Yeast BUB homolog | -0.71 | 3.4E-02 |
| ACLY | Probable ATP-citrate synthase | -0.71 | 4.9E-05 |
| RL27 | 60S ribosomal protein L27 | -0.71 | 4.2E-04 |
| ASNA | ATPase asna-1 | -0.72 | 3.0E-04 |
| Q4JFH6 | SaPosin-like Protein family | -0.73 | 2.7E-02 |
| O61977 | TransThyretin-Related family domain | -0.73 | 2.3E-02 |

| | | | |
|---|---|---|---|
| Q23588 | Uridine PhosPhorylase | -0.73 | 3.1E-02 |
| RL7A | 60S ribosomal protein L7a | -0.74 | 1.6E-06 |
| IMPA1 | Inositol monophosphatase ttx-7 | -0.74 | 7.8E-05 |
| Q9TZL9 | Fatty Acyl-CoA ReDuctase | -0.75 | 2.6E-02 |
| TCPD | T-complex protein 1 subunit delta | -0.75 | 3.8E-10 |
| GMPR | GMP reductase | -0.75 | 3.9E-03 |
| ISW1 | Chromatin-remodeling complex ATPase chain isw-1 | -0.76 | 3.2E-02 |
| G5ECZ0 | Heavy chain, Unconventional Myosin | -0.76 | 4.7E-02 |
| B3WFV9 | Anion exchange protein | -0.76 | 7.2E-04 |
| EIF3A | Eukaryotic translation initiation factor 3 subunit A | -0.77 | 2.9E-12 |
| RSP2 | Probable splicing factor, arginine/serine-rich 2 | -0.77 | 1.6E-04 |
| Q23368 | Yeast SEC homolog | -0.77 | 5.0E-03 |
| UN104 | Kinesin-like protein unc-104 | -0.77 | 1.2E-05 |
| RL4 | 60S ribosomal protein L4 | -0.77 | 4.6E-19 |

| | | | |
|---|---|---|---|
| EIF3B | Eukaryotic translation initiation factor 3 subunit B | -0.77 | 5.2E-08 |
| O62415 | LYSozyme | -0.77 | 5.0E-03 |
| AKAP1 | KH domain-containing protein akap-1 | -0.79 | 4.1E-03 |
| Q9TZD9 | HAlF transporter (PGP related) | -0.79 | 5.8E-06 |
| RL8 | 60S ribosomal protein L8 | -0.79 | 2.5E-13 |
| LDH | L-lactate dehydrogenase | -0.80 | 8.5E-04 |
| PRS10 | Probable 26S protease regulatory subunit 10B | -0.80 | 2.7E-07 |
| NTF2 | Probable nuclear transport factor 2 | -0.80 | 4.9E-02 |
| P92005 | CathePsin Z | -0.80 | 8.6E-11 |
| UNC87 | Protein unc-87 | -0.81 | 1.3E-06 |
| TCPG | T-complex protein 1 subunit gamma | -0.81 | 3.2E-09 |
| EIF3C | Eukaryotic translation initiation factor 3 subunit C | -0.81 | 3.4E-04 |
| Q7YZW5 | VEMA (Mammalian ventral midline antigen) related | -0.81 | 4.3E-02 |
| PHO5 | Putative acid phosphatase 5 | -0.81 | 4.0E-03 |

| | | | |
|---|---|---|---|
| Q21750 | Acid Alpha Glucosidase Relate | -0.82 | 1.5E-05 |
| P91027 | Calponin Homology Domain containing Protein | -0.82 | 1.9E-05 |
| G5ECX8 | Increased Sodium Tolerance Related | -0.82 | 2.7E-02 |
| U2AF2 | Splicing factor U2AF 65 kDa subunit;U2AF splicing factor | -0.82 | 6.1E-03 |
| AKT1 | Serine/threonine-protein kinase akt-1 | -0.82 | 1.9E-02 |
| Q21801 | GAlactosidase/N-Acetylgalactosaminidase | -0.82 | 1.0E-05 |
| PLBL1 | Putative phospholipase B-like 1 | -0.82 | 7.4E-18 |
| OLA1 | Obg-like ATPase 1 | -0.83 | 6.0E-06 |
| RS21 | 40S ribosomal protein S21 | -0.83 | 8.9E-10 |
| P91277 | Heterogeneous nuclear RibonucleoProtein (HnRNP) K homolog | -0.83 | 1.1E-03 |
| CPLX1 | Putative complexin-1 | -0.84 | 2.8E-02 |
| RS12 | 40S ribosomal protein S12 | -0.85 | 6.5E-03 |
| Q18886 | Related to yeast Vacuolar Protein Sorting factor | -0.86 | 1.5E-03 |

| | | | |
|---|---|---|---|
| Q22170 | Intracellular LEctin | -0.86 | 2.5E-02 |
| SYDC | Aspartate--tRNA ligase, cytoplasmic | -0.86 | 9.0E-05 |
| Q22716 | Ribosomal Protein, Large subunit | -0.87 | 9.2E-03 |
| KS6A2 | Putative ribosomal protein S6 kinase alpha-2 | -0.88 | 1.9E-02 |
| CYP3 | Peptidyl-prolyl cis-trans isomerase 3 | -0.88 | 1.2E-02 |
| O44782 | Major sperm protein | -0.88 | 5.5E-03 |
| Q23382 | CCDC (Human Coiled Coil Domain Containing) homolog | -0.88 | 8.2E-07 |
| CAND1 | Cullin-associated NEDD8-dissociated protein 1 | -0.89 | 6.0E-05 |
| PHM | Probable peptidylglycine alpha-hydroxylating monooxygenase 1 | -0.90 | 1.3E-03 |
| PFD3 | Probable prefoldin subunit 3 | -0.90 | 1.1E-03 |
| O62334 | AGL (Amylo-1,6-GLucosidase, 4-alpha-glucanotransferase) glycogen debranching enzyme | -0.91 | 7.3E-04 |
| FBRL | rRNA 2'-O-methyltransferase fibrillarin | -0.91 | 2.4E-04 |
| Q20306 | ABC transporter, class F | -0.91 | 4.8E-03 |

| | | | |
|---|---|---|---|
| SDHF2 | Succinate dehydrogenase assembly factor 2, mitochondrial | -0.91 | 1.3E-02 |
| PROF2 | Profilin-2 | -0.91 | 2.9E-05 |
| G5EF26 | Putative low density lipoprotein receptor associated protein (37.4 kD) | -0.92 | 7.1E-03 |
| H2L0C0 | EMC Endoplasmic Membrane protein Complex (Yeast EMC) homolog | -0.93 | 4.0E-09 |
| PLBL2 | Putative phospholipase B-like 2 | -0.93 | 3.4E-13 |
| COPD | Probable coatomer subunit delta | -0.94 | 3.4E-03 |
| Q20277 | FIP (Fungus-Induced Protein) Related | -0.94 | 1.9E-03 |
| GYG1 | Glycogenin-1 | -0.94 | 1.4E-03 |
| SYSC | Probable serine--tRNA ligase, cytoplasmic | -0.95 | 6.0E-05 |
| DX39B | Spliceosome RNA helicase DDX39B homolog | -0.95 | 6.1E-07 |
| ELO6 | Elongation of very long chain fatty acids protein 6 | -0.96 | 4.7E-02 |
| SYNC | Asparagine--tRNA ligase, cytoplasmic | -0.96 | 1.4E-02 |
| Q86NH9 | TransThyretin-Related family domain | -0.96 | 2.7E-02 |

| | | | |
|---|---|---|---|
| Q9TYS8 | CALUmenin (Calcium-binding protein) homolog | -0.96 | 1.3E-03 |
| Q22240 | Ubiquitin carboxyl-terminal hydrolase | -0.96 | 1.8E-02 |
| KAD1 | Adenylate kinase isoenzyme 1 | -0.96 | 2.3E-03 |
| AT131 | Probable manganese-transporting ATPase catp-8 | -0.96 | 2.3E-08 |
| ADF1 | Actin-depolymerizing factor 1, isoforms a/b | -0.97 | 2.0E-02 |
| O44985 | Tumorous Enhancer of Glp-1(Gf) | -0.97 | 2.6E-08 |
| G8JY03 | DEAD boX helicase homolog;Pre-mRNA-splicing factor ATP-dependent RNA helicase ddx-15 | -0.97 | 7.7E-03 |
| A3QMC6 | ERLin (ER lipid raft associated protein) homolog | -0.97 | 4.5E-04 |
| Q21473 | AQuaPorin or aquaglyceroporin related | -0.98 | 2.2E-02 |
| RS7 | 40S ribosomal protein S7 | -0.98 | 3.1E-07 |
| G5EC40 | Trehalase | -0.98 | 4.0E-02 |
| MDHM | Probable malate dehydrogenase, mitochondrial | -0.99 | 1.8E-05 |

| | | | |
|---|---|---|---|
| SPCS2 | Probable signal peptidase complex subunit 2 | -0.99 | 5.4E-04 |
| Q21746 | Small Glutamine-rich Tetratrico repeat protein | -0.99 | 1.9E-02 |
| A5HU98 | GUanylate Kinase | -0.99 | 3.4E-05 |
| SC61G | Protein transport protein Sec61 subunit gamma | -1.00 | 5.0E-03 |
| Q22947 | RH (Rhesus) antigen Related | -1.00 | 4.6E-02 |
| IF4A | Eukaryotic initiation factor 4A | -1.00 | 1.1E-03 |
| LARP1 | La-related protein 1 | -1.00 | 6.2E-03 |
| Q20829 | Alpha-mannosidase | -1.00 | 7.0E-06 |
| PGK | Probable phosphoglycerate kinase | -1.01 | 3.1E-07 |
| EIF3L | Eukaryotic translation initiation factor 3 subunit L | -1.01 | 2.9E-04 |
| COPB2 | Probable coatomer subunit beta' | -1.01 | 3.7E-04 |
| G5EC23 | HCF1 related | -1.01 | 4.5E-04 |
| RS17 | 40S ribosomal protein S17 | -1.02 | 2.3E-09 |
| Q965V4 | EXPOrtin (Nuclear export receptor) | -1.02 | 2.6E-02 |

| | | | |
|---|---|---|---|
| PRP19 | Pre-mRNA-processing factor 19 | -1.02 | 9.8E-04 |
| YQ4B | Putative pseudouridine synthase B0024.11 | -1.03 | 3.5E-03 |
| RL7 | 60S ribosomal protein L7 | -1.03 | 4.1E-09 |
| RPF2 | Ribosome production factor 2 homolog | -1.04 | 1.7E-02 |
| Q9U1W8 | LSM Sm-like protein | -1.06 | 2.6E-02 |
| PFD4 | Probable prefoldin subunit 4 | -1.07 | 1.2E-02 |
| CNOT7 | CCR4-NOT transcription complex subunit 7 | -1.08 | 9.1E-04 |
| STDH1 | Putative steroid dehydrogenase 1 | -1.08 | 3.7E-02 |
| TORS | Torsin-like protein | -1.08 | 4.7E-04 |
| Q21832 | RNA-binding protein 8A | -1.09 | 1.4E-04 |
| SQV2 | Beta-1,3-galactosyltransferase sqv-2 | -1.09 | 2.5E-02 |
| ADRM1 | Proteasomal ubiquitin receptor ADRM1 homolog | -1.10 | 3.9E-04 |
| PPM1 | Protein phosphatase ppm-1 | -1.10 | 9.5E-03 |
| Q9N363 | mRNA DeCAPping enzyme | -1.10 | 1.1E-02 |

| | | | |
|---|---|---|---|
| RL10 | 60S ribosomal protein L10 | -1.11 | 1.8E-08 |
| RPAB3 | Probable DNA-directed RNA polymerases I, II, and III subunit RPABC3 | -1.11 | 7.4E-03 |
| LMP1 | LAMP family protein lmp-1 | -1.11 | 9.1E-05 |
| Q17796 | Hepatocyte Growth factor-Regulated TK Substrate (HRS) family | -1.11 | 4.2E-02 |
| PP2C2 | Probable protein phosphatase 2C T23F11.1 | -1.11 | 2.3E-02 |
| O61742 | Proteasome Regulatory Particle, Non-ATPase-like | -1.12 | 2.0E-02 |
| Q20107 | Peptidyl-prolyl cis-trans isomerase | -1.12 | 1.4E-02 |
| BTF3 | Transcription factor BTF3 homolog | -1.12 | 2.7E-05 |
| TBA3 | Tubulin alpha-3 chain | -1.13 | 6.7E-04 |
| O16482 | CYtochrome P450 family | -1.13 | 3.7E-03 |
| OTUBL | Ubiquitin thioesterase otubain-like | -1.13 | 2.2E-02 |
| O62289 | TransThyretin-Related family domain | -1.13 | 4.5E-02 |
| P90961 | Clathrin LIght Chain | -1.13 | 1.8E-02 |

| | | | |
|---|---|---|---|
| Q9N4N4 | SWI/SNF nucleosome remodeling complex component | -1.13 | 6.9E-04 |
| PAR5 | 14-3-3-like protein 1 | -1.14 | 2.2E-04 |
| Q27492 | DNA-directed RNA polymerase subunit beta | -1.14 | 4.2E-02 |
| RS5 | 40S ribosomal protein S5 | -1.15 | 3.6E-06 |
| MMSA | Probable methylmalonate-semialdehyde dehydrogenase [acylating], mitochondrial | -1.15 | 2.4E-03 |
| Q1XFY9 | Ribosomal Protein, Small subunit | -1.15 | 1.5E-08 |
| G5EFP5 | Alpha1,3-fucosyltransferase homologue | -1.16 | 7.3E-03 |
| Q9XWP7 | Eukaryotic Initiation Factor | -1.16 | 1.1E-02 |
| UBP14 | Ubiquitin carboxyl-terminal hydrolase 14 | -1.16 | 8.6E-04 |
| RL37A | 60S ribosomal protein L37a | -1.16 | 5.6E-03 |
| MTAP | S-methyl-5'-thioadenosine phosphorylase | -1.17 | 2.8E-02 |
| STT3 | Dolichyl-diphosphooligosaccharide--protein glycosyltransferase subunit stt-3 | -1.17 | 3.5E-03 |

| | | | |
|---|---|---|---|
| G5EGS9 | N-acetyllactosamine synthase | -1.19 | 5.7E-04 |
| Q9N3C9 | RNA Polymerase II (B) subunit | -1.20 | 2.3E-02 |
| Q9U2S6 | Peptidyl-prolyl cis-trans isomerase E | -1.20 | 1.3E-02 |
| TCPZ | T-complex protein 1 subunit zeta | -1.20 | 3.1E-03 |
| HSP7C | Heat shock 70 kDa protein C | -1.20 | 8.5E-09 |
| Q93878 | Yeast ERV (ER to Golgi transport Vesicle protein) homolog | -1.22 | 1.1E-02 |
| Q9XTT9 | Proteasome Regulatory Particle, ATPase-like | -1.22 | 7.6E-03 |
| LSM4 | Probable U6 snRNA-associated Sm-like protein LSm4 | -1.22 | 1.1E-02 |
| DMON2 | DOMON domain-containing protein Y73F4A.1 | -1.22 | 1.9E-05 |
| UNC84 | Nuclear migration and anchoring protein unc-84 | -1.23 | 9.5E-04 |
| Q21559 | Ref/ALY RNA export adaptor family | -1.23 | 7.3E-05 |
| CYP6 | Peptidyl-prolyl cis-trans isomerase 6 | -1.23 | 2.7E-06 |
| AAPK2 | 5'-AMP-activated protein kinase catalytic subunit alpha-2 | -1.25 | 8.4E-04 |

| | | | |
|---|---|---|---|
| H2KZJ9 | Galectin | -1.25 | 1.5E-02 |
| A5Z2S4 | SAPS (Phosphatase associated) domain protein | -1.25 | 2.2E-02 |
| Q21351 | Ras-Gtpase-activating protein SH3 (Three) domain-Binding Protein | -1.25 | 3.6E-08 |
| P91502 | Ubiquitin carboxyl-terminal hydrolase | -1.25 | 1.7E-04 |
| TECR | Probable very-long-chain enoyl-CoA reductase art-1 | -1.26 | 1.9E-09 |
| RCANL | Calcipressin-like protein | -1.26 | 2.2E-03 |
| Q27526 | Beta-galactosidase | -1.27 | 5.5E-03 |
| NACA | Nascent polypeptide-associated complex subunit alpha | -1.27 | 1.9E-02 |
| TPSTA | Protein-tyrosine sulfotransferase A | -1.27 | 1.6E-02 |
| IF5A2 | Eukaryotic translation initiation factor 5A-2 | -1.28 | 9.6E-03 |
| TCPB | T-complex protein 1 subunit beta | -1.28 | 1.2E-13 |
| ADR2 | Probable double-stranded RNA-specific adenosine deaminase | -1.28 | 7.4E-03 |

| | | | |
|---|---|---|---|
| RSP1 | Probable splicing factor, arginine/serine-rich 1 | -1.28 | 6.1E-05 |
| Q21831 | SNF chromatin remodeling Complex component | -1.32 | 3.6E-03 |
| H2KZJ5 | GLutaRedoXin | -1.33 | 2.9E-02 |
| EIF3H | Eukaryotic translation initiation factor 3 subunit H | -1.33 | 7.4E-06 |
| RUVB2 | RuvB-like 2 | -1.33 | 7.0E-05 |
| PSA2 | Proteasome subunit alpha type-2 | -1.34 | 1.3E-04 |
| Q9NLD1 | HnRNP A1 homolog | -1.34 | 1.2E-09 |
| GRP1 | GTP exchange factor for ARFs 1 | -1.35 | 2.5E-02 |
| Q20752 | STCH (Truncated HSP) family | -1.35 | 5.3E-05 |
| Q94207 | SUPpressor | -1.36 | 2.5E-02 |
| IF2B | Eukaryotic translation initiation factor 2 subunit 2 | -1.37 | 2.3E-03 |
| ALF2 | Fructose-bisphosphate aldolase 2 | -1.38 | 5.0E-02 |
| G5ECL3 | Pre-RNA processing 21 | -1.38 | 8.9E-03 |
| G5EE04 | Hsp-70 Interacting Protein homolog | -1.39 | 5.4E-04 |

| | | | |
|---|---|---|---|
| Q9U2F6 | Related to yeast Vacuolar Protein Sorting factor | -1.39 | 1.2E-02 |
| H2KYR1 | VIG (Drosophila Vasa Intronic Gene) ortholog | -1.39 | 2.3E-07 |
| ARL3 | ADP-ribosylation factor-like protein 3 | -1.39 | 5.1E-06 |
| G1K108 | Amino acid Transporter GlycoProtein subunit | -1.40 | 1.5E-04 |
| Q8IA75 | Phosphoinositide phospholipase C | -1.40 | 4.5E-02 |
| Q93565 | ToLLIp homolog | -1.40 | 2.1E-02 |
| HEXA | Beta-hexosaminidase A | -1.40 | 1.8E-09 |
| Q9XXA2 | Microtubule End Binding Protein | -1.40 | 4.1E-02 |
| O16720 | Protein argonaute | -1.42 | 1.9E-02 |
| O76371 | Proteasome Regulatory Particle, ATPase-like | -1.43 | 2.9E-05 |
| RPB2 | DNA-directed RNA polymerase II subunit RPB2 | -1.43 | 7.2E-05 |
| G5EBQ9 | SIPA (Vertebrate Signal-Induced Proliferation-Associated) homolog | -1.43 | 2.0E-03 |
| PDI2 | Protein disulfide-isomerase 2 | -1.43 | 2.2E-02 |

| | | | |
|---|---|---|---|
| O44729 | Yeast PRP (Splicing factor) related | -1.44 | 3.5E-03 |
| G5EES3 | Protein argonaute | -1.45 | 8.2E-09 |
| P90787 | alpha-1,2-Mannosidase | -1.47 | 1.6E-04 |
| H2KZK7 | Dipeptidyl Peptidase Four (IV) family | -1.47 | 1.0E-03 |
| RS23 | 40S ribosomal protein S23 | -1.47 | 1.5E-06 |
| ADPGK | Probable ADP-dependent glucokinase | -1.47 | 2.5E-07 |
| DAF31 | N-alpha-acetyltransferase daf-31 | -1.47 | 4.0E-02 |
| YQ83 | GYF domain-containing protein C18H9.3 | -1.47 | 1.8E-03 |
| Q9TXU7 | Eukaryotic Initiation Factor | -1.48 | 3.3E-02 |
| Q21152 | Fatty Acid/Retinol binding protein | -1.48 | 1.3E-02 |
| RL21 | 60S ribosomal protein L21 | -1.49 | 3.5E-06 |
| Q9U1V9 | DNaJ domain (Prokaryotic heat shock protein) | -1.49 | 6.6E-05 |
| O17406 | AT hook Transcription Factor family | -1.51 | 2.3E-07 |
| K8ERX1 | Zipcode Binding Protein homolog | -1.51 | 1.1E-02 |

| | | | |
|---|---|---|---|
| TMED2 | Suppressor/enhancer of lin-12 protein 9 | -1.53 | 2.7E-03 |
| A3QMC5 | Ribosomal Protein, Large subunit | -1.53 | 5.7E-06 |
| SRP19 | Probable signal recognition particle 19 kDa protein | -1.53 | 4.1E-03 |
| Q95002 | Serine/threonine-protein phosphatase | -1.53 | 1.3E-03 |
| LI15B | Protein lin-15B | -1.54 | 2.8E-07 |
| PFD1 | Probable prefoldin subunit 1 | -1.54 | 4.5E-02 |
| Q23463 | Elongation Factor TU family | -1.54 | 3.7E-10 |
| G5EDL7 | RILP (Rab7-Interacting Lysosomal Protein) homolog | -1.54 | 2.7E-02 |
| Q19440 | Phosphotransferase | -1.54 | 3.0E-04 |
| Q20684 | GaLECtin | -1.55 | 9.6E-05 |
| TOP2 | Probable DNA topoisomerase 2 | -1.56 | 5.7E-12 |
| Q21551 | Coiled coil Helix Coiled coiled Helix domain | -1.56 | 3.7E-02 |
| Q19162 | Ribosomal Protein, Large subunit | -1.56 | 4.3E-02 |
| RS19 | 40S ribosomal protein S19 | -1.57 | 8.0E-07 |

| | | | |
|---|---|---|---|
| GSLG1 | Golgi apparatus protein 1 homolog | -1.58 | 1.2E-11 |
| G5ED46 | CAlSYntenin/Alcadein homolog | -1.58 | 1.1E-02 |
| DAD1 | Dolichyl-diphosphooligosaccharide--protein glycosyltransferase subunit dad-1 | -1.59 | 4.2E-02 |
| Q9U1X6 | EMC Endoplasmic Membrane protein Complex (Yeast EMC) homolog | -1.59 | 1.1E-03 |
| Q22743 | RAs-related GTP-binding protein A | -1.59 | 2.5E-03 |
| O76630 | EMC Endoplasmic Membrane protein Complex (Yeast EMC) homolog | -1.59 | 1.3E-08 |
| Q9XVE9 | Ribosomal Protein, Large subunit | -1.62 | 3.6E-07 |
| LEM2 | LEM protein 2 | -1.62 | 9.3E-05 |
| G5ED01 | INOsitol-3-phosphate Synthase | -1.63 | 3.1E-11 |
| Q967F1 | Eukaryotic Initiation Factor | -1.64 | 2.7E-02 |
| Q9GUP2 | Enhancer of EfL-1 mutant phenotype | -1.64 | 3.0E-02 |
| SMD3 | Small nuclear ribonucleoprotein Sm D3 | -1.64 | 9.6E-06 |
| RUXG | Probable small nuclear ribonucleoprotein G | -1.65 | 1.6E-03 |

| | | | |
|---|---|---|---|
| RL36 | 60S ribosomal protein L36 | -1.65 | 7.6E-07 |
| HDA1 | Histone deacetylase 1 | -1.65 | 1.7E-06 |
| PRP8 | Pre-mRNA-splicing factor 8 homolog | -1.65 | 3.7E-13 |
| Q20774 | DNaJ domain (Prokaryotic heat shock protein) | -1.66 | 6.6E-03 |
| Q9XW17 | Cytokinesis, Apoptosis, RNA-associated | -1.66 | 6.8E-03 |
| WDR51 | WD repeat-containing protein wdr-5.1 | -1.68 | 1.3E-03 |
| YLC1 | ER membrane protein complex subunit 7 homolog | -1.68 | 6.1E-04 |
| G5EDQ2 | HMG | -1.68 | 1.1E-03 |
| G5EFS2 | MuSashI (Fly neural) family | -1.71 | 3.5E-04 |
| Q21295 | Nuclear Pore complex Protein | -1.72 | 6.8E-03 |
| Q9U2F5 | TIA-1/TIAL RNA binding protein homolog | -1.73 | 1.6E-02 |
| PLDL | Probable phospholipase D F09G2.8 | -1.75 | 8.8E-04 |
| VMP1 | Ectopic P granules protein 3 | -1.77 | 2.1E-02 |

| | | | |
|---|---|---|---|
| SRRT | Serrate RNA effector molecule homolog | -1.77 | 4.0E-04 |
| Q8WTJ4 | UBX-containing protein in Nematodes | -1.77 | 1.5E-02 |
| CYC21 | Cytochrome c 2.1 | -1.78 | 2.9E-06 |
| EIF3I | Eukaryotic translation initiation factor 3 subunit I | -1.81 | 2.5E-04 |
| SPT5H | Transcription elongation factor SPT5 | -1.82 | 2.4E-02 |
| NPL11 | Neprilysin-11 | -1.83 | 1.1E-06 |
| O17373 | Intestinal acid PHOsphatase | -1.84 | 2.5E-06 |
| G5EDT9 | UBR E3 ubiquitin ligase homolog | -1.84 | 6.9E-03 |
| RSP6 | Probable splicing factor, arginine/serine-rich 6 | -1.85 | 1.2E-03 |
| O17218 | Ribosomal Protein, Small subunit | -1.87 | 1.7E-05 |
| O62213 | C. Elegans Y-box | -1.87 | 4.3E-06 |
| G5EFE7 | Alpha-(1,6)-fucosyltransferase | -1.87 | 2.7E-03 |
| MGN | Protein mago nashi homolog | -1.87 | 5.6E-05 |
| PPT1 | Palmitoyl-protein thioesterase 1 | -1.88 | 6.4E-03 |

| | | | |
|---|---|---|---|
| CEC4 | Chromo domain-containing protein cec-4 | -1.89 | 1.4E-02 |
| SMC3 | Structural maintenance of chromosomes protein 3 | -1.92 | 2.5E-05 |
| Q21945 | Glutathione S-Transferase | -1.93 | 3.5E-03 |
| RS8 | 40S ribosomal protein S8 | -1.93 | 5.9E-09 |
| TACC1 | Transforming acid coiled-coil-containing protein 1 | -1.93 | 1.4E-02 |
| LMN1 | Lamin-1 | -1.94 | 4.5E-22 |
| MEC6 | Mechanosensory abnormality protein 6 | -1.94 | 1.4E-02 |
| Q9XTV4 | MBF (Multiprotein bridging factor) transcriptional coactivator | -1.95 | 3.4E-03 |
| RAB6B | Ras-related protein Rab-6.2 | -1.96 | 4.0E-04 |
| ROA1 | Heterogeneous nuclear ribonucleoprotein A1 | -1.96 | 1.6E-09 |
| PSB3 | Proteasome subunit beta type-3 | -1.96 | 1.7E-02 |
| H12 | Histone H1.2 | -1.96 | 2.1E-07 |

| | | | |
|---|---|---|---|
| RU2A | Probable U2 small nuclear ribonucleoprotein A' | -1.98 | 6.1E-06 |
| TNKS1 | Tankyrase-like protein | -1.98 | 4.2E-07 |
| CLAT | Choline O-acetyltransferase | -1.98 | 2.7E-10 |
| RL35 | 60S ribosomal protein L35 | -1.98 | 1.6E-05 |
| Q93641 | SAC1 PIP phosphatase (Yeast Suppressor of ACtin) homolog | -2.00 | 1.4E-03 |
| EF1B1 | Probable elongation factor 1-beta/1-delta 1 | -2.00 | 4.0E-02 |
| G5ED26 | MEL-46 | -2.03 | 1.8E-02 |
| O02325 | PRotein arginine MethylTransferase | -2.05 | 2.9E-02 |
| TCPQ | T-complex protein 1 subunit theta | -2.05 | 2.9E-03 |
| P90949 | Intestinal acid PHOsphatase | -2.06 | 8.3E-04 |
| RS27A | Ubiquitin-like protein 1-40S ribosomal protein S27a | -2.06 | 4.2E-04 |
| MEK2 | Dual specificity mitogen-activated protein kinase kinase mek-2 | -2.08 | 1.8E-02 |
| J7S164 | Histone H2A | -2.12 | 9.6E-09 |

| | | | |
|---|---|---|---|
| Q17832 | ViGiLN homolog | -2.13 | 6.6E-03 |
| Q17740 | ALG-1 INteracting protein | -2.13 | 2.2E-05 |
| Q19579 | Polyadenylate-binding protein | -2.13 | 4.3E-03 |
| H13 | Histone H1.3 | -2.15 | 1.5E-02 |
| SRP72 | Signal recognition particle subunit SRP72 | -2.15 | 1.3E-04 |
| C0KDV0 | K+/Cl-Cotransporter | -2.16 | 5.7E-03 |
| H11 | Histone H1.1 | -2.18 | 3.4E-10 |
| SMC1 | Structural maintenance of chromosomes protein 1 | -2.19 | 3.1E-04 |
| ATAT1 | Alpha-tubulin N-acetyltransferase 1 | -2.21 | 3.4E-04 |
| RL24 | 60S ribosomal protein L24 | -2.25 | 7.0E-08 |
| G5EFC4 | Cell Division Cycle related | -2.27 | 2.1E-02 |
| G5EDV3 | C. Elegans Y-box | -2.28 | 1.8E-04 |
| DAF7 | Dauer larva development regulatory growth factor daf-7 | -2.30 | 1.8E-04 |
| SCC3 | Cohesin subunit scc-3 | -2.30 | 2.6E-02 |

| | | | |
|---|---|---|---|
| Q9U2M1 | Condensin complex subunit 1 | -2.34 | 1.1E-02 |
| G5EFV4 | HMG | -2.34 | 5.2E-09 |
| CMK1 | Calcium/calmodulin-dependent protein kinase type 1 | -2.40 | 3.1E-02 |
| TPPP | Tubulin polymerization-promoting protein homolog | -2.42 | 2.5E-06 |
| TBB1 | Tubulin beta-1 chain | -2.46 | 1.7E-04 |
| SAE2 | SUMO-activating enzyme subunit uba-2 | -2.49 | 2.0E-02 |
| RS28 | 40S ribosomal protein S28 | -2.55 | 9.0E-07 |
| Q21793 | Prion-like-(Q/N-rich)-domain-bearing protein | -2.56 | 8.7E-05 |
| RS25 | 40S ribosomal protein S25 | -2.56 | 2.0E-05 |
| UFSP | Ufm1-specific protease | -2.59 | 4.5E-09 |
| RL22 | 60S ribosomal protein L22 | -2.64 | 4.1E-07 |
| CDC37 | Probable Hsp90 co-chaperone cdc37;Cell Division Cycle related | -2.66 | 1.6E-02 |
| RL36A | Ribosomal protein L36.A | -2.70 | 2.9E-03 |

| | | | |
|---|---|---|---|
| H2AV | Histone H2A.V | -2.73 | 2.8E-04 |
| Q8WQA8 | Ribosomal Protein, Small subunit | -2.73 | 7.6E-06 |
| RS6 | 40S ribosomal protein S6 | -2.75 | 2.3E-05 |
| G5EEL9 | HMG | -2.79 | 2.4E-05 |
| Q22622 | Major sperm protein | -2.84 | 3.3E-03 |
| Q9XWK3 | Nuclear Pore complex Protein | -2.90 | 6.3E-03 |
| RUVB1 | RuvB-like 1 | -2.94 | 1.9E-04 |
| AN321 | Acidic leucine-rich nuclear phosphoprotein 32-related protein 1 | -2.95 | 2.4E-02 |
| Q95QQ1 | DNaJ domain (Prokaryotic heat shock protein) | -3.00 | 3.7E-02 |
| Q17561 | Ref/ALY RNA export adaptor family | -3.02 | 4.1E-04 |
| Q9BIB7 | HnRNP F homolog | -3.10 | 7.4E-03 |
| RSP3 | Probable splicing factor, arginine/serine-rich 3 | -3.13 | 3.8E-09 |
| PP2C1 | Probable protein phosphatase 2C F42G9.1;Uncharacterized protein | -3.14 | 4.0E-03 |
| H4 | Histone H4 | -3.36 | 9.7E-19 |

| | | | |
|---|---|---|---|
| TOP1 | DNA topoisomerase 1 | -3.42 | 5.5E-05 |
| Q9UAV3 | Ubiquitin carboxyl-terminal hydrolase | -3.49 | 4.1E-04 |
| RIC3 | Resistance to inhibitors of cholinesterase protein 3 | -3.66 | 5.2E-04 |
| RL28 | 60S ribosomal protein L28 | -3.76 | 5.0E-09 |
| CNEP1 | CTD nuclear envelope phosphatase 1 homolog | -3.89 | 2.9E-02 |
| TM2D4 | TM2 domain-containing protein ZK858.5 | -3.96 | 1.1E-02 |
